# Systematic evaluation using SigMatch reveals how sample tissues shape transcriptomic disease relationships

**DOI:** 10.64898/2026.11.17.766813

**Authors:** Kewalin Samart, Landon Buskirk, Abhirupa Ghosh, Arjun Krishnan, Janani Ravi

## Abstract

**Motivation:** Diseases often share underlying molecular characteristics captured by transcriptional responses. Leveraging this knowledge transfer can aid in uncovering novel disease biology and drug repurposing. Large-scale transcriptomic datasets and disease signatures provide an opportunity to computationally identify these shared disease relationships based on shared genome-wide gene expression changes. However, disease signatures are strongly influenced by sample tissue, making it difficult to distinguish shared disease biology from shared tissue origin. Despite this challenge, existing computational approaches do not systematically take into account or evaluate the impact of tissue on their predictions.

**Results:** We present SigMatch, a systematic evaluation of the impact of sample tissue on transcriptomic-based disease relationship prioritization. We evaluate this impact on existing correlation- and rank-based methods, and on an alternative family called reconstruction-based methods. In particular, we examine (i) mechanistic disease relationships given the context of sample tissues and (ii) prediction agreement with external evidence of disease relatedness based on shared genes, symptoms, drugs, and comorbidities. We found that predicted disease relationships by all the methods are heavily influenced by sample tissue. Tissue-specific genes are not the sole driver of the prediction scores. Across methods, removal of tissue-specific genes did not reduce the observed tissue-dependent patterns or improve recovery of evidence-supported disease pairs. This work demonstrates substantial sample-tissue influence in transcriptomic similarities between diseases and that sample tissue should be considered when identifying and interpreting inferred transcriptomic disease relationships.

## 1. Introduction

Transcriptomic data provide a rich resource for capturing disease responses, commonly in the form of gene signatures obtained from differential expression analysis comparing disease samples to respective control samples. These transcriptomic signatures can be compared across diseases to reveal shared biological characteristics, providing opportunities for knowledge transfer across diseases such as drug repurposing. Current methods infer such transcriptomic disease relationships via gene overlap [1–4], pathway overlap [1], network-based overlaps [5], overall rank correlation [6], or rank-biased similarity [7]. For instance, previous studies have reported similarities in transcriptomic signatures of tuberculosis (TB) and rheumatoid arthritis (RA), revealing shared molecular pathways including T-cell receptor signaling pathway and Toll-like receptor signaling pathway, which motivated the investigation of common therapeutic targets [8].

However, differential expression may not only capture disease effects, but also effects from interactions between the disease and sample tissue. The sample tissue could either be a clinically accessible tissue like whole blood or the tissue where the disease manifests (*i.e.*, its “mechanistic tissue,” *e.g.*, brain tissue in the case of Alzheimer’s disease). Either way, distinguishing between disease signature similarity arising from shared disease biology and that arising from sample tissue remains challenging. To our knowledge, all current approaches only evaluate the overall similarity between disease signatures and do not evaluate the contribution of sample tissue to the predicted relationships. Additionally, these methods mainly evaluate the identified disease relationships based on whether they are from the same (or different) disease systems, used as a proxy for disease relatedness. However, disease systems are confounded by tissue and do not capture non-obvious relationships across systems with potential therapeutic transfer opportunities, such as the mechanistic connection between TB and RA.

In this study, we develop SigMatch to evaluate the impact of sample tissue on prioritizing disease relationships using similarity of transcriptomic signatures. While current similarity methods include those using gene overlap [1–4], pathway overlap [1], and network-based overlap [5], they often depend on additional analytical choices such as significance thresholds, enrichment procedures, or network construction parameters. Therefore, we focus on Spearman [6] and SimSIP [7], methods that use entire disease signatures, treating them as continuous variables across the whole transcriptome, to capture shared characteristics. Another approach for using entire signatures to define disease relationships is to reconstruct a target disease signature using multiple other disease signatures using regression, within a statistical/machine learning framework. This reconstruction-based approach has shown promise in uncovering biological relationships between diseases based on sample-level absolute gene expression [9]. In contrast to Spearman and SimSIP, which calculate similarities between a target disease and every other disease independently, the reconstruction-based approach learns relationships competitively (jointly) through internal feature selection. Here, we implement the reconstruction-based approach using both linear and tree-based models: Lasso, ElasticNet, Gradient Boosting Machine (GBM), and Random Forest (RF). Below, we present a systematic evaluation of predicted disease relationships from all these methods against independent sources of disease relationships derived from shared drugs, symptoms, genetic information, and comorbidity, stratified by sample source tissue and disease system, and an analysis of the impact of tissue-specific genes on these results. Together, these results reveal several insights into sample tissue impact on identification of transcriptomically similar diseases.

## 2. Materials and methods

### 2.1 Disease transcriptomic data and preprocessing

We began with human RNAseq datasets from DiSignAtlas [4], a curated resource of disease differential signatures with the unified gene space of 19,910 common measured genes across all the signatures. Each signature was associated with a disease label and, when available, a tissue label. We standardized disease identifiers by mapping Unified Medical Language System (UMLS) [10] provided by DiSignAtlas to Monarch Disease Ontology (MONDO) [11] disease identifiers; signatures without MONDO identifiers were dropped (**Figure 1a**). We used Claude Sonnet 4.6 to assign disease system categories based on Disease Ontology (DO) anatomical entities (**Figure 1c)** [12] and standardized GTEx v11 [13] tissue labels, followed by careful manual, blinded validation (**Figure 1d**). Six diseases known to affect more than one organ system were assigned the “multi-organ system” label: adenocarcinoma, metabolic diseases, bacterial infectious disease, Gram-negative bacterial infections, metastatic carcinoma, and inflammatory disease.

**Figure 1.**
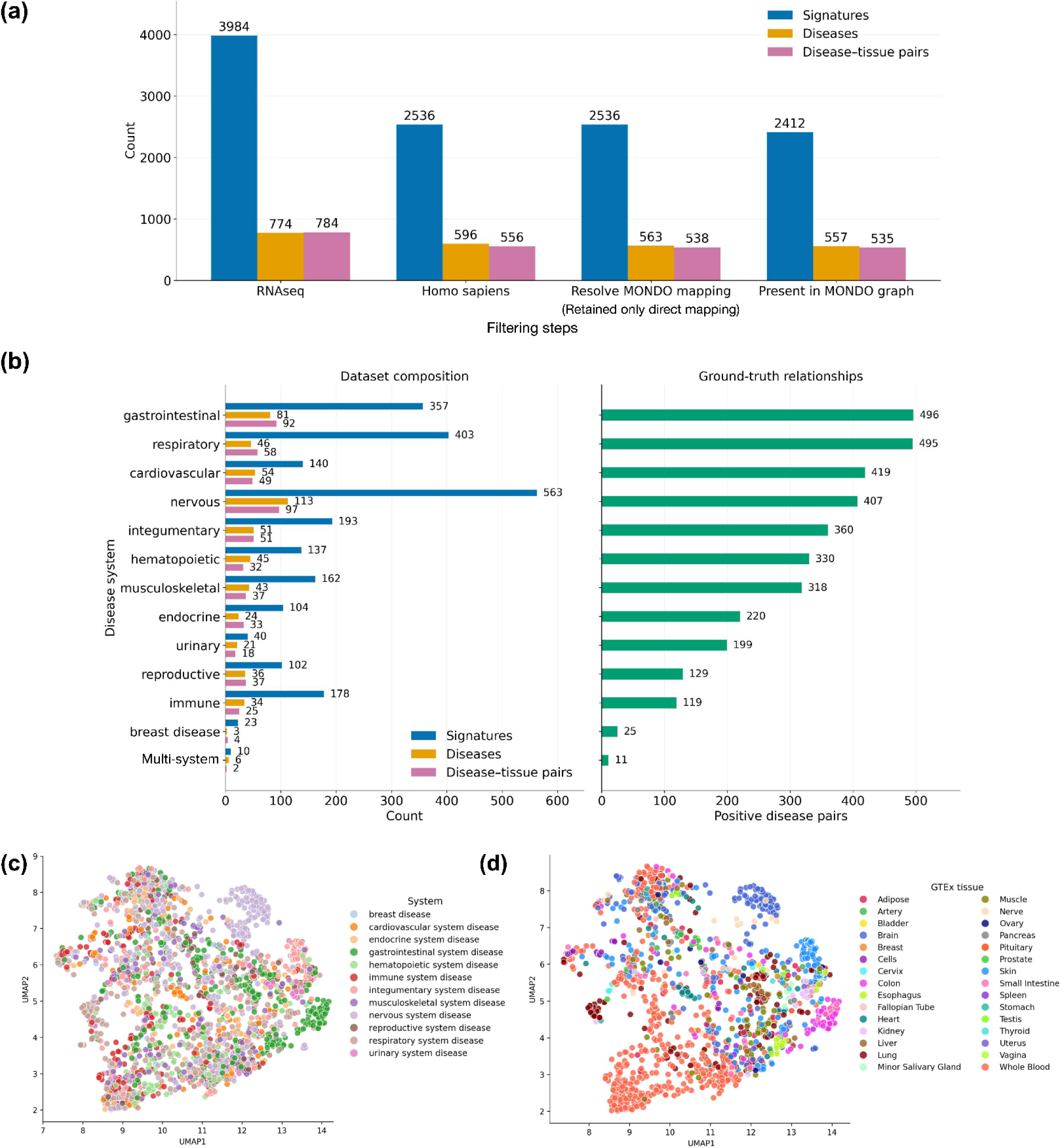
Data underlying SigMatch evaluation. **(a)** Numbers of disease signatures, diseases, and disease_tissue pairs leveraged by SigMatch at each filtering step. **(b)** Disease signature composition by affected organ system (referred to as disease system), with numbers of signatures, unique diseases, and disease_tissue pairs (left) and evidence-supported disease pairs count by system (right). Overall, SigMatch is powered by 2,412 disease signatures spanning 557 diseases with 535 disease_tissue pairs. **(c)** Uniform Manifold Approximation and Projection (UMAP) low-dimensional projection of all disease signatures colored by 12 disease systems (multi-system diseases were excluded) and **(d)** 30 generalized GTEx tissues.

### 2.2 Transcriptomic disease relationship prediction methods

We evaluated similarity between diseases by applying two approaches: (i) assessing similarity between disease signature pairs (**Figure 2a**) and (ii) reconstructing a disease signature from all other available disease signatures (**Figure 2b**). We then assessed reconstruction accuracy, biological relevance, and the effects of tissue context on predicted disease relationships (**Figure 2e**).

**Figure 2.**
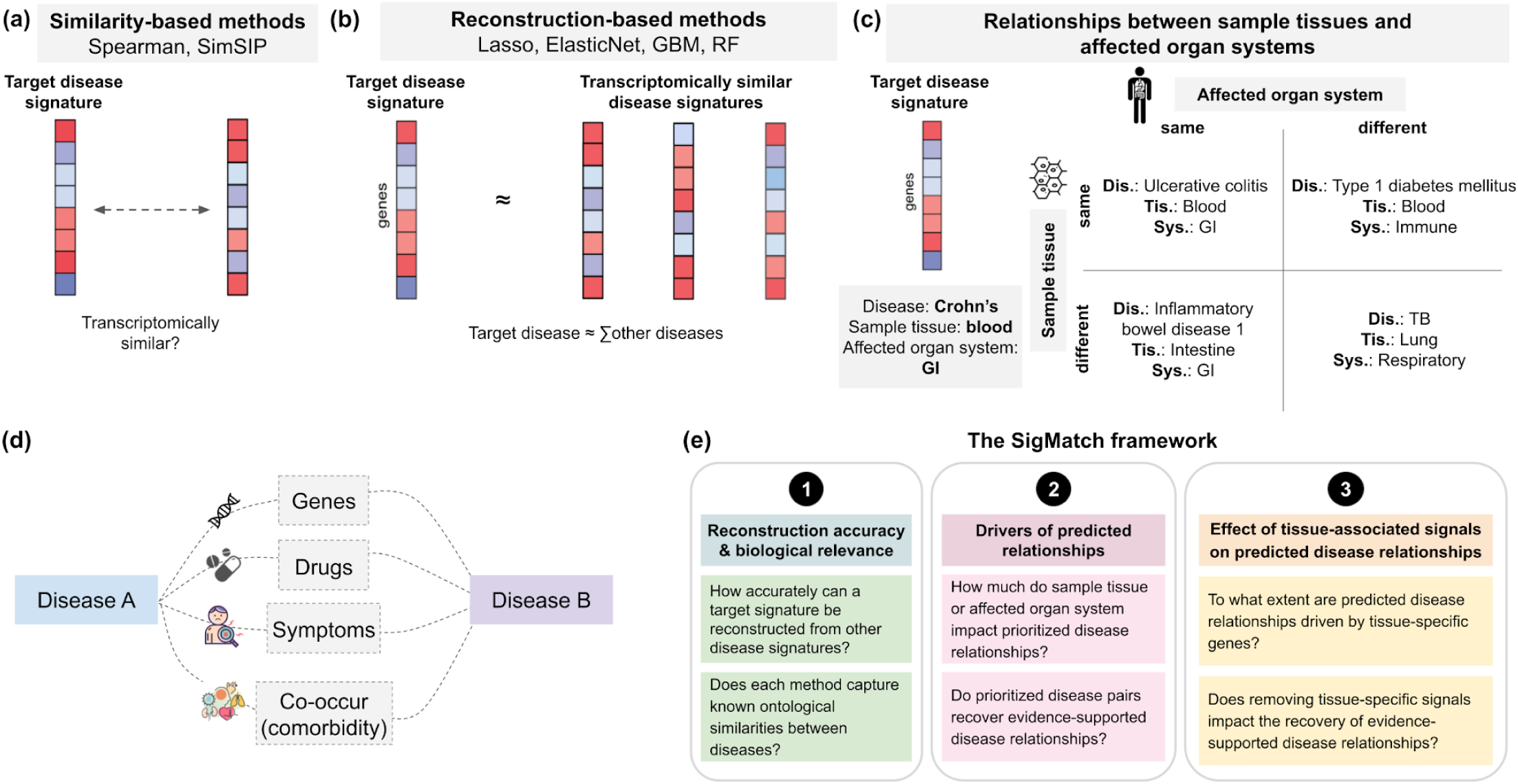
SigMatch evaluation scheme: methods used and key questions. **(a)** Similarity-based methods measuring global disease similarity of two disease signatures. **(b)** Reconstruction-based methods identifying transcriptomic disease similarity based on feature selection. **(c)** Four quadrants representing characteristics of predicted disease relationships used in SigMatch: same or different affected organ system/sample tissue with respect to the target disease. Crohn’s disease profiled in blood is depicted as an example target signature. **(d)** Schematic illustration of the ground-truth disease relationships used for SigMatch evaluation. Positive disease pairs are defined as those supported by at least two of four independent evidence sources: shared genes, shared drugs, shared symptoms, and disease co-occurrence (comorbidity). **(e)** The SigMatch framework is centered on (i) reconstruction accuracy and biological relevance based on ontology-related diseases, (ii) drivers of predicted relationships: how predictions are driven by the same sample tissue and/or mechanistic system, and (iii) effect of tissue-specific signals absence on predictions and evidence-supported disease relationship recovery.

#### 2.2.1 Similarity-based methods

1. <u>Spearman rank correlation</u> [14] (**Figure 2a**) measures similarity between diseases by assessing the correlation of differential gene expression across shared genes. Positive and negative correlations indicate similar and opposing transcriptional patterns, respectively. Spearman correlation among disease transcriptomic signatures revealed molecular similarities underlying co-occurring diseases [6]. In this study, we calculated Spearman to assess overall agreement in transcriptional changes between a pair of disease signatures. We considered only positive Spearman values as indicators of similarity, as we focused on identifying diseases with similar transcriptional patterns.
2. <u>SimSIP</u> (similarity score based on gene ranks; **Figure 2a**; [7]) quantifies disease similarity using reciprocal square-root weighting of gene ranks, assigning greater importance to genes strongly associated with both diseases. Unlike simple correlation methods, SimSIP emphasizes highly ranked disease-associated genes without requiring arbitrary gene-selection thresholds [7]. We computed SimSIP by ranking genes according to the absolute magnitude of their log_2_FC within each disease signature, assigning higher ranks to genes with larger expression changes regardless of direction.

#### 2.2.2 Reconstruction-based methods, model setups, and hyperparameter tuning

Reconstruction-based methods (**Figure 2b**) predict a “target” disease signature from “feature” signatures of other diseases using regression-based machine learning models. The data were organized as a gene-by-disease-signature matrix, with genes as observations, the target disease signature as the response variable, and all other disease signatures as predictors.

All reconstruction-based models were trained and evaluated using three-fold cross-validation with stratification based on gene regulation status (up-, down-, or non-regulated) per seed and were repeated across five random seeds. Lasso, ElasticNet, and RF regression models were implemented using the Python library scikit-learn (version 1.6.1) [15]. For the Gradient Boosting Machine, LightGBM (version 4.6.0) [16] was implemented with base learners constrained to decision stumps to enhance interpretability [17]. Remaining key hyperparameters were optimized using nested cross-validation on a few representative target signatures.

As previous pan-cancer transcriptomic analyses have identified shared transcriptional programs across cancer types, including those associated with cell-cycle regulation and immune responses [18], we assessed the influence of cancer-associated signals within our mixed pool of cancer and non-cancer signatures by projecting all the signatures onto a low-dimensional space using Uniform Manifold Approximation and Projection (UMAP [19]; **Figure S1**). As no clear clustering patterns distinguishing cancer from non-cancer signatures were observed, we applied a combined setting including both signature types for hyperparameter tuning (**Figure S2**). We performed cluster-based hyperparameter selection by first grouping targets via *k*-means (with number of clusters where *n* is the number of features). We formed a set of representative targets by picking the closest target to the centroid of each cluster. Next, we tuned the hyperparameters on each representative target and used the median of each hyperparameter across targets as the optimal hyperparameter configuration. By selecting one target per cluster and aggregating hyperparameters, this approach ensured that tuning captured variation across the target space while preventing a final configuration from being disproportionately driven by overrepresented signature profiles. The configuration was applied to remaining targets to evaluate model training and testing (**Methods S1.1; Figure S2**). Details of the hyperparameter tuning can be found in **Table S1**.

#### 2.2.3 Random gene-holdout evaluation

We evaluated the ability of reconstruction-based methods to capture transcriptional signals across disease signatures by performing leave-target-disease-out experiments. We trained one model per target signature (genes x 1) where all other signatures of the target disease were removed from the feature matrix (genes x signatures). To establish holdout-baseline-reconstruction performance for comparison with the tissue-gene holdout experiments described in **Methods 2.8**, we used a random-gene holdout strategy where the number of held-out genes matched the number of tissue-specific genes associated with the target’s sample tissue (**Figure S3**). The goal was to determine whether the models outperform their corresponding shuffled baselines (**Methods 2.2.4; Figure 3a**). For each target, we randomly sampled an equally sized set of genes from the full gene set of 19,910 genes to serve as the independent test set, and the remaining genes were used for model training and validation using three-fold cross-validation as described in **Methods 2.2.5**.

**Figure 3.**
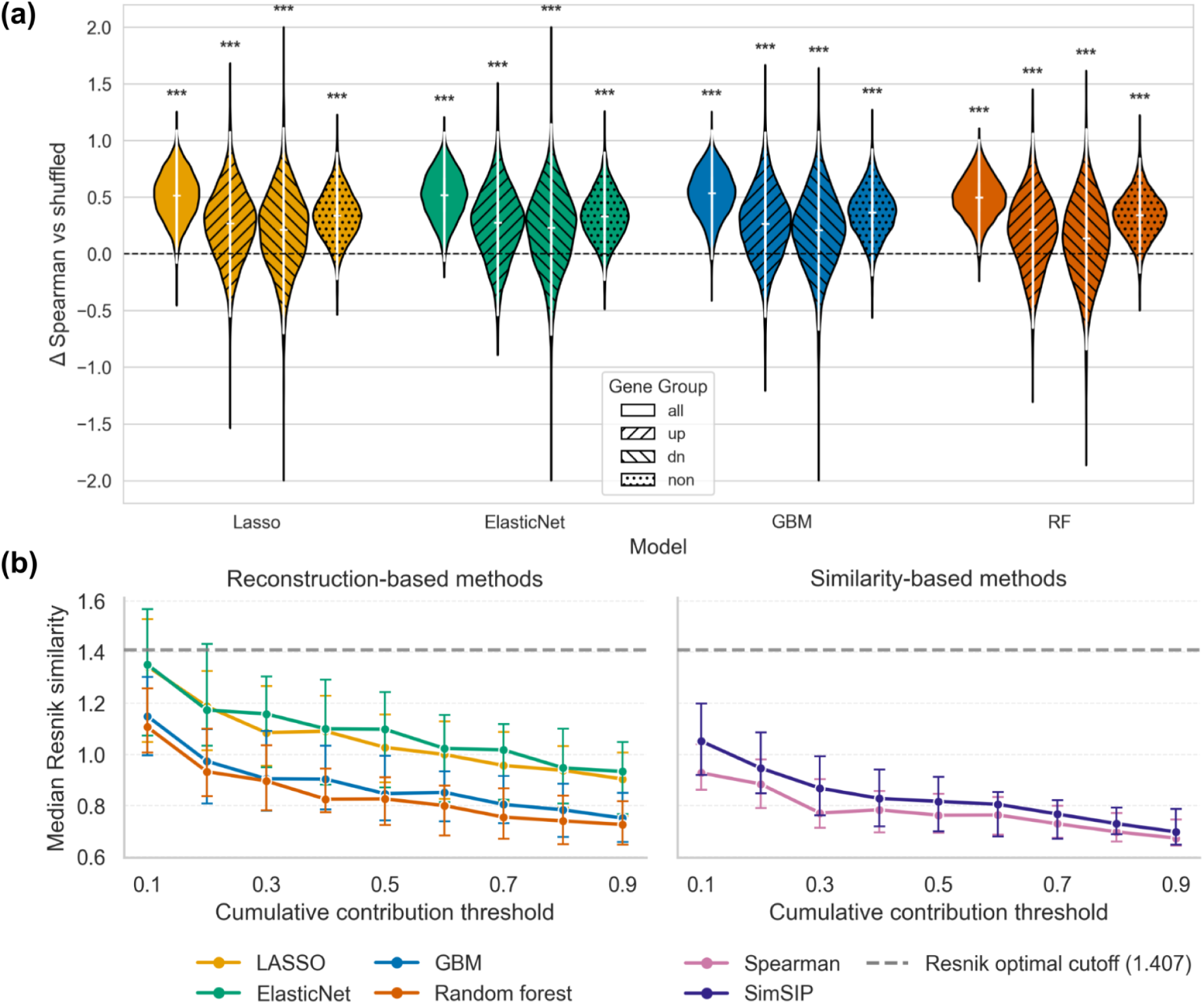
Reconstruction-based methods predict disease signatures beyond random expectation, and all methods recover ontology-related disease relationships. **(a)** Distribution of the improvement in reconstructed Spearman relative to shuffled controls (Δ Spearman vs. shuffled) for reconstruction-based models across four gene groups (all, upregulated, downregulated, and non-differentially expressed genes). Positive values indicate better predictive performance than shuffled controls. Negative values indicate worse predictive performance than shuffled controls. A zero indicates no improvement. *** indicates p-value < 0.001 (right-tailed paired Wilcoxon signed-rank test). **(b)** Median of mean Resnik similarity, based on MONDO, among disease relationships included up to each cumulative contribution threshold (0.1, 0.2, …, 0.9). For each target disease, disease relationships were ranked by their contribution to its prediction and included until their cumulative contribution reached the specified threshold. Higher median Resnik similarity indicates that prioritized disease pairs are more closely related within the ontology based on their most informative common ancestor. A Resnik similarity of zero indicates that the disease pair shares no informative common ancestor in the ontology. The optimal Resnik cutoff (∼1.4) indicates the Resnik score that best separates the same-system diseases from those from different systems (**Methods S1.3, Figures S6, S7**).

#### 2.2.4 Shuffled baseline models

For each target signature, we created permuted versions (baselines) by shuffling gene values (log fold change; log_2_FC) within the disease signature. SigMatch and the baseline models were then trained using the optimized hyperparameters in **Table S1** and evaluated on these permuted targets to estimate the random baseline performance of each model.

#### 2.2.5 Model performance evaluation

We evaluated model performance of all the holdout setups (**Methods 2.2.3, 2.8**) using three-fold cross-validation stratified by gene regulation status, with approximately 66.67% of the genes remaining after excluding the holdout-specific test set used for training and 33.33% for validation in each fold. We defined gene regulation groups based on the log_2_FC values of genes in each target disease signature. We classified genes as “upregulated” if their log_2_FC values were at or above the 80th percentile, “downregulated” if their values were at or below the negative of the 80th-percentile value, and “non-regulated” otherwise. We selected these thresholds to maintain consistent gene group sizes across disease signatures. Cross-validation performance was averaged across the three folds, and final model performance was evaluated separately on held-out test genes. We measured model performance using Spearman correlation between logFC values in the target and predicted signatures (*i.e.*, reconstructed Spearman; **Figure 3a**). Higher reconstructed Spearman indicates better reconstruction performance. For further biological interpretability, we also reported reconstructed Spearman stratified by gene regulation status (upregulated, downregulated, and non-regulated genes).

#### 2.2.6 Feature score aggregation

The feature importance scores were estimated during model training within each 3-fold cross-validation run and aggregated across folds using the mean feature importance. We ran each model with five different seeds to obtain estimated feature importance given expected variability in seed setting. To summarize aggregated scores across random seeds, we computed the median feature importance scores across five runs. We summarized features differently within and across targets depending on specific questions and analyses (the details can be found in **Table S2**).

### 2.3 Rank-based cumulative contribution of features to target

For each target signature, we ranked feature signatures according to their importance scores, with lower ranks indicating greater importance and average ranks assigned to ties. We then collapsed feature signatures to the disease_tissue level by calculating their median rank. To account for differences in the number of eligible feature signatures across targets, these median ranks were converted into normalized percentile ranks within each target signature. Across target signatures within a target disease_tissue, we calculated the median normalized percentile ranks for each feature disease_tissue and reranked these values to obtain the final rank for each target-feature pair of disease_tissue, with a lower rank indicating greater importance. For each target disease_tissue, the rank-based contribution score of feature disease_tissue (i) was defined as rank-based contribution using linear weighting to ensure a gradual decline in contribution between adjacent ranks: 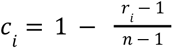 and we then calculated normalized rank-based contribution: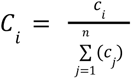, where *r* is the final rank obtained by reranking the median percentile ranks of feature disease_tissue signature, and n is the total number of candidate disease_tissue signatures with positive importance for that target disease_tissue so that 1 ≤ *i*, *j* ≤ *n*. In the absence of tied ranks, the highest-ranked candidate *r _i_*= 1 receives a rank-based contribution of 1, whereas the lowest-ranked candidate *r_i_* = n receives a contribution of 0. When ties occur, average ranks are assigned, and the corresponding weights are calculated using the same linear weighting formula.

Given differences in the distributions and scales of feature importance scores across target signatures, we used rank-based contributions to facilitate comparisons without relying on raw importance magnitudes. For each target disease_tissue pair, candidate feature disease_tissue pairs were ordered by their final ranks, from highest to lowest importance. The cumulative rank-based contribution of the top *k* candidates was calculated as: 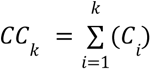 where *C_i_* is the normalized rank-based contribution of the candidate at rank *i*, and *k* is the number of highest-ranked candidates included. We applied cumulative rank-based contribution thresholds to identify the minimum set of highest-ranked feature disease_tissue pairs whose combined normalized rank-based contributions reached or exceeded a specified fraction of the total: *k_τ_* = *min*{*k*,*CC*(*k*)≥*τ*} where τ denotes the cumulative contribution threshold. We evaluated thresholds of τ = 0. 1, 0. 2, 0. 3, …, 0. 9. For instance, a threshold of 0.1 represents the set of highest-ranked features (disease_tissue signatures) that cumulatively account for 10% of the total contribution for a target.

While we described cumulative contribution concepts and calculation here using disease_tissue units, which were primarily used for downstream evaluation of sample tissue effects (**Methods 2.5, 2.9; Figures 4**, **5**, **6**, **7; Results 3.3–3.5, 3.7**), we instead used disease-disease units for all of our disease-level analyses (**Methods 2.4, 2.10; Figures 3b, 9b,c; Results 3.3, 3.6, 3.8**).

**Figure 4.**
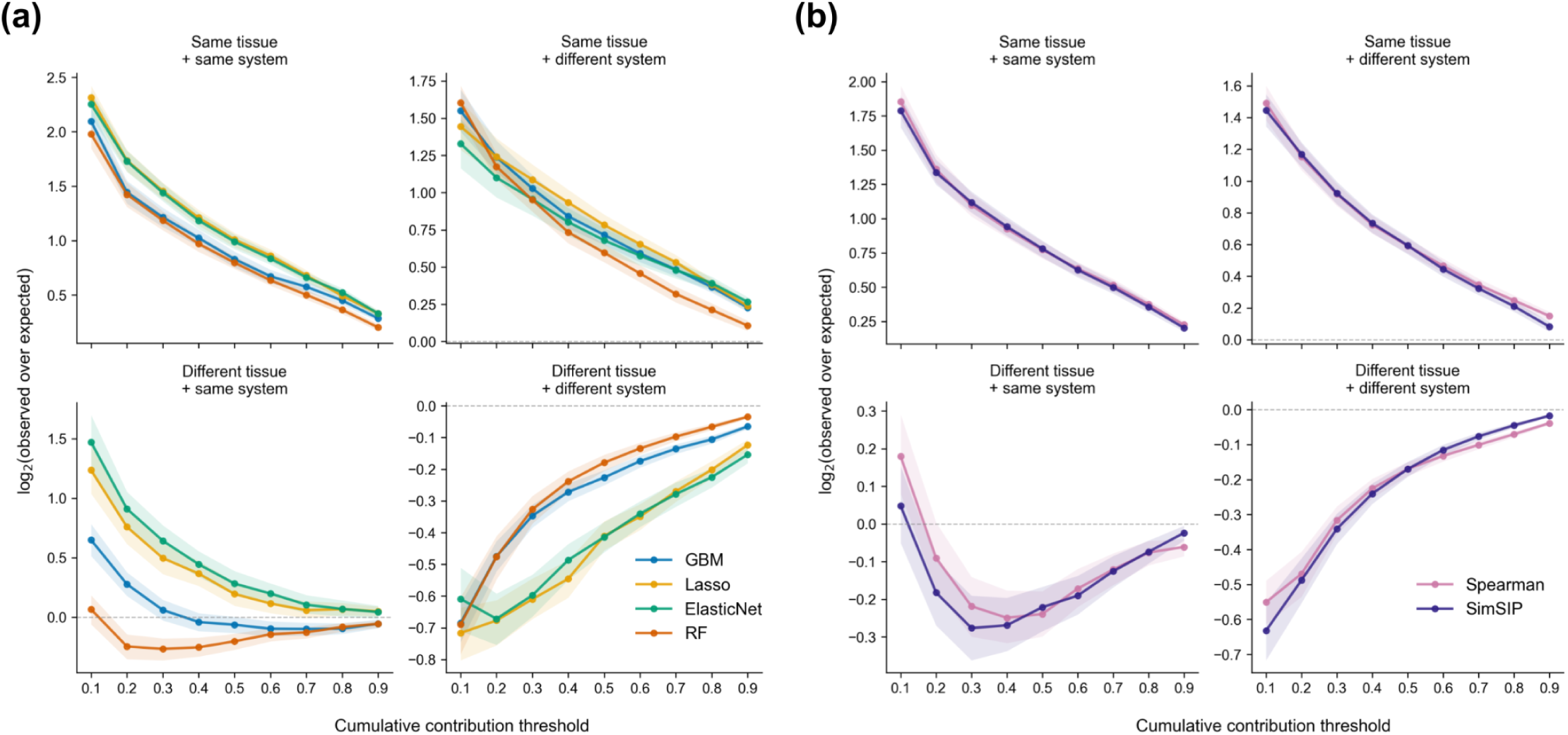
Disease signatures from the same sample tissue dominate the predicted disease relationships. Enrichment of prioritized disease signatures in tissue/system relationship categories at cumulative contribution thresholds (0.1, 0.2, …, 0.9) for **(a)** reconstruction- and **(b)** similarity-based methods. The y-axis shows log2(mean fold enrichment), where fold enrichment is the proportion of prioritized disease signatures in each tissue/system relationship category divided by the corresponding proportion among all candidate disease signatures. Positive values indicate overrepresentation, zero indicates no enrichment, and negative values indicate underrepresentation relative to background expectations (**Methods 2.5**).

**Figure 5.**
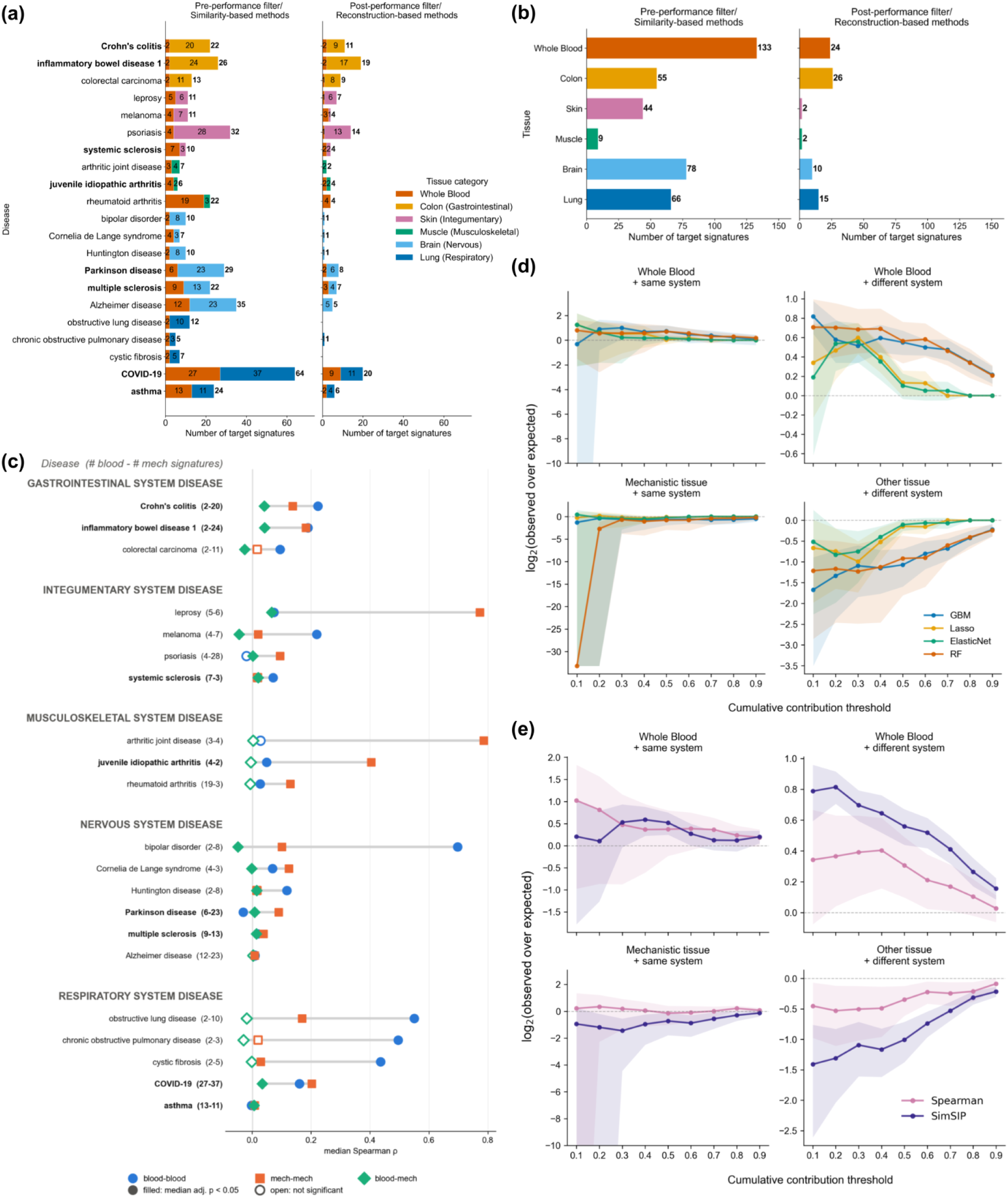
Reduced variation in dataset for biological interpretation and within-disease signature agreement and case-study selection. **(a)** Composition of the reduced dataset comprising 21 diseases, each with ≥2 whole-blood and ≥2 disease-associated mechanistic-tissue signatures. Shown are targets shared by both similarity-based methods before filtering (left) and the union of unique targets across reconstruction-based methods after filtering for reconstruction Spearman correlation >0.1 for both up- and downregulated genes (right). The diseases that pass the performance thresholds and have at least two whole-blood and two mechanistic-tissue signatures are bolded. **(b)** Total target signature counts by tissue for similarity-based methods equivalent to before reconstruction performance filtering (left) and the union of unique target signatures across reconstruction-based methods after filtering (right; eight bolded diseases only). **(c)** Within-disease concordance of signature pairs by tissue pairing, scored on all genes with Spearman ρ. Each row is a disease, grouped by disease system. Diseases meeting the reconstruction performance thresholds and having at least two whole-blood and two mechanistic-tissue signatures are indicated in bold, as in (a). The numbers in parentheses give the count of whole-blood and mechanistic-tissue signatures. Markers show the median score across within-disease pairs for blood-blood (circle), mech-mech (square) and blood-mech (diamond) pairs; grey bars span the range of the three medians. Filled markers indicate a median Bonferroni-adjusted p-value *P* < 0.05; open markers are not significant. Spearman: the vertical line marks ρ = 0. Enrichment of whole-blood or mechanistic tissue/system relationship categories among the disease relationships cumulatively contributes to different contribution thresholds for each. **(d)** reconstruction-based method and **(e)** similarity-based method: 0.1, 0.2, …, 0.9 given whole-blood target disease signatures. The y-axis shows log2(mean fold enrichment), where fold enrichment is the proportion of prioritized disease signatures in each tissue/system relationship category divided by the corresponding proportion among all candidate disease signatures. Positive values indicate overrepresentation, zero indicates no enrichment, and negative values indicate underrepresentation relative to background expectations (**Methods 2.5**).

**Figure 6.**
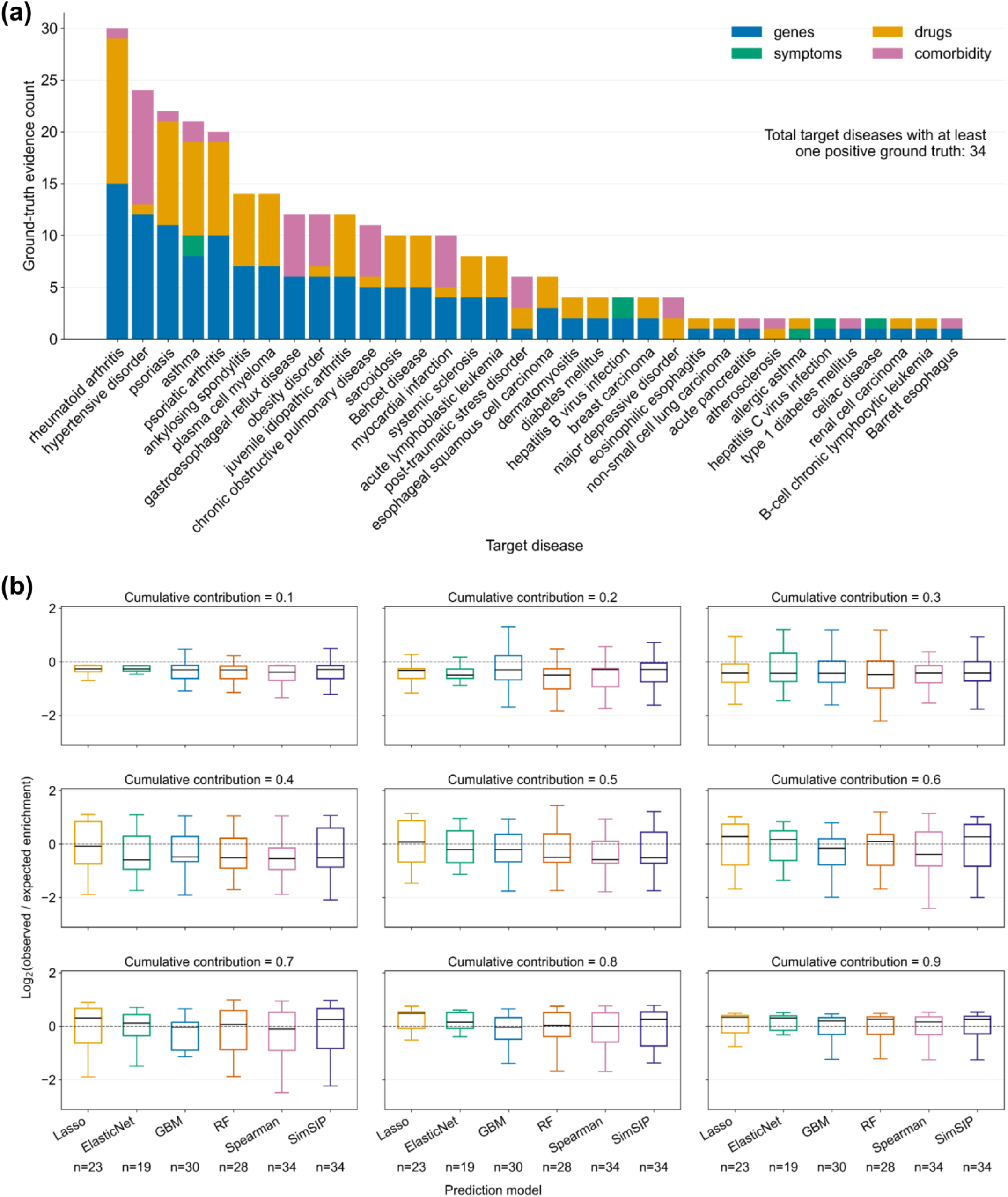
Ground-truth evidence and enrichment of predicted disease relationships under random gene holdout. **(a)** Distribution of four evidence types across 34 target diseases with at least one ground-truth-positive relationship, defined by support from at least two evidence types. **(b)** Log₂ observed compared with expected enrichment across six prediction methods at cumulative contribution thresholds of 0.1–0.9. Positive values indicate that random-holdout (**Methods 2.2.3**) predictions recover more evidence-supported disease relationships than expected under permutation, whereas negative values indicate fewer recoveries than expected. The dashed line indicates enrichment of zero, and n denotes the number of target diseases per method.

**Figure 7.**
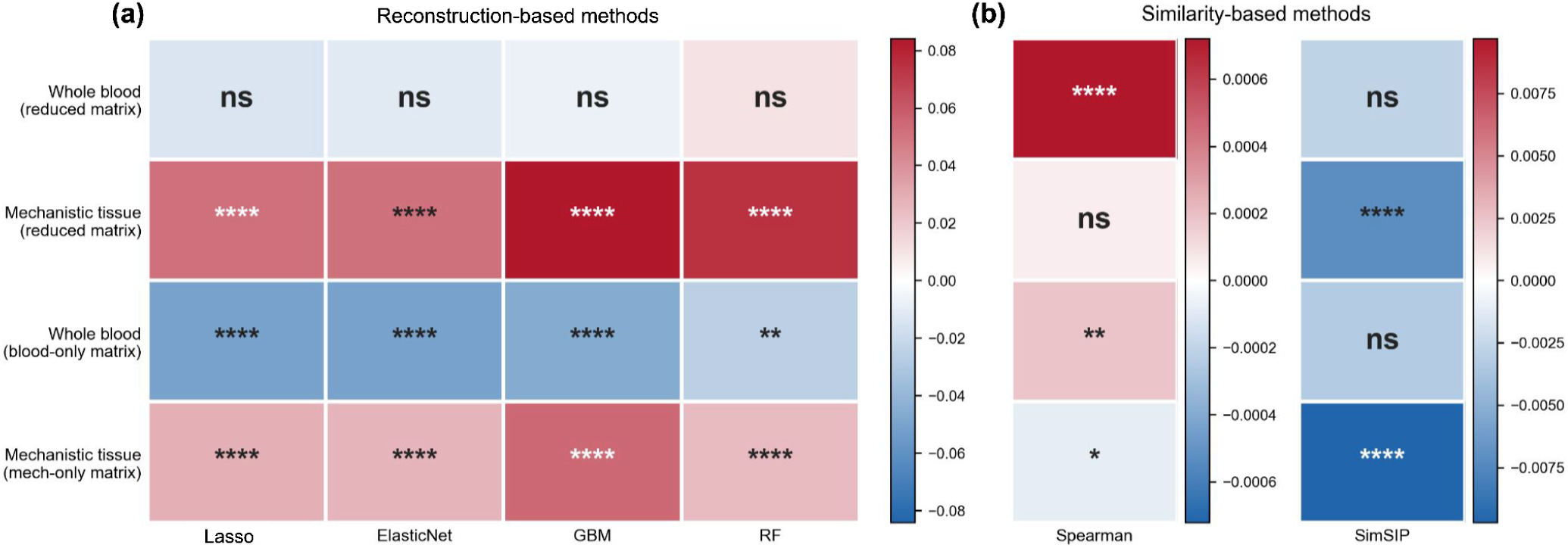
Removal of tissue-specific genes has relatively small effects on target disease reconstruction performance and similarity metrics. **(a)** Median paired differences in reconstruction Spearman between non-tissue and tissue holdouts (non-tissue holdout − tissue holdout) across reconstruction methods (Lasso, ElasticNet, GBM, and RF), shown for target diseases profiled in whole blood and mechanistic tissues from the reduced dataset, and for target diseases from the whole-blood-only and mechanistic-tissue-only datasets, respectively. **(b)** Median paired differences in Spearman and SimSIP similarities between non-tissue and tissue holdouts for the same target disease groups as in **(a)**. In both panels, heatmap colors indicate the median paired differences between non-tissue and tissue holdout values (non-tissue holdout − tissue holdout), with red indicating higher values under non-tissue holdouts and blue indicating higher values under tissue holdouts. For **(a)**, positive differences indicate more accurate target disease reconstruction using other diseases under non-tissue holdouts than under tissue holdouts. For both **(a)** and **(b)**, statistical significance of differences between the two holdout conditions was assessed using a paired, two-sided Wilcoxon signed-rank test with BH correction for multiple testing. BH-adjusted p-values (*P*) are denoted as follows: ns, not significant; *, *P* < 0.05; **, *P* < 0.01; ***, *P* < 0.001; and ****, *P* < 0.0001.

### 2.4 Assessment of biological relevance of feature importance

At each contribution threshold (**Methods 2.3**) per method, we obtained its contributing feature disease and obtained the corresponding Resnik similarity [20] of each feature disease and the target disease. We then calculated mean Resnik semantic similarity per contribution threshold within each method (**Figure 3b**). To evaluate whether overall the Resnik similarities of higher-ranked diseases (at a lower contribution threshold) were significantly higher than lower-ranked diseases (at a higher contribution threshold), we performed a series of statistical tests as described in **Methods S1.4**.

### 2.5 Assessment of tissue and organ system relationships among contributing disease signatures

To evaluate the overall effect of holdouts on feature prioritization, we analyzed the composition of tissue/system relationships of the feature disease signatures contributing to the predefined cumulative contribution thresholds (**Methods 2.3; Figure 4**). We only considered well-performing models (defined as models with reconstructed Spearman correlation of up-/downregulated gene groups > 0.1). This threshold was selected based on the ability of the reconstructed Spearman correlation to separate the real models from the shuffled models, which was highest at 0.1 among 7 thresholds tested (0, 0.01, 0.02, 0.05, 0.1, 0.2, 0.5) (**Methods S1.2; Figures S4, S5)**. We classified each target-feature disease signature pair into four mutually exclusive relationships based on their sample tissue and affected organ system: (i) same tissue and same system, e.g., Crohn’s (Gastrointestinal; GI) and Ulcerative Colitis (GI) profiled in blood (ii) same tissue and different system, e.g., Crohn’s (GI) and Type 1 diabetes mellitus (immune) profiled in blood (iii) different tissue and same system, e.g., Crohn’s (GI) profiled in blood and Inflammatory Bowel Disease 1 (GI) profiled in colon and (iv) different tissue and different system, e.g., Crohn’s (GI) profiled in blood and tuberculosis (respiratory) profiled in lung (**Figure 2c**). For each target disease_tissue signature, we identified disease signature features contributing to a specific cumulative contribution threshold: 0.1, 0.2, …, 0.9 representing 10–90% cumulative contribution, respectively (**Methods 2.3**), and which relationship they belong to as relationship fold enrichment. We defined relationship fold enrichment as observed/expected: the proportion of prioritized disease signatures belonging to a tissue/system relationship category at each cumulative contribution threshold divided by the corresponding proportion among all candidate disease signatures. For each method and contribution threshold, we calculated the mean fold enrichment across target disease_tissue signatures and applied a log2 transformation for interpretation and visualization. Positive log2(mean fold enrichment) values indicate overrepresentation, zero indicates no enrichment, and negative values indicate underrepresentation relative to background expectations (**Figure 4**). We also applied this procedure to assess the composition of signature types prioritized in different gene holdout experiments (**Methods 2.9**).

### 2.6 Assessment of same-disease similarity between whole-blood and mechanistic-tissue signatures

We assessed how sample tissue affects agreement among signatures of the same disease using diseases with at least two signatures from whole blood and at least two signatures from one mechanistic tissue (brain, lung, colon, skin or skeletal muscle), giving 385 signatures across 21 diseases and 42 unique disease_tissue pairs spanning five disease systems (**Figures 5a,b**). Within a disease, signatures were compared in one of three ways: blood-blood, mechanistic-mechanistic (mech-mech), or blood-mechanistic (blood-mech). Each signature pair was scored on two gene sets: (i) all 19,910 genes , and (ii) only tissue-specific genes (**Figure 5c, S11**). For each disease, the tissue-specific set was the union of the GTEx-derived whole blood genes and mechanistic tissue genes (**Figure S3**), and was used for all pairs so that the three ways of comparison were scored on the same genes. Similarity within a pair was quantified with two metrics: Spearman’s correlation coefficient and SimSIP. Spearman’s ρ was computed on average ranks of log₂FC. Its two-sided p-value was computed using the t-distribution approximation, evaluated in log space to avoid numerical underflow. SimSIP [7] was computed by ranking each signature’s genes by absolute log₂FC (strongest = 1, ties given average ranks), transforming ranks to weights 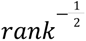, and taking the dot product of the two weight vectors. SimSIP p-values were one-sided, testing whether the observed score exceeded that expected under random pairing of genes. The score was standardized with its exact permutation mean and variance. As the null distribution is strongly right-skewed, tail probabilities were estimated by conditional Monte Carlo (10⁵ draws). In each draw, the placements of the 64 largest weights were sampled exactly, and the remaining terms were approximated as normal. These estimates agreed with brute-force permutation within 10–20% down to p ≈ 10⁻⁵. P-values were Bonferroni-corrected within each metric and gene set (m = 5,284). For each disease and tissue pairing, we report the median score across pairs. A pairing was called significant when the median Bonferroni-adjusted p-value was below 0.05, meaning at least half of its pairs were significant after correction.

### 2.7 Evaluation of predicted disease-disease relationships against known relevant disease associations

We paired diseases based on at least two of four independent evidence sources: shared drugs [21], shared genetic associations (GWAS studies) [22, 23], shared symptoms [24], and comorbidity [25] **(Figures 1b right, 9a, 2d; Methods S1.8)**. We calculated fold enrichment to determine how much greater than chance the known disease-disease relationships of a disease were predicted by each method (**Figures 6, 9**). For each target signature, we obtained the final disease-level ranking and retained top-ranked diseases at 10–90% cumulative contribution thresholds (**Methods 2.3**). We computed fold enrichment as the observed proportion of predicted disease pairs supported by external evidence relative to the expected proportion under a null distribution generated by permuting evidence labels 1,000 times for each target disease. Fold-enrichment values were log_2_-transformed for visualization, with negative values indicating observed proportions below the permutation-null expectation. Empirical *p*-values were calculated using a right-tailed permutation test with 1,000 permutations testing whether evidence-supported disease pairs are enriched among the contributing feature diseases at the specific contribution threshold. P-values were adjusted for multiple testing using the Benjamini-Hochberg (BH) false discovery rate (FDR) procedure. Statistical significance of depletion was not assessed, as the permutation tests were designed to evaluate enrichment only.

### 2.8 Gene holdout experiments for reconstruction-based approaches

#### 2.8.1 Tissue-gene-holdout strategy

We tested for transferability of signals across tissue-specific genes and non-tissue-specific genes following the confound-free train-test data split scheme described by Chyzhyk et al., [26]. This procedure of data splitting aims to provide prediction accuracy scores that are reflective of the training set independent of the confound variable (in our case, genes likely carrying tissue-specific signals). By including the target’s corresponding tissue-specific genes in only the test set, we can assess whether the tissue-specific genes drive the predictions. The tissue-specific genes were obtained by applying the classification algorithm proposed by Uhlén et al., 2015 on GTEx data (version 11; downloaded on March 28, 2026) using *teGeneRetrieval* function via *TissueEnriched* R package [27]. We defined tissue-specific genes as genes classified as “Tissue-Enriched”, “Group-Enriched”, and “Tissue-Enhanced” of one of the tissues. We defined non-tissue-specific genes as genes classified as “Not-Expressed”, “Mixed”, and “Expressed-In-All” (please see detailed definitions of these categories in Uhlén et al., (2015) [28]).

#### 2.8.2 Non-tissue gene holdout strategies

We established a matched control experiment complementing the tissue-gene-held-out models to evaluate the influence of tissue-specific genes on predictive performance while controlling for holdout gene-set size. The matched controls removed gene sets of identical size from non-tissue-specific genes, generating corresponding matched holdout test sets.

### 2.9 Gene holdout experiments for similarity-based approaches

For each target signature, we identified its associated GTEx tissue and retrieved the set of genes associated with that tissue. We then computed target-versus-all similarity scores under three gene-removal conditions: (i) tissue holdout: removal of the target’s tissue-associated genes (tissue-specific genes), (ii) non-tissue holdout: removal of an equally sized, randomly sampled set of genes drawn from the non-target-tissue gene pool, and (iii) random holdout: removal of an equally sized, randomly sampled set of genes drawn without restriction from the full gene set. For each condition, we recomputed rank-based Spearman correlation and SimSIP scores between the target signature and all other signatures in the dataset using the reduced gene expression matrix. Removing the same number of genes from each of the three holdout conditions allowed us to determine the specific contribution of tissue-specific genes to similarity scores relative to the effect of removing an equivalent number of genes at random (**Figure 7**).

### 2.10 Assessment of gene holdout effects on signature prediction ranks and their tissue/system relationship composition

We evaluated the changes in disease_tissue prediction composition with respect to the four tissue/system relationships described in **Methods 2.5 (Figures S16, S17)** and systematic changes in prediction ranks relative to the defined relationships described in **Methods S1.7 (Figures 8, S18, S19, S20, S21)**.

**Figure 8.**
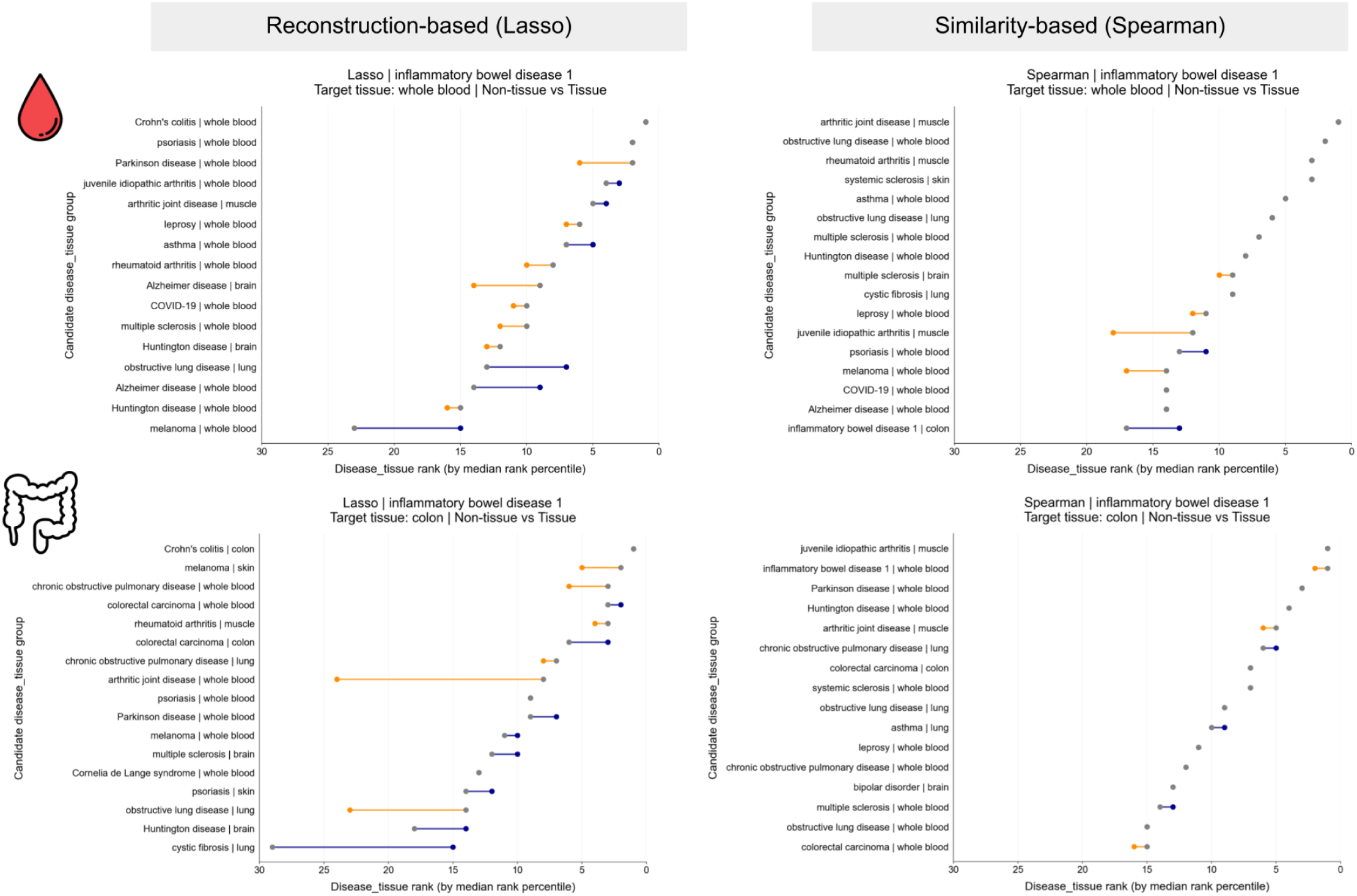
Tissue holdouts altered the prioritization ranks of disease_tissue predictions compared to non-tissue holdouts for IBD1 targets derived from whole blood and colon. Rank shifts of feature disease_tissue predictions between non-tissue and tissue holdouts for inflammatory bowel disease 1 (IBD1) targets derived from whole blood (top) and colon (bottom), using reconstruction-based Lasso (left) and Spearman similarity (right). Each panel displays the union of the top 15 ranked features under either holdout, selected independently for each method and target tissue. Disease-tissue features are ordered by their ranks under non-tissue holdouts. The x-axis shows ranks derived from median normalized rank percentiles across target signatures within each holdout, with rank 1 indicating the highest prioritization. Gray dots indicate ranks under non-tissue holdouts, while colored dots indicate ranks under tissue holdouts, connected by lines to illustrate rank shifts. Orange indicates decreased prioritization (higher rank), whereas blue indicates increased prioritization (lower rank) following the removal of tissue-specific genes.

## 3. Results

### 3.1 Overview of SigMatch

#### 3.1.1 SigMatch evaluation is powered by public disease transcriptomic signatures

A transcriptomic signature of a disease is not only a representation of disease, but also how the disease interacts with the sample’s tissue baseline: Δ*gene* = *disease* + (*disease* · *sample tissue*). These disease_tissue interaction effects could cause diseases affecting similar tissues to appear molecularly related even when the similarity is not driven by shared disease biology. SigMatch evaluates the impact of tissue context on the prioritization of disease relationships. We focused on 2,412 pre-calculated human RNAseq disease signatures from DiSignAtlas [4] (mapped to their corresponding terms in MONDO) (**Figure 1a**). The resulting dataset spans diseases affecting 12 individual organ systems, as well as multi-organ diseases (**Figure 1b left**). We aggregated signatures into disease_tissue units to facilitate biologically interpretable comparisons (see **Methods 2.3** for signature aggregation steps). We also inferred a ground truth of disease pairs supported by independent evidence sources (**Methods 2.7, S1.8; Figure 1b right**). An unsupervised low-dimensional projection of the 2,412 disease signatures revealed clustering by affected disease systems and by sample tissues (**Figure 1c,d**). This observation motivated our systematic evaluation of the impact of sample tissue on transcriptomic disease similarity while accounting for disease systems.

#### 3.1.2 SigMatch evaluates multiple transcriptomic disease signature similarity methods along three axes

The methods evaluated in SigMatch cover both similarity-based (comparing target disease signature with every other signature pairwise one-by-one; **Figure 2a**) and reconstruction-based methods (regressing target signature against a full matrix of signatures from other diseases, with genes as examples and signatures as features; **Figure 2b**). For the goal of prioritizing diseases related to a target disease (**Figure 2a,b**), similarity-based methods provide a score for every other disease signature while reconstruction-based methods select a subset of disease signatures that together recapitulate the target signature (**Figure 2b**). SigMatch presents an evaluation framework to systematically account for the impact of sample tissue and affected organ system for each disease pair (**Figure 2c**), and a novel ground truth dataset of biologically related disease pairs defined based on four sources of external evidence: shared i) genes, ii) drugs, iii) symptoms, and iv) comorbidities (**Figure 2d**). Using this SigMatch framework, we assess (**Figure 2e**): (i) predictive performance of reconstruction-based methods and whether each method prioritizes biologically related diseases as established in MONDO (**Methods 2.4**) [11], (ii) characteristics of the disease relationships prioritized by each method: whether the predictions are influenced by sample tissue and disease organ system (**Methods 2.5**) and whether they correspond to evidence-supported disease relationships and (iii) how much tissue-specific genes contribute to the predictions (**Methods 2.8, 2.9**), and impact of removing tissue-specific genes on recovery of evidence-supported disease pairs (**Methods 2.7, 2.10**). For evaluations (i) and (ii), we randomly removed genes from each target, with the number of removed genes matched to the size of the tissue-specific gene set associated with the target’s sample tissue. We refer to this scheme as “random holdout” (**Methods 2.2.3**), which served as a reference for the tissue-and non-tissue-gene removal experiments in (iii).

### 3.2 Reconstruction-based methods recapitulate target diseases using signatures from ontologically-related diseases; direct similarities also reflect ontological relatedness

We first evaluated the predictive performance of reconstruction-based methods (Spearman correlation between the predicted and the real target signature). This performance is an indicator of whether to trust if the model-selected diseases are biologically related to the target disease. All models consistently show a median predictive performance significantly higher than the shuffled models on all, up-, down-, and non-regulated genes in the real target signature (**Figure 3a**).

To evaluate the validity of the disease relationships unearthed by the reconstruction- and similarity-based methods, we assessed them against known disease-disease relationships in MONDO. Across the cumulative contribution thresholds (**Methods 2.3**), all reconstruction- and similarity-based methods tend to recover closely (ontologically) related diseases (relatively higher Resnik scores at lower contribution values) (**Figure 3b**). For instance, with Crohn’s colitis as target, all methods consistently prioritized ulcerative colitis and inflammatory bowel disease 1 (IBD1) (Resnik score of 5.755), and assigned low ranks to Parkinson disease, a nervous system disease (Resnik score of 0.68). With IBD1 as the target, Lasso and ElasticNet highly ranked hemorrhoids, a condition unrelated to IBD1 in the ontology (Resnik score of zero). The decreasing trends in Resnik scores with increasing contribution thresholds were statistically significant across methods (**Table S3**).

Though the median of mean Resnik scores did not exceed the optimal Resnik threshold at any cumulative contribution threshold (**Figure 3b)**, methods that exceeded this cutoff at cumulative contribution thresholds of 0.08 or lower are presented in **Figure S8**. The overall trend, along with the specific observations, suggests that these methods partly recover known related diseases and may help discover novel relationships.

### 3.3 Predicted disease signals were dominated by shared sample tissue

Given the observed clustering of signatures by sample tissue and affected disease system (**Figure 1c,d**), we next investigated if the recovery of ontologically-related diseases, which we observed above (**Results 3.2**) is driven by tissue signals over disease signals. We did this by partitioning each method’s prioritized disease relationships into four groups based on sample tissue and affected system (**Methods 2.5**). We found that, for every method, signatures of the same sample tissue showed higher enrichment than expected across all cumulative contribution thresholds (**Figure 4a,b top**). On the other hand, disease_tissue signatures from different tissues generally showed lower enrichment than same-tissue signatures, even when they belonged to the same disease system (**Figure 4a,b bottom**). Signatures differing in both tissue and disease system were consistently underrepresented relative to background expectations (**Figure 4a,b, bottom right**). Together, these findings suggest that shared sample tissue has a stronger influence on disease relationship prioritization than shared disease system.

### 3.4 Disease signatures from whole blood and mechanistic tissue are only weakly similar

To tease apart the tissue-dependent patterns, in a controlled manner, we narrowed down the full disease data (**Figure S9**) to the 21 diseases with signatures available from whole blood and the disease’s corresponding affected tissue (*i.e.*, mechanistic tissue) (**Methods 2.6; Figures 5a, S10**). This reduced dataset comprises disease signatures from whole blood and five mechanistic tissues: brain, lung, colon, skin, and muscle (**Figure 5a,b**) from five disease-affected organ systems: nervous, respiratory, gastrointestinal, integumentary, and musculoskeletal systems, respectively. While the full reduced 21-disease dataset was used for similarity-based methods, only eight target diseases met the reconstruction performance threshold and contained at least two whole-blood and two mechanistic tissue signatures (**Figure 5a**; bold). These were used with reconstruction-based methods and further downstream analyses and biological interpretation.

For each of the 21 diseases, using the similarity-based methods (Spearman and SimSIP), we first assessed the reproducibility of expression patterns across different signatures of the same disease, which in turn influences the stability and reliability of relationships to other diseases when used as targets (**Methods 2.6**). We found many diseases with blood signatures were significantly similar to each other with substantial effect sizes, as with mechanistic-tissue signatures, but blood and mechanistic-tissue signatures were only ever weakly similar to each other (median ρ near zero, although many remain statistically significant; **Figures 5c, S11**). This is consistent with the tissue-dependent effects in **Results 3.3**. Inflammatory bowel disease I (IBD1) and COVID-19 exhibited consistent signatures within blood and within mechanistic-tissue (IBD1: median Spearman’s ρ ≈ 0.19 and 0.18; median SimSIP ≈ 4.3 and 5.1, respectively; COVID-19: ρ ≈ 0.16 and 0.20; SimSIP ≈ 4.7 and 4.9).

### 3.5 Whole-blood signatures favor whole-blood relationships, with similar trends in mechanistically related tissues

Using the reduced dataset with disease signatures from whole blood and other five mechanistic tissues (**Figure 5b**), we repeated the tissue/system relationship enrichment analysis (**Methods 2.5**) with whole blood disease signatures as targets, and found that, in general, using whole blood signature as the target led to prioritizing other diseases profiled in whole blood, (**Figure 5d,e top left and right)**, and rarely prioritizing those profiled in mechanistic and other tissues, both regardless of whether the other disease was in the same system as the target or not (**Figure 5d,e bottom left and right)**. Similarly, mechanistic-tissue disease targets favor the same mechanistic tissue context over whole blood and other tissues independent of the disease system (**Figure S12**).

We closely examined these tissue-dependent patterns in IBD1 and COVID-19, two diseases that passed similarity- and reconstruction-based thresholds (**Results 3.4**), at the 50% cumulative contribution. The whole-blood-IBD1 target (GI) prioritized whole blood signatures of diseases across multiple systems: Crohn’s colitis (GI), Huntington disease (nervous), and cystic fibrosis (respiratory) (**Table S4**). In parallel, the colon-IBD1 target prioritized Crohn’s colitis profiled in colon tissues across methods (**Table S4**). This pattern was also observed in COVID-19. Whole-blood-COVID-19 target (respiratory) prioritized whole-blood signatures of diseases from various systems: cystic fibrosis (respiratory), leprosy (integumentary), Huntington disease (nervous), and systemic sclerosis (integumentary) (**Table S4**). Interestingly, the mechanistic-tissue target of COVID-19 consistently prioritized obstructive lung disease profiled in whole blood (respiratory) (**Table S4**).

For IBD1, the prioritization of Crohn’s colitis based on both whole blood and colon signatures is expected, as IBD1 represents the NOD2-associated form of Crohn’s disease [29]. The relationships with Huntington disease (degenerative) and cystic fibrosis (genetic/congenital) in whole blood perhaps reflect broader systemic inflammatory and stress responses beyond intestinal pathology. Similarly, for COVID-19, the prioritization of leprosy (infectious) suggests shared host responses to infection characterized by dysregulated interferon signaling, robust T-cell mediated response, and neutrophil activation [30–33], while the relationships with cystic fibrosis (genetic/congenital) and systemic sclerosis (fibrotic) reflect shared airway biology along with immune modulation [34]; Huntington disease (degenerative) could involve systemic inflammation. Lung-derived COVID-19 targets also prioritized whole-blood signatures from obstructive lung disease (non-autoimmune inflammatory), suggesting that respiratory inflammatory processes contribute to the predicted relationships across tissue contexts.

Overall, sample tissue dominates most transcriptomic-derived disease relationships, and relationships across different disease systems can be prioritized potentially based on shared systemic signals such as infection/immune response or broad inflammation beyond affected organ systems.

### 3.6 Predicted disease relationships show limited enrichment for evidence-supported ground truth under random holdout

Besides the observed dominant tissue effects, we examined how well the predictions captured independent evidence-supported disease relationships derived from shared information based on diseases’ treatments, genes, symptoms, and comorbidity (**Methods 2.7**). We evaluated only disease targets with at least one ground-truth-positive relationship among the available candidate diseases. We reduced false positives by defining a positive as a disease pair with at least two types of supported evidence. Of the 1,764 disease pairs supported by at least one evidence type, 92 were supported by at least two evidence types, of which 86 had available target disease signatures for downstream evaluation. We expanded this analysis to the full disease dataset (**Figures 1, S9**) given the sparse external evidence-supported relationships among the reduced dataset of 21 diseases (**Figures S10, S22**). Across methods, we identified 34 union unique target diseases that recovered at least one positive in the positive set (**Figure 6a**). We included all eligible target diseases in each method, without requiring them to be shared between methods. Enrichment of evidence-supported disease relationships (median) was below zero at the more selective cumulative contribution thresholds (0.1–0.4) across all methods. Even though enrichment increased and became positive at higher contribution thresholds, values overall remained close to zero, indicating limited enrichment relative to the permutation-based expectation (**Figure 6b**). This low enrichment trend for independent evidence of disease relationships suggests that predicted disease relationships that are highly dominated by same sample-tissue prioritization do not necessarily overlap with those supported by available external evidence.

### 3.7 Tissue-specific genes are not the sole drivers of transcriptomic disease relationship predictions

As tissue-specific genes are a likely driver of the outsized influence of sample tissue, we compared the difference in reconstruction model performance (or similarity score) with and without the genes specifically expressed in the sample tissue being included during model training (or similarity calculation).

All reconstruction-based models maintained positive predictive power regardless of whether genes specifically expressed in the sample tissue or those expressed in other tissues (labelled ‘non-tissue’) were removed during training (**Figure 7a**). Stratifying these results by sample tissue revealed that holding out sample-tissue-specific genes led to an increase (not always statistically significant) in performance for whole-blood targets and, conversely, led to a significant reduction in performance for mechanistic-tissue targets. These trends remained in the whole-blood-only and mechanistic-tissue-only datasets (**Figure 7a**), with variation in the statistical significance of these differences across datasets and disease systems of different sample sizes (**Figure S13**).

With similarity-based methods, removing sample-tissue-specific genes resulted in small changes with variable statistical significance across methods and data subsets (**Figure 7b**).

Nevertheless, across all methods, despite having variable impact on performance/similarity, holding out sample-tissue-specific genes did not significantly impact the ontology-based relatedness of the prioritized disease relationships or the dominance of shared sample tissue contexts among them (**Figures S15–S17**).

Revisiting IBD1 (with targets derived from whole blood and colon, and the mechanistic tissue of the GI system), we observed that tissue holdouts did not consistently deprioritize disease_tissue features sharing the same sample tissue as the target. Crohn’s colitis remained highly prioritized for both whole blood and colon IBD1 targets across multiple methods, though other disease_tissue features showed different magnitudes of rank shifts. The shifts are also variable across methods: Lasso, ElasticNet, and GBM show larger shifts for selected features than RF, Spearman, and SimSIP (**Figures 8, S18**).

A similar pattern was observed for COVID-19 targets derived from whole blood and lung (**Figure S19**). Cystic fibrosis derived from whole blood was highly ranked across holdouts and several methods. Conversely, other disease_tissue features underwent unstable rank shifts. These changes were also method-dependent, with larger shifts observed for selected features for Lasso and ElasticNet, compared to generally smaller changes for RF and Spearman (**Figure S19**).

These two cases illustrated how removing tissue-specific genes can affect disease_tissue prioritization differently across methods. This supported the systematic rank-shift analysis (**Methods 2.10; Figures S20**) where we found no consistent reduction in the prioritization of signature relationships from the same sample tissue.

### 3.8 Evidence-supported disease relationship enrichment did not differ significantly between tissue and non-tissue holdouts across methods

To evaluate the impact of our tissue and non-tissue holdouts on disease relationship prioritization, we compared disease predictions from all methods performed on the full dataset against external-evidence-supported disease relationships derived from four independent sources, as described in **Results 3.6** (**Methods 2.7, S1.8; Figure S23**). Restricting the analysis to target diseases shared between tissue and non-tissue holdouts within each method, and taking the union across methods, we identified 28 unique target diseases with at least one ground-truth-positive relationship (**Figure 9a**).

**Figure 9.**
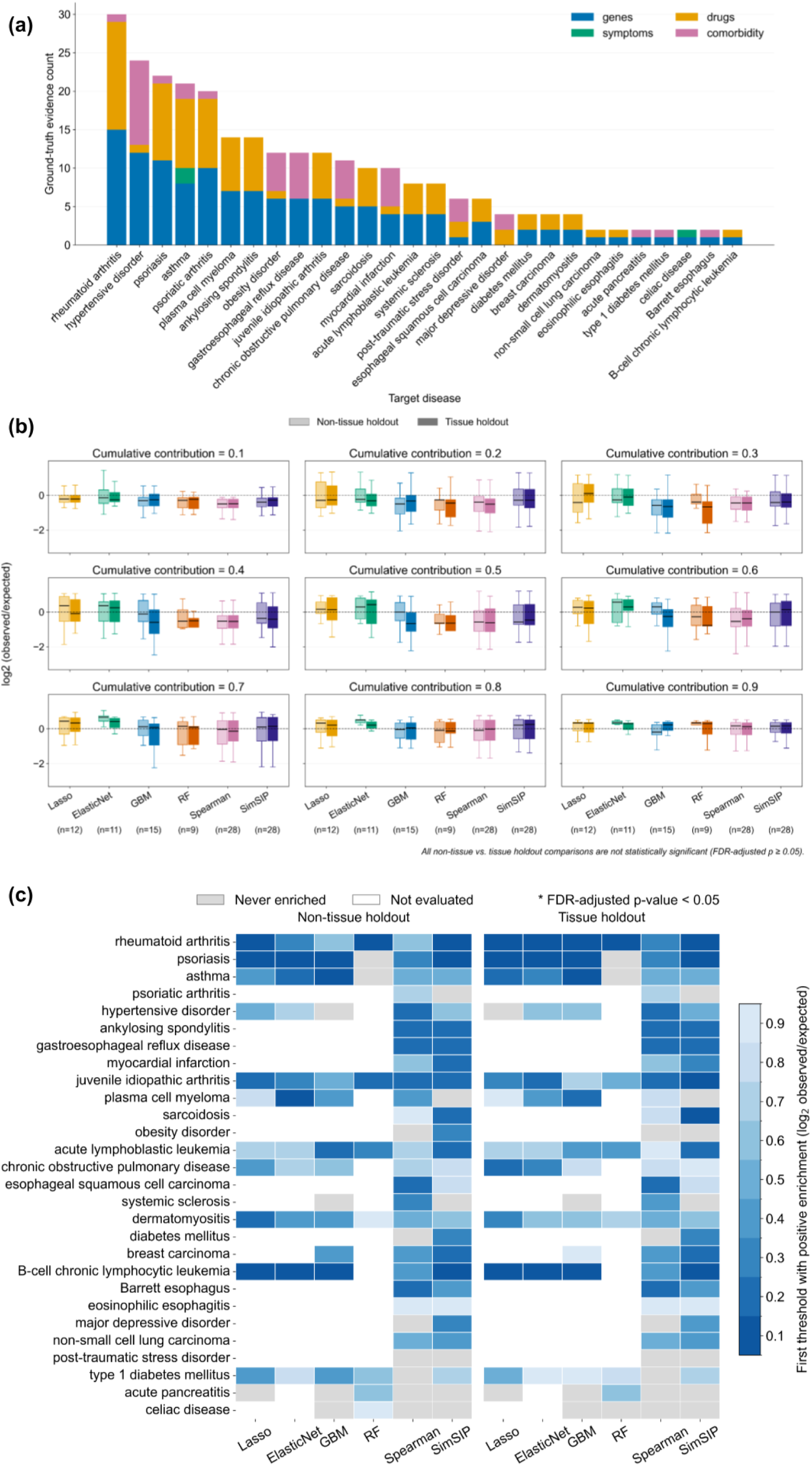
Tissue and non-tissue holdouts show no significant differences in the enrichment of contributing diseases for evidence-supported disease relationships. **(a)** Distribution of evidence types across target diseases based on shared genes, symptoms, drugs, and comorbidity. Only diseases with at least one ground-truth-positive relationship (supported by ≥2 evidence types) are shown. Disease pairs may belong to multiple evidence categories. **(b)** Distributions of enrichment: log_2_(observed/expected) (y-axis) across cumulative contribution thresholds (0.1–0.9) for all the methods (x-axis). Lighter and darker shades represent non-tissue and tissue holdouts, respectively. Boxplots summarize enrichment across matched target diseases with available positive evidence in both holdouts (n). Paired Wilcoxon signed-rank tests compare holdouts within each method and threshold, with BH correction (ns, adjusted *p*-value ≥ 0.05). **(c)** Heatmaps showing the earliest contribution threshold at which each target disease (row) exhibits positive enrichment across methods (columns). Target diseases are ordered by the mean number of evidence-supported disease relationships recovered across methods and holdouts at a cumulative contribution threshold of 0.9, from highest to lowest. Darker blue indicates enrichment at an earlier threshold; gray indicates evaluated but never enriched, and white indicates not evaluated. None of the positive enrichment is statistically significant.

For each method, we identified shared target diseases across holdouts (**Methods 2.8, 2.9**) and compared their enrichment in evidence-supported ground truths across cumulative contribution thresholds (**Figure 9b**). Although we observed some positive enrichment at higher thresholds in some methods, the degree of enrichment remained low across holdouts, where the tissue holdouts did not show significant improvement in predicting evidence-supported disease relationships over the non-tissue holdouts (**Figure 9b**). We also examined evidence enrichment among the individual target diseases across holdouts and methods at each cumulative contribution threshold (**Figure 9c**). Our findings showed that the enrichment patterns across methods and thresholds of tissue and non-tissue holdouts were conserved (**Figure 9c**). For instance, at relatively low cumulative contribution thresholds, rheumatoid arthritis, psoriasis, and asthma showed positive enrichment across multiple methods in both holdouts. Juvenile idiopathic arthritis and dermatomyositis exhibited positive enrichment across several methods, although the earliest thresholds varied between methods and holdouts.

We also investigated the evidence-supported disease relationships predicted at the 50% contribution threshold by at least one method under tissue and non-tissue holdouts. Most of the disease pairs were identified by both holdouts across the methods (**Figure S24**). This observed pattern was consistent across the disease relationships belonging to the same and different organ systems (**Figure S24**). The target → feature relationships predicted exclusively under the tissue holdout setting by at least one of the six methods are hypertensive disorder and psoriasis, hypertensive disorder and obesity disorder, and juvenile idiopathic arthritis and psoriasis. The target → feature relationships predicted exclusively under the non-tissue holdout setting by at least one of the six methods include hypertensive disorder and rheumatoid arthritis, psoriasis and ankylosing spondylitis, dermatomyositis and psoriasis, rheumatoid arthritis and hypertensive disorder, and psoriasis and dermatomyositis. These unique relationships did not reveal a clear pattern in either holdout setting based on whether the diseases belonged to the same or different organ systems. Across available target diseases and ground truths, we found no statistically significant difference between holding out tissue-and non-tissue-specific genes in the evidence-supported disease relationships predicted across methods.

## 4. Discussion

Different underlying sample tissues are known to distort the measured disease gene expression [13]. These tissue effects can still be present at the disease expression signature level as part of combinatorial effects of disease and tissue. This poses a challenge for identifying transcriptomic disease relationships, as similarities between disease signatures could reflect shared sample tissue rather than shared disease characteristics. Our study systematically evaluated how sample tissue influences transcriptomic similarities defined by methods of varying complexity. Sample tissue strongly drove transcriptomic disease relationships across all methods, and removing tissue-specific genes did not mitigate this effect.

### Key findings

Disease differential expression signatures can retain tissue-dependent transcriptional responses, even when derived using tissue-matched controls. Our work assessed the impact of sample tissue on similarities across diseases at a large scale using uniformly processed public disease differential expression signatures. Using existing similarity-and reconstruction-based methods incorporating feature selection principles, our analyses showed that shared tissue context strongly influenced transcriptomic disease similarity prioritization. The tissue effect was consistent across all methods regardless of complexity, including rank-based correlation (Spearman) [6, 14], rank-based similarity (SimSIP) [7], and reconstruction-based methods employing linear and tree-based regression models (Lasso, ElasticNet, GBM, and RF) (**Figure 4**). This extends the known tissue dependence of disease signatures to similarities between diseases.

Comparing two specific sample tissue types: whole blood and mechanistic tissue, defined as organ tissue affected by the disease, our results showed that disease responses depended strongly on the sample tissue, even within the same disease (**Figures 5c, S11**). This may be either because the tissue effect is substantial enough to obscure disease responses or because disease responses are tissue-specific, as established by many studies [35–38] that disease responses could be tissue-specific. Our gene holdout experiments, designed to exclude tissue-associated genes from disease similarity predictions, did not mitigate the observed tissue effects, as shown by the persistent dominance of same-tissue predictions (**Figures S16, S17**), their biological relevance within the disease ontology (**Figure S15**), and the effect sizes of the predicted similarity scores (**Figures S20, S21**). This suggests that excluding GTEx-defined tissue-specific genes is not sufficient to eliminate tissue effects, possibly because of more complex transcriptional structure such as gene co-expression patterns. Overall, our findings show that tissue context should be considered when interpreting transcriptomic disease relationships and their potential for knowledge transfer.

### Limitations and future directions

Our study used public RNAseq disease transcriptomic signatures and disease/tissue labels from DiSignAtlas [4], which carry several sources of heterogeneity among signatures representing the same disease, such as differences in disease states, differential expression comparisons, sample groups that do not necessarily represent disease vs. healthy controls, and study-specific variation. Additionally, signatures originating from the same study and pairwise comparisons sharing individual signatures were not explicitly accounted for as non-independent observations in our statistical analyses. Although we applied multiple-testing correction, this does not address potential study-level confounding. In particular, same-tissue comparisons may be more likely to involve signatures from the same study than cross-tissue comparisons, potentially contributing to the observed differences in within-disease signature similarity. We harmonized tissue labels by mapping the original tissue labels to high-level GTEx [13] tissue labels and performed analyses at the disease_tissue level. This standardization could lose more specific tissue information that matters when interpreting the results. Disease system labels, assigned with a large language model and manually validated (**Methods 2.1**), were restricted to the affected organ system of each disease, while additional associated organ systems were noted for result interpretation.

We identified tissue-specific genes using independent healthy tissue expression data across all available tissues in GTEx. However, genes identified using healthy tissue expression may not fully represent tissue-associated expression patterns under disease conditions. Our study demonstrates strong tissue dependence but does not establish how much of that dependence comes from tissue-specific disease biology, cell-type composition, or other study-related variation. Also, removing tissue-specific GTEx genes alone does not account for `disease x tissue` interaction term.

Due to limited ground-truth disease relationship coverage in the external evidence enrichment analysis, we could not conclude whether disease relationships lacking external evidence in the tissue and non-tissue gene holdout experiments were biologically unrelated or simply not represented in the available reference data. With <28 target diseases per method shared between holdouts (**Figure 9**), a non-significant difference also cannot be overinterpreted. Future studies should use more consistently defined disease differential expression comparisons and more detailed tissue and cell-type annotations to better distinguish tissue-specific disease responses from other sources of variation. More comprehensive reference data on known disease relationships could improve the evaluation of biological relevance and allow a more reliable comparison of predictions from tissue and non-tissue gene holdout experiments.

## 5. Conclusion

This study systematically evaluated the impact of sample tissue on disease relationship prioritization using a large collection of public transcriptomic disease signatures and computational approaches of varying complexity. Our findings demonstrated a strong influence of shared sample tissue on predicted disease similarities across all methods. Removing tissue-specific genes did not reduce tissue dependence in disease similarity predictions or improve the recovery of evidence-supported disease pairs. Our study extends the known tissue dependence of disease signatures to inferred relationships between diseases and emphasizes the importance of considering sample tissue when interpreting transcriptomic disease similarities and their biological relevance.

## Acknowledgement

We would like to thank Dr. Evan Brenner and Kat Terwelp for critical feedback on SigMatch through its development course, and Keenan Manpearl and Parker Hicks for UMLS-MONDO mapping and assistance with Resnik similarity calculation.

## Funding

This work was supported by the NIH NIAID R21 AI169301 awarded to AK and JR, the Colorado Translational Research Scholars Program to JR, and CU Anschutz start-up funds awarded to AK and JR.

## Notes

### Competing Interest Statement

The authors have declared no competing interest.

