## Supplemental material for "Systematic evaluation using SigMatch reveals how sample tissues shape transcriptomic disease relationships"

**1. Supplementary material and methods** **3**
- 1.1 Additional details on tuning scheme 3
- 1.2 Sensitivity analysis to determine the optimal performance threshold for reconstruction-based methods in downstream disease relationship analyses 3
- 1.3 Sensitivity analysis to determine a Resnik score threshold that best separates diseases 3
- 1.4 Statistical tests for decreasing trends in Resnik scores across cumulative contribution thresholds 4
- 1.5 Inclusion criteria for the reduced, whole-blood-only, and mechanistic-tissue-only disease datasets 4
- 1.6 Tissue/system relationship classification for whole-blood/mechanistic-tissue targets 4
- 1.7 Evaluation of systematic changes in prediction ranks relative to the defined tissue/system relationships 5
- 1.8 Disease relevance data and preprocessing 5

**2. Supplementary figures** **6**
- Figure S1. UMAP projection of cancer and non-cancer disease signatures showed substantial overlap with no clear separation between the two groups. 6
- Figure S2. Tuning settings using representative disease signatures to find optimal hyperparameters. 7
- Figure S3. Number of transcriptomically tissue-associated genes identified using GTEx data. 8
- Figure S4. Sensitivity analysis of reconstruction Spearman thresholds for distinguishing real from shuffled model performance. 10
- Figure S5. Sensitivity analysis of reconstruction Spearman thresholds based on the number of target diseases passing the evaluated thresholds. 10
- Figure S6. Diseases from the same system are significantly more related in the disease ontology compared to different systems. 11
- Figure S7. Sensitivity analysis to determine the optimal Resnik similarity threshold for distinguishing disease pairs from the same and different disease systems. 11
- Figure S8. Median Resnik similarity among disease relationships at lower cumulative contribution thresholds. 12
- Figure S9. Distribution of disease signatures across tissues and disease systems in the full dataset. 13
- Figure S10. Selection of tissue-disease pairs for the reduced, whole-blood-only, and mechanistic-tissue-only datasets. 14
- Figure S11. Within-disease agreement across whole-blood and mechanistic-tissue signatures using all genes and tissue-associated genes. 15
- Figure S12. Mechanistic tissue-disease targets prioritize mechanistic tissue-disease features. 16
- Figure S13. Effects of tissue-specific holdout on reconstruction performance and similarity scores across disease systems. 18
- Figure S14. Number of target signatures and diseases retained across tissue holdout settings and filtering steps using the full disease dataset. 20
- Figure S15. Comparison of mean Resnik similarity between tissue and non-tissue holdout settings across cumulative contribution thresholds using the full dataset. 21
- Figure S16. Enrichment of tissue and disease system relationships among prioritized disease features of reconstruction-based methods across cumulative contribution thresholds using reduced datasets. 21

Figure S17. Enrichment of tissue and disease system relationships among prioritized disease features of similarity-based methods across cumulative contribution thresholds using reduced datasets. 22
Figure S18. Tissue holdouts altered the prioritization ranks of feature disease_tissue predictions compared to non-tissue holdouts for IBD1 targets derived from whole blood and colon. 23
Figure S19. Tissue holdouts altered the prioritization ranks of feature disease_tissue predictions compared to non-tissue holdouts for COVID-19 targets derived from whole blood and lung. 25
Figure S20. Shifts in feature disease_tissue prioritization between non-tissue and tissue holdouts across tissue and disease system relationship categories using reduced datasets. 25
Figure S21. Shifts in feature disease_tissue prioritization between non-tissue and tissue holdouts across tissue and disease system relationship categories, stratified by target disease system, using reduced datasets. 26
Figure S22. Comparison of biological evidence coverage between the full and reduced datasets. 27
Figure S23. Distribution of positive disease pairs across target diseases supported by multiple relevance evidence types in the full dataset. 28
Figure S24. Evidence-supported pairs predicted at the contribution threshold of 0.5 by at least one method under either the tissue or non-tissue holdouts. 30
**3. Supplementary tables 30**
Table S1. Hyperparameter tuning details for reconstruction-based methods. 30
Table S2. Summary of feature summarization criteria across analyses. 31
Table S3. Statistical tests of changes in mean Resnik similarity across cumulative contribution thresholds. 31
Table S4. Prioritized disease_tissue features for whole-blood and mechanistic-tissue IBD1 and COVID-19 target signatures. 33
Table S5. Data sources used to identify evidence-supported disease relationships. 34
**References 34**

### 1. Supplementary material and methods

#### 1.1 Additional details on tuning scheme

As cancer and noncancer signatures did not show clear separation in the UMAP representation (**Figure S1**), we performed hyperparameter tuning under the combined setting (**Figure S2**): combined cancer and noncancer signatures. We used $k$-means clustering to identify representative target signatures from a pool of diseases, including cancers, for hyperparameter tuning (**Figure S2)**. The optimal hyperparameters obtained from the representative targets were aggregated using a median estimator and subsequently applied to nonrepresentative targets for model evaluation.

#### 1.2 Sensitivity analysis to determine the optimal performance threshold for reconstruction-based methods in downstream disease relationship analyses

In addition to evaluating the performance of the real reconstruction-based models, we shuffled gene values within each target signature and repeated the procedure to establish baseline performance. We used these baselines to determine whether a reconstruction-based model's performance was sufficient to interpret its prioritized disease relationships. We performed a sensitivity analysis to choose a reconstruction Spearman correlation threshold (a Spearman correlation indicating level of agreement between the target signature and the predicted signature) that best distinguished reconstruction performance between real and shuffled models. We evaluated thresholds of 0, 0.01, 0.02, 0.05, 0.1, 0.2, and 0.5. We defined real models as the positive class and the shuffled models as the negative class. For each threshold $t$, we calculated classification accuracy as demonstrated in **Equation S1**:

**Equation S1:** $Accuracy = \frac{TP+TN}{TP+FN+FP+TN}$

where $TP$ (true positives) and $FN$ (false negatives) represent the numbers of real models with Reconstruction Spearman ≥ $t$ and < $t$, respectively. $FP$ (false positives) and $TN$ (true negatives) represent the numbers of shuffled models with Reconstruction Spearman ≥ $t$ and < $t$, respectively.

We used accuracy because the numbers of positives and negatives were balanced (paired real and shuffled model performances). A reconstruction Spearman threshold of 0.1 yielded the highest classification accuracy for every reconstruction-based model and gene group (**Figure S4**). We selected this threshold for downstream analyses, considering both classification accuracy (**Figure S4**) and the number of target diseases retained (**Figure S5**).

#### 1.3 Sensitivity analysis to determine a Resnik score threshold that best separates diseases

In our biological relevance analysis of the predicted feature diseases (**Methods 2.4**), we used Resnik similarity as a proxy for disease similarity within the ontology structure. Disease relatedness is based on the ontology concept that diseases sharing a more specific common ancestor term with fewer descendants are more closely related than diseases sharing a broader ancestor term with more descendants. MONDO also classifies diseases according to their related organ systems. We verified that, overall, diseases from the same system had significantly higher Resnik similarity scores than diseases from different systems (**Figure S6a**). We observed this pattern consistently across different disease systems (**Figure S6b**). We used the classification of the same/different system to identify a baseline Resnik score for ontology-related diseases in **Figure 3b**. We evaluated all unique observed Resnik scores as potential thresholds for separating disease pairs from the same and different systems. For each threshold, we classified disease pairs with Resnik scores greater than or equal to the threshold as belonging to the same system and those with scores below the threshold as belonging to different systems. We considered same-system disease pairs as positives and different-system disease pairs as negatives. Both accuracy (**Equation S1**) and balanced accuracy (**Equation S2**) were calculated for each threshold (**Figure S7a**). We used balanced accuracy (**Figure S7b**) to account for the imbalance in the numbers of positive and negative disease pairs. The selected Resnik score threshold was the one that maximized balanced accuracy which was the optimal threshold for separating disease pairs from the same and different systems.

**Equation S2**: $Balanced\ accuracy\ = \frac{(Sensitivity + Specificity)}{2}$

#### 1.4 Statistical tests for decreasing trends in Resnik scores across cumulative contribution thresholds

As part of our biological relevance analysis of the predicted feature diseases (**Methods 2.4**), we evaluated whether the mean Resnik similarity of predicted feature diseases decreased as the cumulative contribution threshold increased. As the disease sets selected at different thresholds were nested, we accounted for repeated measurements of the same target disease. For each method, we compared mean Resnik similarity between adjacent cumulative contribution thresholds using one-sided paired Wilcoxon signed-rank tests. We included only target diseases with available Resnik similarity values at both thresholds. We adjusted the resulting p-values for multiple testing within each method using the Holm procedure (**Table S3**).

#### 1.5 Inclusion criteria for the reduced, whole-blood-only, and mechanistic-tissue-only disease datasets

For all curated datasets, we only considered signatures from whole blood tissue or the disease's corresponding mechanistic tissue. First, we filtered our dataset to diseases with at least 2 signatures in both whole blood and mechanistic tissue. To select disease systems with adequate examples, we excluded all diseases belonging to a system with fewer than 3 remaining diseases. This resulted in a reduced dataset comprising 21 diseases, spanning five disease systems defined according to the affected organ systems (**Figures 5a**, **S10**). Similarly, to create the whole-blood-only and mechanistic-tissue-only datasets, we filtered to diseases with at least 2 signatures in the respective tissue, dropping systems with fewer than 3 remaining diseases.

#### 1.6 Tissue/system relationship classification for whole-blood/mechanistic-tissue targets

The predefined mechanistic tissue-system relationships were "Brain" for nervous system diseases, "Colon" for gastrointestinal system diseases, "Lung" for respiratory system diseases, "Skin" for integumentary system diseases, and "Muscle" for musculoskeletal system diseases. For the tissue/system relationship enrichment analyses using whole-blood and mechanistic-tissue disease targets separately, we established more specific tissue categories for candidate disease signatures based on their tissue origin relative to the target disease system. Candidate signatures were classified as originating from whole blood, the mechanistic tissue corresponding to the target disease system, or other tissues. Each candidate signature was further classified according to whether its disease system matched that of the target, resulting in six mutually exclusive tissue/system relationship categories: Whole Blood + same system, Whole Blood + different system, Mechanistic tissue + same system, Mechanistic tissue + different system, Other tissue + same system, and Other tissue + different system.

However, we ended up with only four relationship categories, as each disease system was associated with only one mechanistic tissue. Therefore, there were no candidate disease_tissue features from the target's mechanistic tissue but a different disease system, or from other tissues within the same disease system. The four remaining relationships were whole blood with the same system, whole blood with a different system, mechanistic tissue with the same system, and other tissue with a different system (**Figures 5d,e, S12**).

#### 1.7 Evaluation of systematic changes in prediction ranks relative to the defined tissue/system relationships

For each target, we converted prediction ranks into rank percentiles, and calculated the change in rank percentile for each candidate feature by subtracting its non-tissue holdout percentile from its tissue holdout percentile (Tissue – Non-tissue holdouts). Each candidate feature was then classified into one of four tissue/system relationship categories based on whether its tissue and system matched those of the target. The mean rank-percentile change was calculated across all candidate features within each relationship category for each target signature. These mean changes were subsequently averaged across target signatures belonging to the same disease_tissue pair. Finally, the median of these disease_tissue-level mean changes was calculated for each relationship category to evaluate systematic shifts in feature prioritization.

#### 1.8 Disease relevance data and preprocessing

We accessed the datasets in **Table S5** on June 30, 2026. We harmonized disease identifiers to MONDO using the corresponding identifier mappings, including DOID for genetic associations and symptoms, UMLS CUI for drugs, and ICD10 identifiers mapped through DOID for comorbidity.

For each type of shared information between diseases (drugs, genes, and symptoms; here referred to as units "unit") used to indicate disease relevance (**Table S5**), we applied the same procedure to evaluate the significance of overlap between disease pairs. We defined the background universe separately for each information type as the union of all unique units associated with at least one disease. For each disease pair, we calculated the number of units associated with each disease and the number of shared units. We evaluated the statistical significance of the observed overlap using the upper-tail hypergeometric test (**Equation S3**), which calculated the probability of observing the same number or more shared units given the corresponding background universe and the sizes of the two disease-specific unit sets.

We calculated three complementary measures of overlap for each information type: fold enrichment (**Equation S4**), Jaccard index (**Equation S5**), and overlap coefficient (Szymkiewicz-Simpson coefficient; **Equation S6**). For each disease pair, we defined $A$ and $B$ as the sets of units associated with the two diseases, $k = |A \cap B|$ as the number of shared units, and $N$ as the total number of unique units in the corresponding background universe.

**Equation S3:** $P(X \geq k) = \sum_{i=k}^{\min(|A|,|B|)} \frac{\binom{|A|}{i}\binom{N-|A|}{|B|-i}}{\binom{N}{|B|}}$

**Equation S4**: $Fold\ enrichment = \frac{k}{E(k)}$

where $E(k) = \frac{|A| \times |B|}{N}$ is the expected overlap under random assignment.

**Equation S5:** $Jaccard\ index = \frac{|A \cap B|}{|A \cup B|}$

**Equation S6:** $Overlap\ coefficient = \frac{|A \cap B|}{\min(|A|,|B|)}$

For each unit, we adjusted the hypergeometric p-values for multiple testing using the Benjamini-Hochberg (BH) false discovery rate (FDR) procedure. For each information type, we considered disease pairs to have significant overlap if they shared at least one unit and had an FDR-adjusted p-value below 0.05.

For comorbidity, we used the ICD10 adjacency matrices released by Dervić et al., 2025 [1]. We calculated the median estimated odds ratios across the available matrices representing sex and years. We mapped the ICD10 identifiers to MONDO and considered disease pairs with positive median values as having comorbidity and those with zero or missing values as having no comorbidity. We did not apply the hypergeometric overlap test to comorbidity data.

#### 2. Supplementary figures

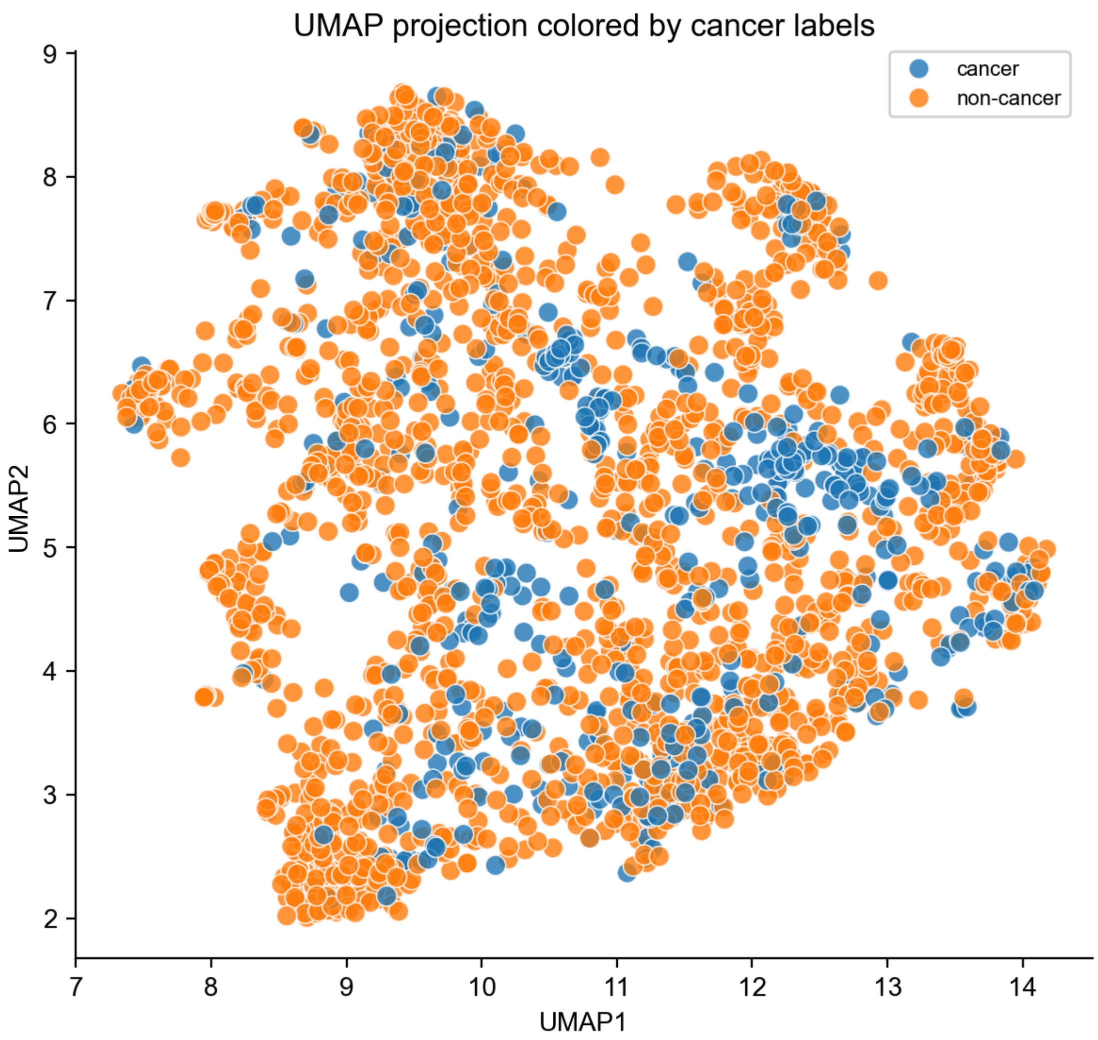


Figure S1. UMAP projection of cancer and non-cancer disease signatures showed substantial overlap with no clear separation between the two groups.

Two-dimensional Uniform Manifold Approximation and Projection (UMAP) of disease signatures, colored by cancer (blue) and non-cancer (orange) labels. Each point represents a disease signature.

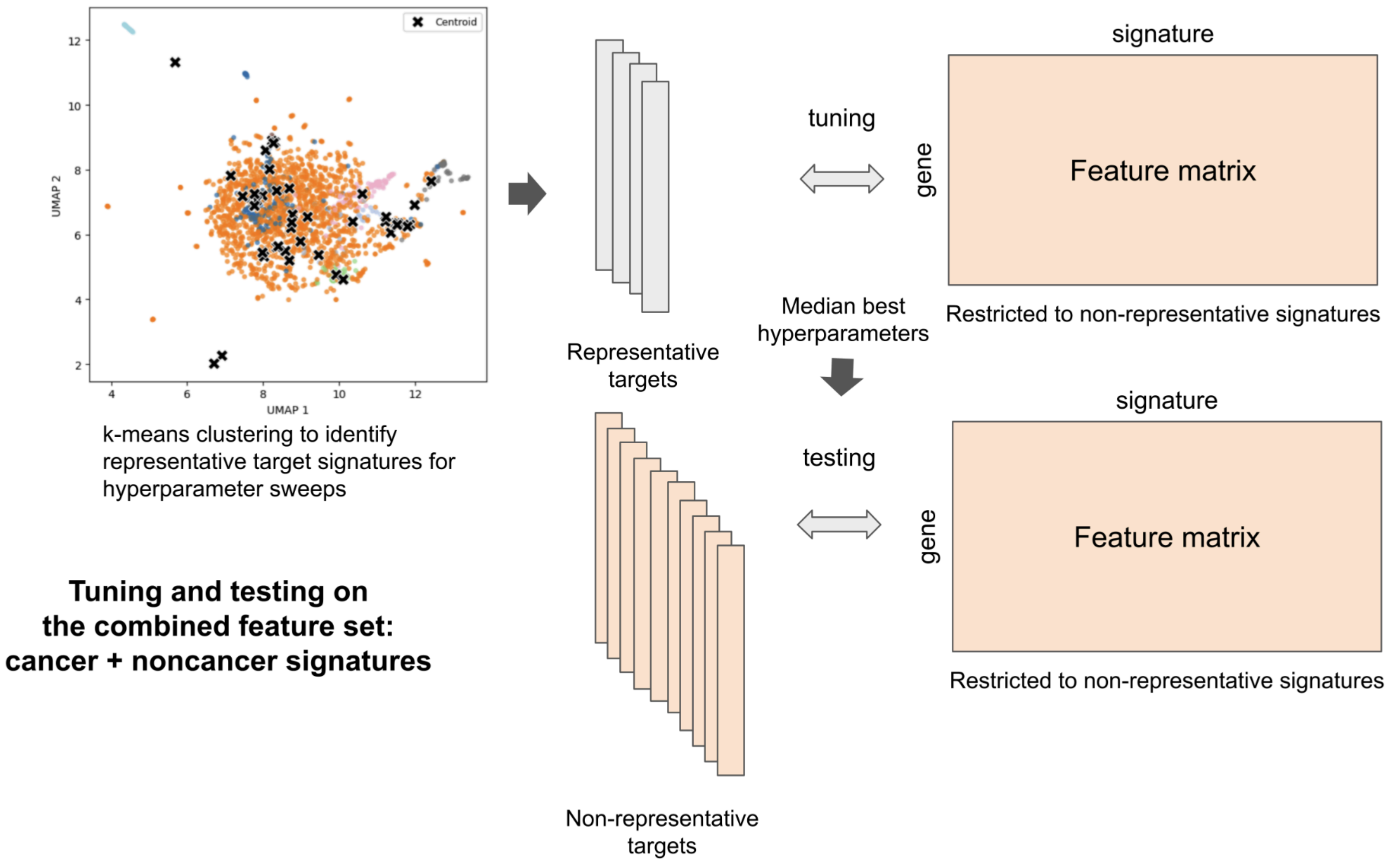


Figure S2. Tuning settings using representative disease signatures to find optimal hyperparameters.

Combined disease signatures were included in tuning and testing. Representative targets could belong to any disease, while non-representative targets were used for testing. The feature matrices for both tuning and testing were restricted to non-representative signatures.

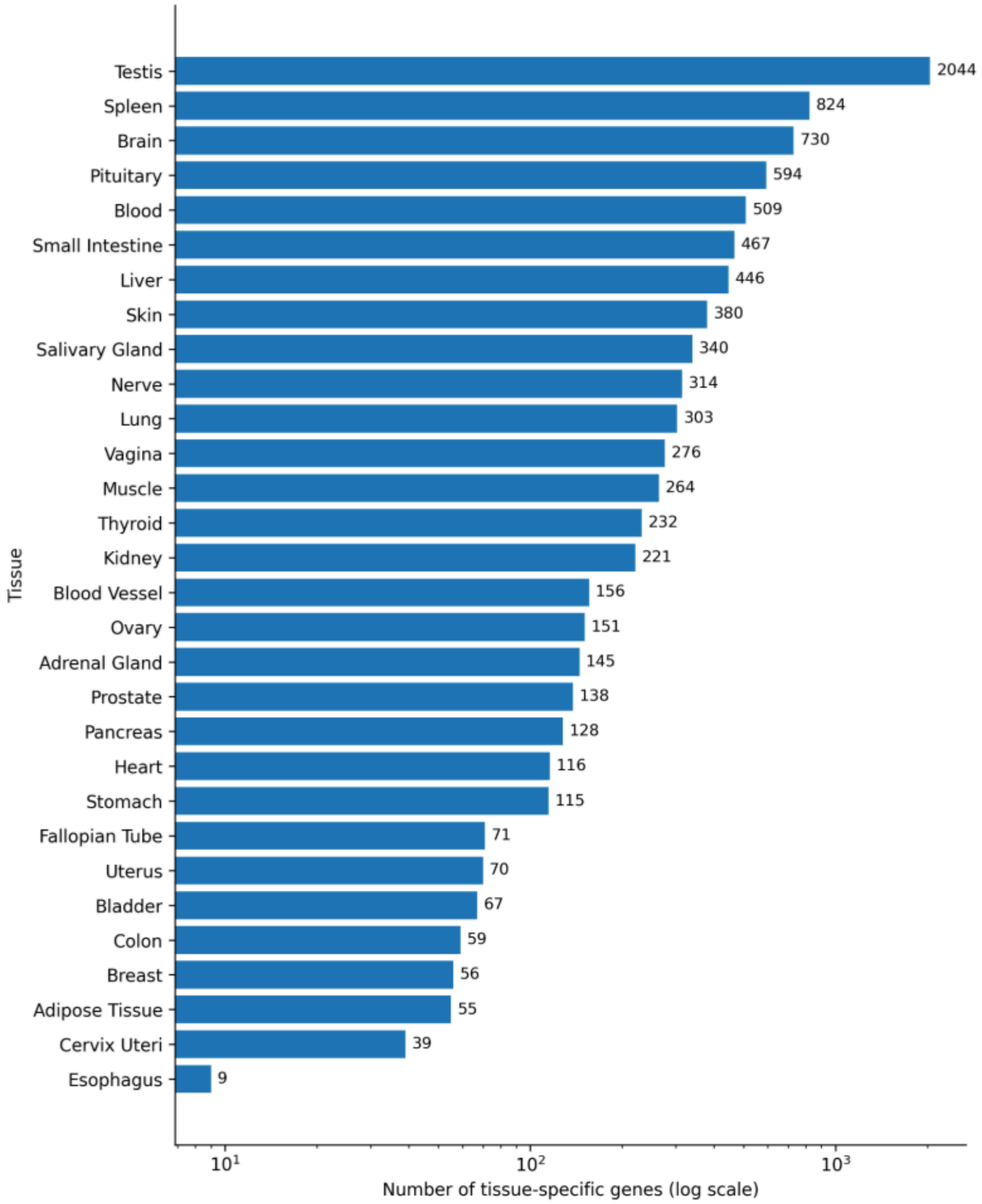


Figure S3. Number of transcriptomically tissue-associated genes identified using GTEx data.

Number of identified tissue-associated genes on a log scale (x-axis) across 30 GTEx high-level tissue names (y-axis).

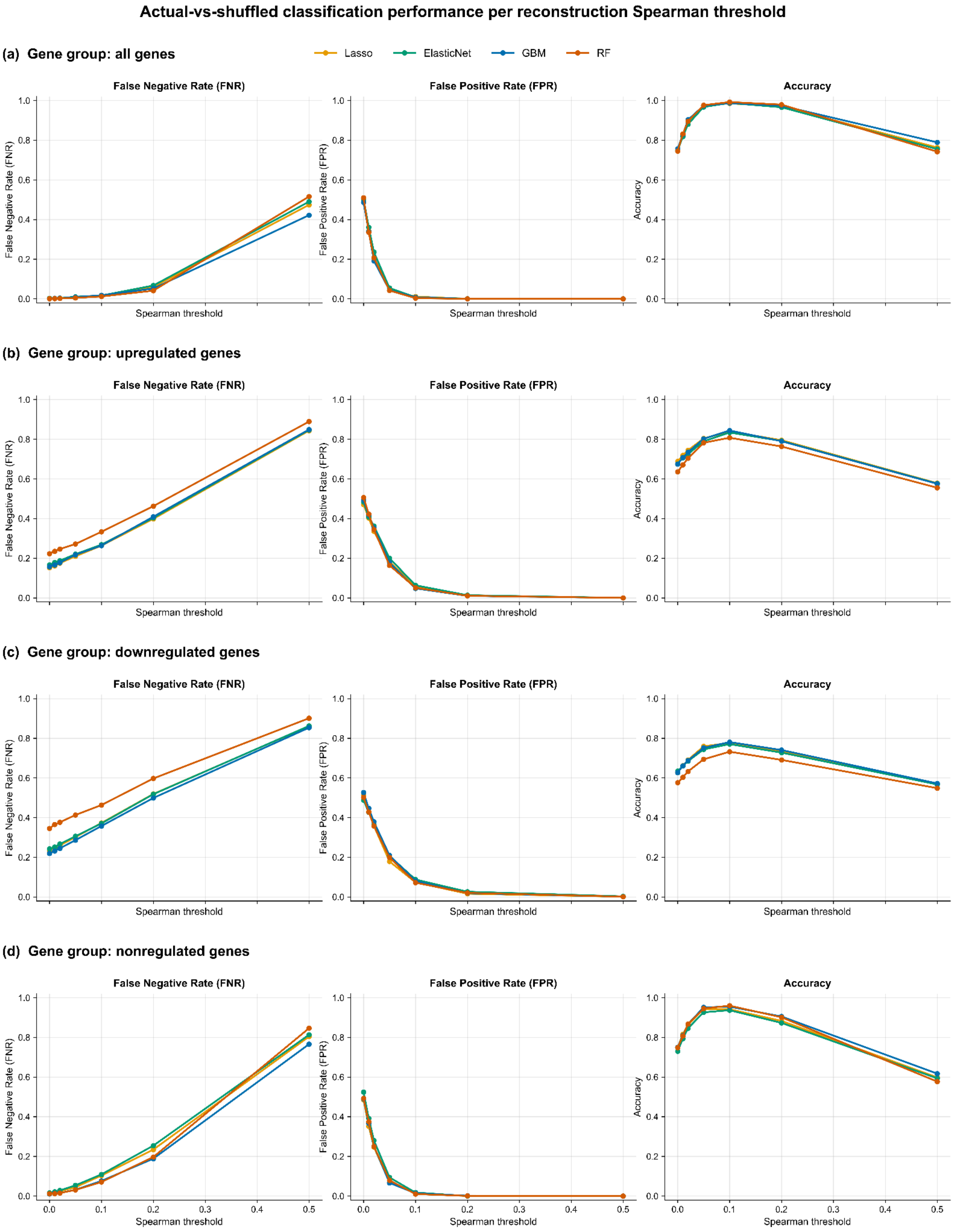
Actual-vs-shuffled classification performance per reconstruction Spearman threshold
(a) Gene group: all genes
Lasso
ElasticNet
GBM
RF
False Negative Rate (FNR)
False Positive Rate (FPR)
Accuracy
Spearman threshold
(b) Gene group: upregulated genes
(c) Gene group: downregulated genes
(d) Gene group: nonregulated genes
1.0
0.8
0.6
0.4
0.2
0.0
0.0
0.1
0.2
0.3
0.4
0.5

Figure S4. Sensitivity analysis of reconstruction Spearman thresholds for distinguishing real from shuffled model performance.

False negative rate (FNR), false positive rate (FPR), and classification accuracy across reconstruction Spearman correlation thresholds for **(a)** all genes, **(b)** upregulated genes, **(c)** downregulated genes, and **(d)** nonregulated genes. Results are shown for four regression methods: Lasso (yellow), ElasticNet (green), GBM (blue), and random forest (RF; orange).

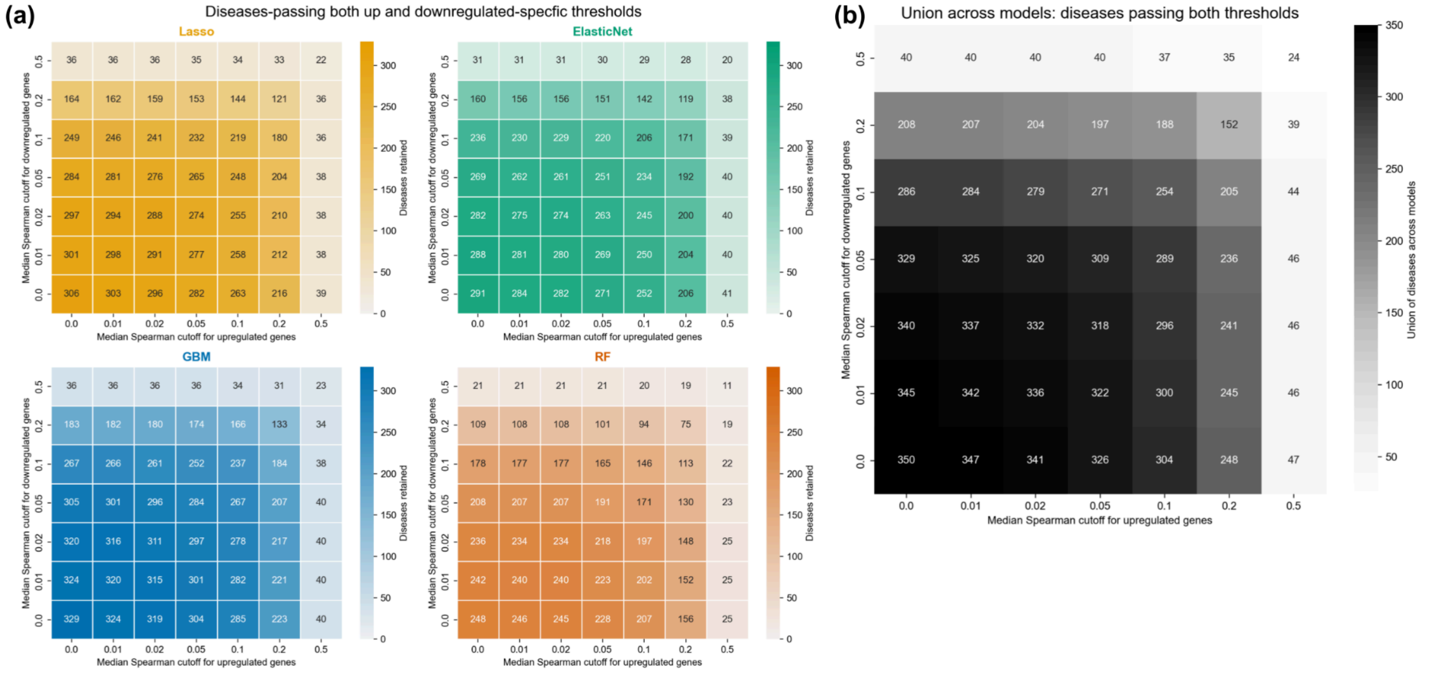


Figure S5. Sensitivity analysis of reconstruction Spearman thresholds based on the number of target diseases passing the evaluated thresholds.

Number of diseases passing both upregulated and downregulated gene median Spearman correlation thresholds across different cutoff combinations. **(a)** Number of diseases passing both thresholds for each regression method: Lasso, ElasticNet, Gradient Boosting Machine (GBM), and random forest (RF). **(b)** Number of unique diseases passing both thresholds in at least one of the four regression methods. Rows represent downregulated gene median Spearman correlation cutoffs, and columns represent upregulated gene median Spearman correlation cutoffs. Cell values indicate the number of diseases passing both thresholds.

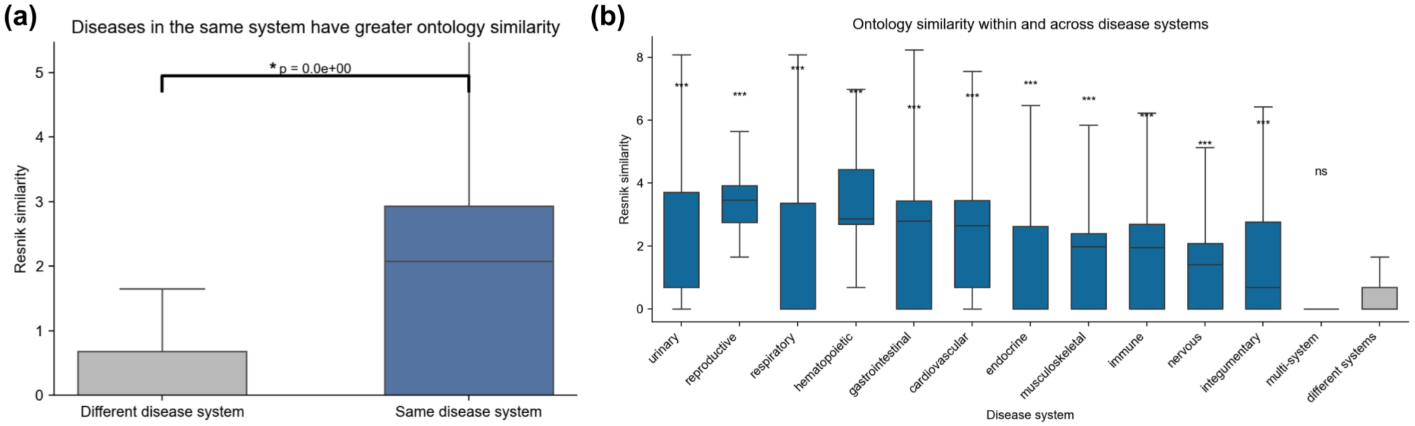


Figure S6. Diseases from the same system are significantly more related in the disease ontology compared to different systems.

**(a)** Distributions of Resnik similarity scores between disease pairs from the same system and those from different systems. **(b)** Distributions of Resnik similarity scores for disease pairs within each distinct disease system compared with the distribution for disease pairs from different systems. For both panels, statistical significance was assessed using a one-sided Mann-Whitney U test with the alternative hypothesis that same-system disease pairs had higher Resnik similarity scores than different-system disease pairs. Statistical significance was indicated as p-value < 0.05 (*), p-value < 0.01 (**), and p-value < 0.001 (***). ns indicates not statistically significant.

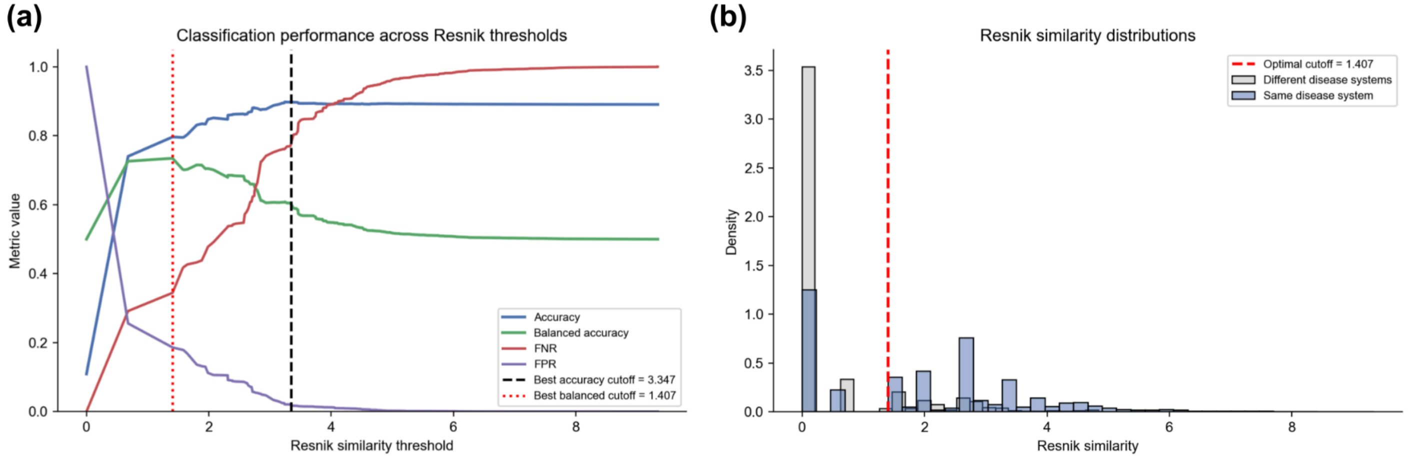


Figure S7. Sensitivity analysis to determine the optimal Resnik similarity threshold for distinguishing disease pairs from the same and different disease systems.

**(a)** Classification performance across Resnik similarity thresholds, including accuracy, balanced accuracy, false negative rate (FNR), and false positive rate (FPR). Disease pairs from the same system were considered positives, while pairs from different systems were considered negatives. The optimal threshold based on balanced accuracy red dotted line, compared with based on accuracy (black dashed line). **(b)** Distributions of Resnik similarity scores for disease pairs from the same and different disease systems. The red dashed line indicates the optimal Resnik similarity threshold selected by maximizing balanced accuracy.

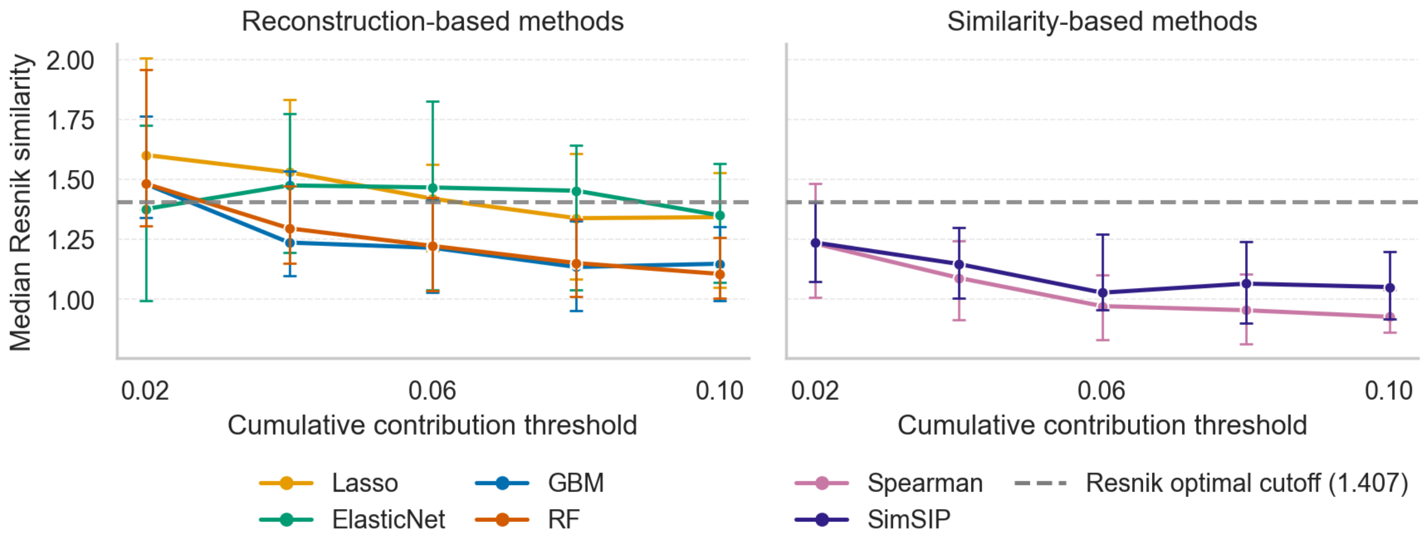


Figure S8. Median Resnik similarity among disease relationships at lower cumulative contribution thresholds.

Median of mean Resnik similarity, based on the MONDO disease ontology, among disease relationships included up to each cumulative contribution threshold (0.02, 0.04, ..., 0.1). For each target disease, disease relationships were ranked by their contribution to its prediction and included until their cumulative contribution reached the specified threshold. Higher median Resnik similarity indicates that prioritized disease pairs are more closely related within the ontology based on their most informative common ancestor. A Resnik similarity of zero indicates that the disease pair shares no informative common ancestor in the ontology. The optimal Resnik cutoff indicates the Resnik score that best separates the same-system diseases from those from different systems.

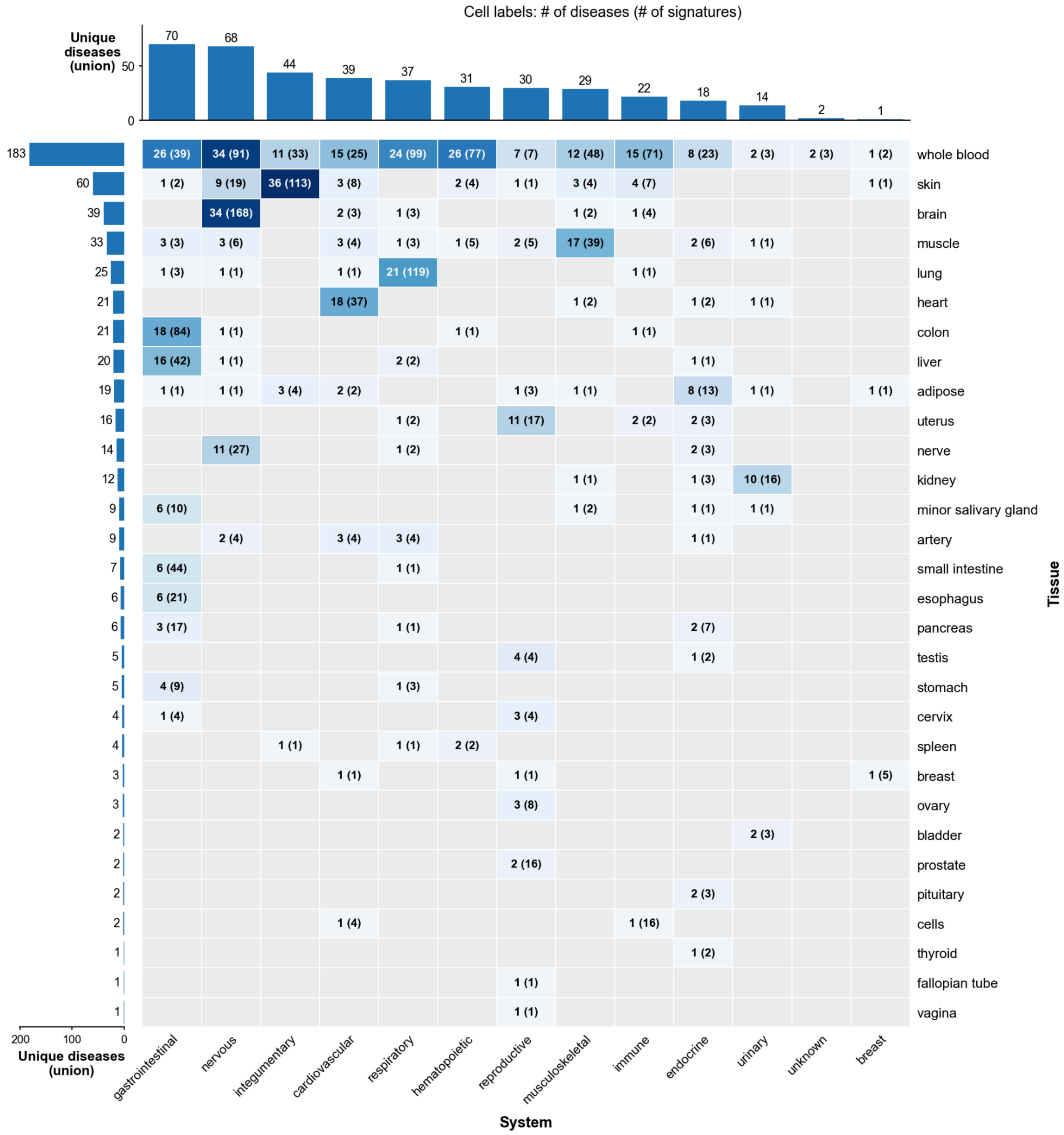


Figure S9. Distribution of disease signatures across tissues and disease systems in the full dataset.

Heatmap showing the number of unique diseases represented across tissue and disease system combinations of samples with known tissue labels from the full, unfiltered dataset. Cell shading indicates the number of unique diseases in each tissue-system combination, and cell labels report the number of unique diseases followed by the total number of signatures (diseases–signatures). Marginal bar plots show the union of unique diseases represented in each tissue (left) and disease system (top), counting each disease once within each marginal regardless of the number of signatures or tissues represented.

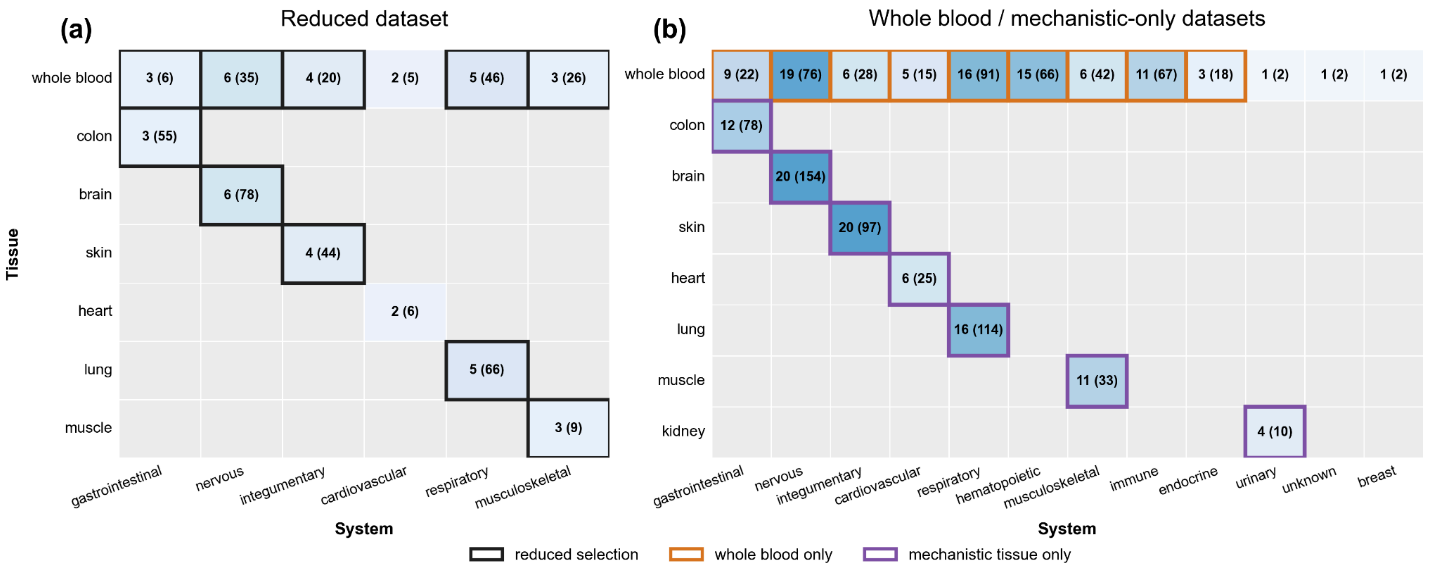


Figure S10. Selection of tissue-disease pairs for the reduced, whole-blood-only, and mechanistic-tissue-only datasets.

Heatmaps showing disease and signature counts across whole blood and corresponding mechanistic tissues for (a) the reduced dataset, requiring at least two signatures in both tissues, and (b) the whole-blood-only and mechanistic-tissue-only datasets, requiring at least two signatures in the respective tissue. Cell shading indicates the number of unique diseases, and cell labels report the number of diseases followed by the total number of signatures. Bold black outlines in (a) identify tissue_system combinations retained in the reduced dataset, while orange and purple outlines in (b) identify those retained in the whole-blood-only and mechanistic-tissue-only datasets, respectively. Disease systems with fewer than three eligible diseases were excluded from each dataset. Rows and columns with no eligible diseases are not displayed.

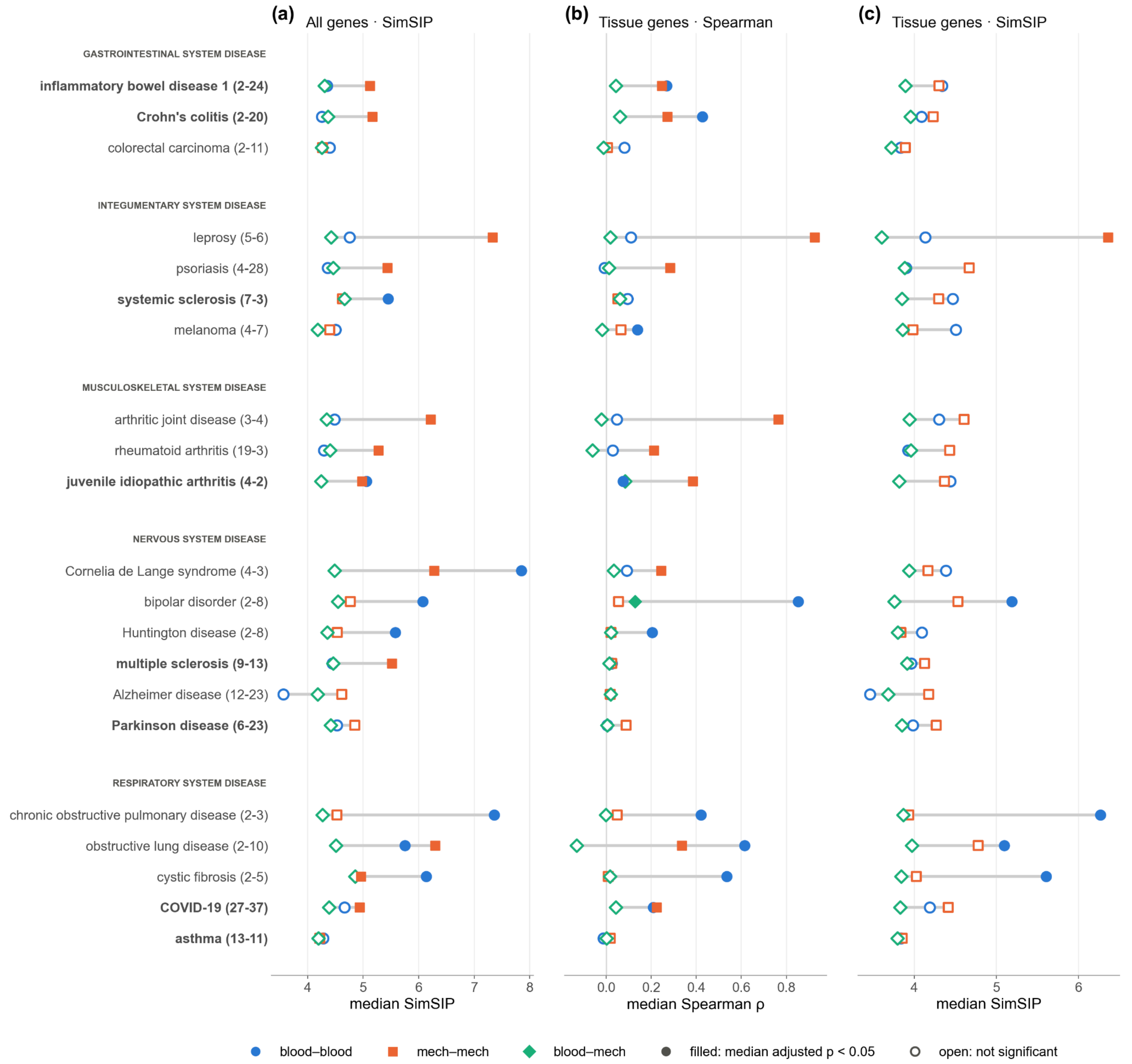


Figure S11. Within-disease agreement across whole-blood and mechanistic-tissue signatures using all genes and tissue-associated genes.

Within-disease concordance of signature pairs by tissue (**Methods 2.6**) pairing, assessed using (a) SimSIP on all genes and (b) Spearman and (c) SimSIP on GTEx tissue-associated genes (**Methods 2.8.1; Figure S3**). Each row represents a disease, grouped by disease system. Diseases meeting the reconstruction performance thresholds and having at least two whole-blood and two mechanistic-tissue signatures are indicated in bold (**Figure 5a,b**). Numbers in parentheses indicate the counts of whole-blood and mechanistic-tissue signatures. Markers show median similarity across blood-blood (circle), mech-mech (square), and blood-mech (diamond) signature pairs; gray bars span the range of the three medians. Filled markers indicate a median Bonferroni-adjusted p-value ($P$) < 0.05; open markers indicate nonsignificance. Chance-level SimSIP is approximately 4.

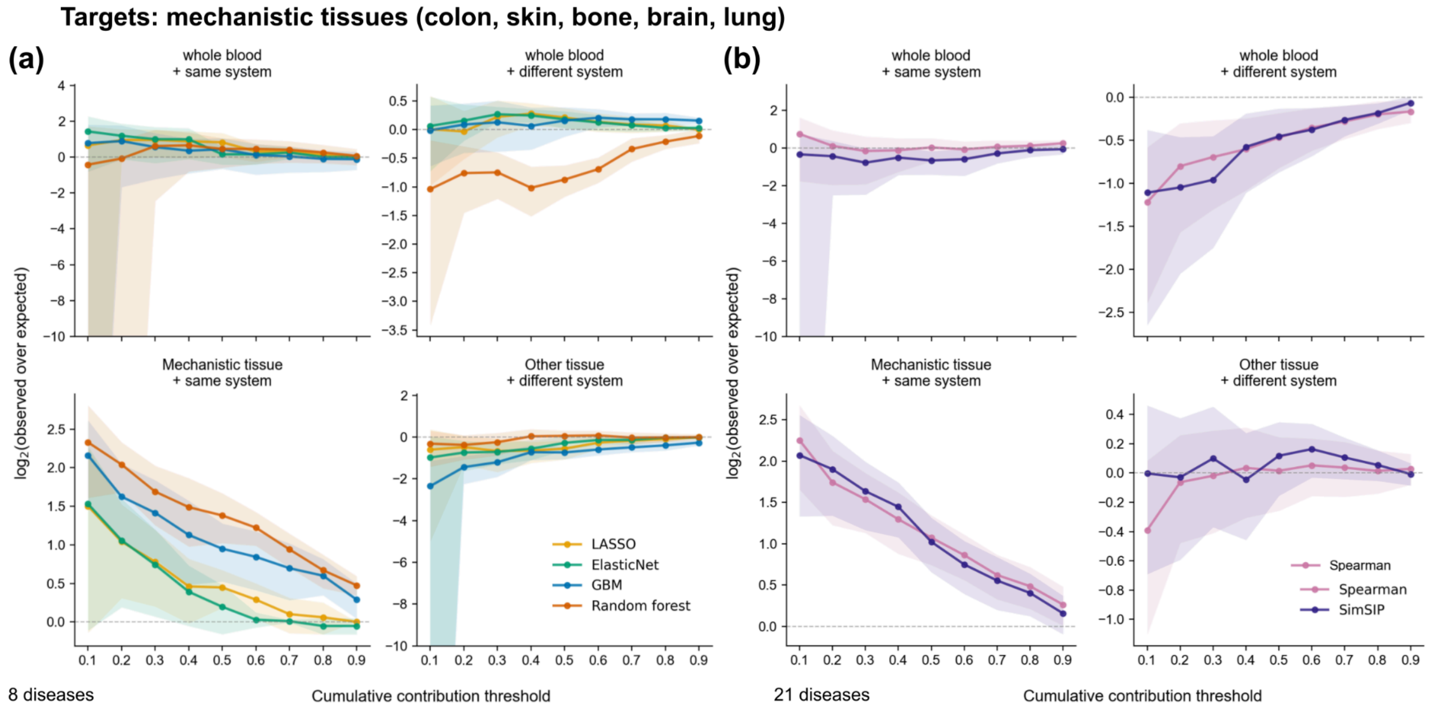


Figure S12. Mechanistic tissue-disease targets prioritize mechanistic tissue-disease features.

Enrichment of whole-blood or mechanistic tissue/system relationship categories among the disease relationships cumulatively contributes to different contribution thresholds for each **(a)** reconstruction-based method and **(b)** similarity-based method: 0.1, 0.2, ..., 0.9 given mechanistic-tissue (colon, skin, muscle, brain, or lung) target disease signatures. The y-axis shows the average $\log_2$(fold enrichment), where fold enrichment is defined as the proportion of prioritized disease signatures belonging to a given tissue/system relationship category divided by the corresponding proportion among all disease signatures at each cumulative contribution threshold. A positive value indicates that the relationship is overrepresented relative to the background expectation, whereas a value of zero indicates no enrichment (as expected by chance), and a negative value indicates underrepresentation relative to the background.

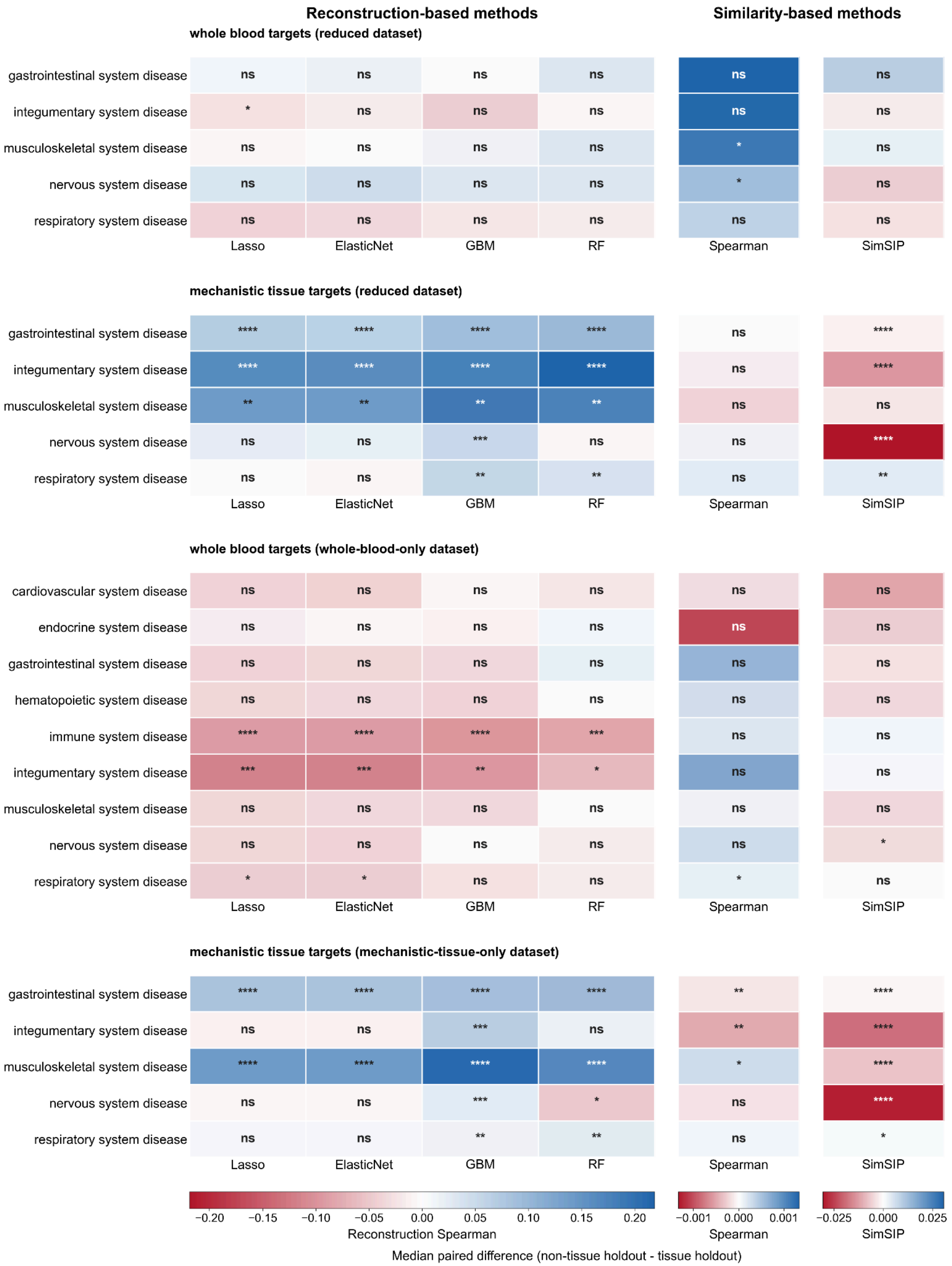

Reconstruction-based methods
Similarity-based methods
whole blood targets (reduced dataset)
gastrointestinal system disease
integumentary system disease
musculoskeletal system disease
nervous system disease
respiratory system disease
ns ns ns ns ns ns
* ns ns ns ns ns
ns ns ns ns * ns
ns ns ns ns * ns
ns ns ns ns ns ns
Lasso
ElasticNet
GBM
RF
Spearman
SimSIP
mechanistic tissue targets (reduced dataset)
gastrointestinal system disease
integumentary system disease
musculoskeletal system disease
nervous system disease
respiratory system disease
**** **** **** **** ns ****
**** **** **** **** ns ****
** ** ** ** ns ns
ns ns *** ns ns ****
ns ns ** ** ns **
Lasso
ElasticNet
GBM
RF
Spearman
SimSIP
whole blood targets (whole-blood-only dataset)
cardiovascular system disease
endocrine system disease
gastrointestinal system disease
hematopoietic system disease
immune system disease
integumentary system disease
musculoskeletal system disease
nervous system disease
respiratory system disease
ns ns ns ns ns ns
ns ns ns ns ns ns
ns ns ns ns ns ns
ns ns ns ns ns ns
**** **** **** *** ns ns
*** *** ** * ns ns
ns ns ns ns ns ns
ns ns ns ns ns *
* * ns ns * ns
Lasso
ElasticNet
GBM
RF
Spearman
SimSIP
mechanistic tissue targets (mechanistic-tissue-only dataset)
gastrointestinal system disease
integumentary system disease
musculoskeletal system disease
nervous system disease
respiratory system disease
**** **** **** **** ** ****
ns ns *** ns ** ****
**** **** **** **** * ****
ns ns *** * ns ****
ns ns ** ** ns *
Lasso
ElasticNet
GBM
RF
Spearman
SimSIP
−0.20 −0.15 −0.10 −0.05 0.00 0.05 0.10 0.15 0.20
Reconstruction Spearman
−0.001 0.000 0.001
Spearman
−0.025 0.000 0.025
SimSIP
Median paired difference (non-tissue holdout - tissue holdout)

#### Figure S13. Effects of tissue-specific holdout on reconstruction performance and similarity scores across disease systems.

Heatmaps showing the median paired differences between non-tissue and tissue holdout conditions for reconstruction-based methods: Lasso, ElasticNet, GBM, and RF, and similarity-based methods: Spearman and SimSIP, stratified by disease system. Results are shown for **(a)** whole-blood target diseases using the reduced feature matrix, **(b)** mechanistic-tissue target diseases using the reduced feature matrix, **(c)** whole-blood target diseases using the whole-blood-only feature matrix, and **(d)** mechanistic-tissue target diseases using the mechanistic-tissue-only feature matrix. For each method and disease system, the median paired difference was calculated across target diseases as the non-tissue holdout value minus the corresponding tissue holdout value. Reconstruction-based methods were evaluated using test reconstruction Spearman, while similarity-based methods were evaluated using their respective similarity scores. Positive values indicate higher performance or similarity scores under non-tissue holdout, whereas negative values indicate higher values under tissue holdout. Statistical significance was assessed using two-sided paired Wilcoxon signed-rank tests. Significance was indicated as p-value < 0.05 (*), p-value < 0.01 (**), p-value < 0.001 (***), and p-value < 0.0001 (****); ns indicates not statistically significant.

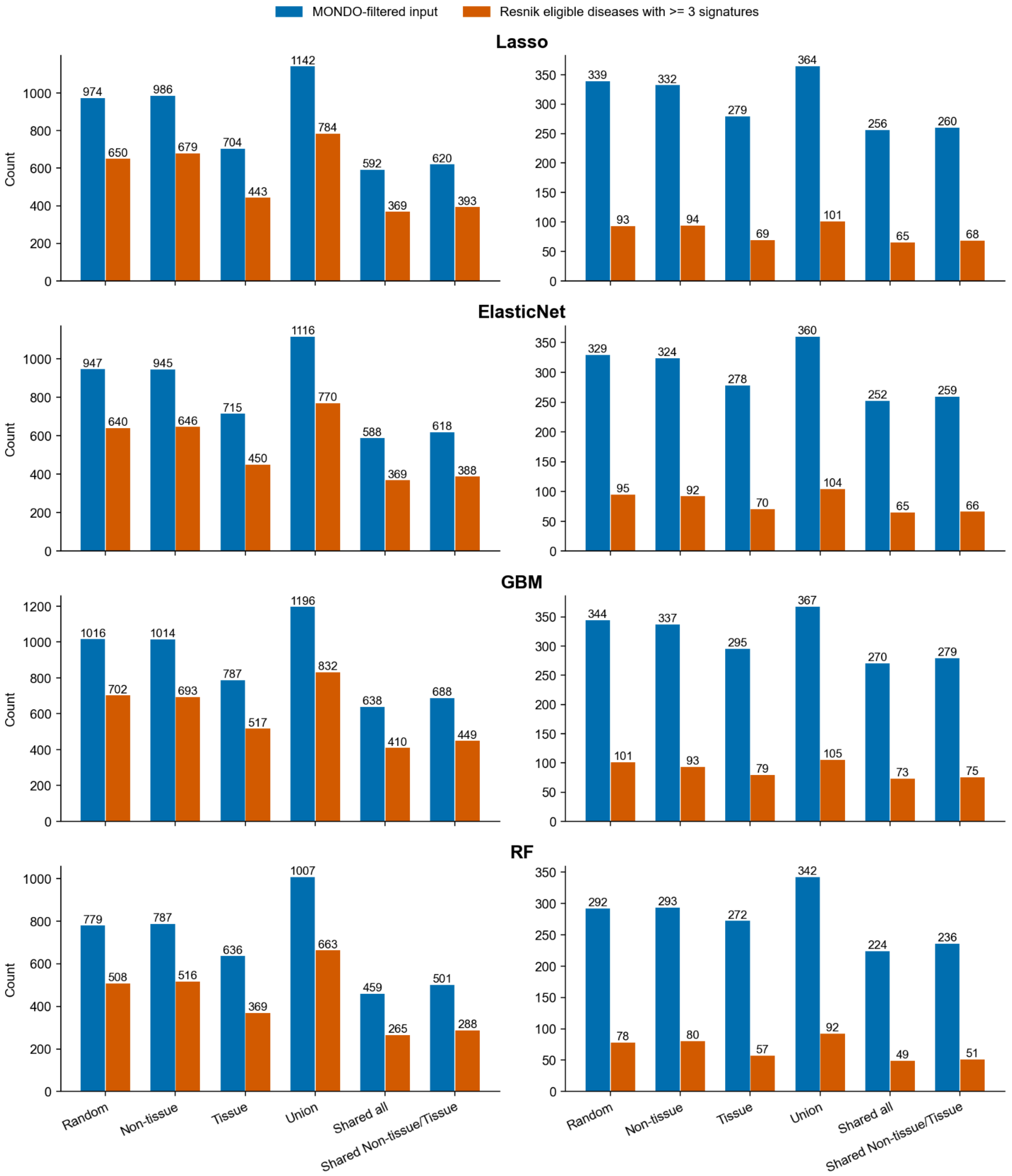
MONDO-filtered input
Resnik eligible diseases with >= 3 signatures
Lasso
ElasticNet
GBM
RF
Count
Random
Non-tissue
Tissue
Union
Shared all
Shared Non-tissue/Tissue

Figure S14. Number of target signatures and diseases retained across tissue holdout settings and filtering steps using the full disease dataset.

Number of target signatures (left column) and unique target diseases (right column) passing the performance filter for four reconstruction-based methods: Lasso, ElasticNet, GBM, and RF, shown from top to bottom. Results are presented across six tissue holdout settings: Random, Non-tissue, Tissue, Union, Shared all, and Shared Non-tissue/Tissue. Bars represent the number of input targets (blue), targets remaining after MONDO filtering (orange), eligible targets associated with at least three signatures (green), and final valid Resnik targets (red). Values above the bars indicate the corresponding counts at each filtering step. These target signatures were used in the analyses presented in **Figure S15**.

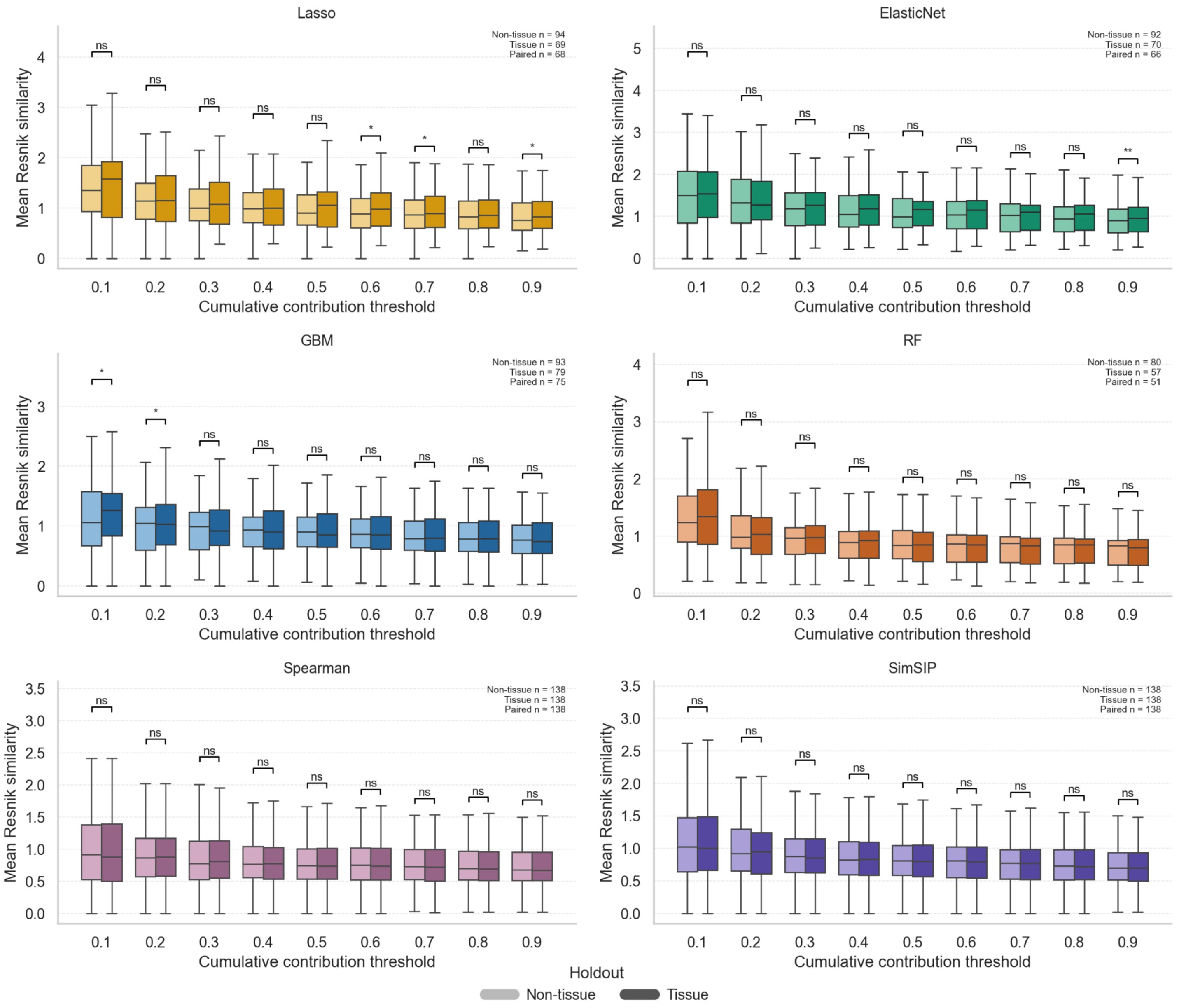

Figure S15. Comparison of mean Resnik similarity between tissue and non-tissue holdout settings across cumulative contribution thresholds using the full dataset.

Distributions of mean Resnik similarity between target diseases and their prioritized feature diseases across cumulative contribution thresholds (0.1 to 0.9) for four reconstruction-based methods: Lasso, ElasticNet, GBM, and RF, and two similarity-based methods: Spearman and SimSIP. For each target disease, mean Resnik similarity was calculated across the feature diseases retained at each cumulative contribution threshold. Boxplots show the distributions of these target-level means for tissue and non-tissue holdout settings, restricted to target diseases shared between the two settings. Boxes represent the interquartile range, with the central line indicating the median. Comparisons between holdout settings were performed using two-sided paired Wilcoxon signed-rank tests, with target diseases as matched pairs. P-values were adjusted using the Benjamini-Hochberg method separately for each method across the nine thresholds. Statistical significance is indicated as adjusted p-value (*P*) < 0.05 (*), *P* < 0.01 (**), *P* < 0.001 (***), and *P* < 0.0001 (****); ns indicates not statistically significant. Sample sizes indicate the number of target diseases in each holdout setting and the number of matched target diseases.

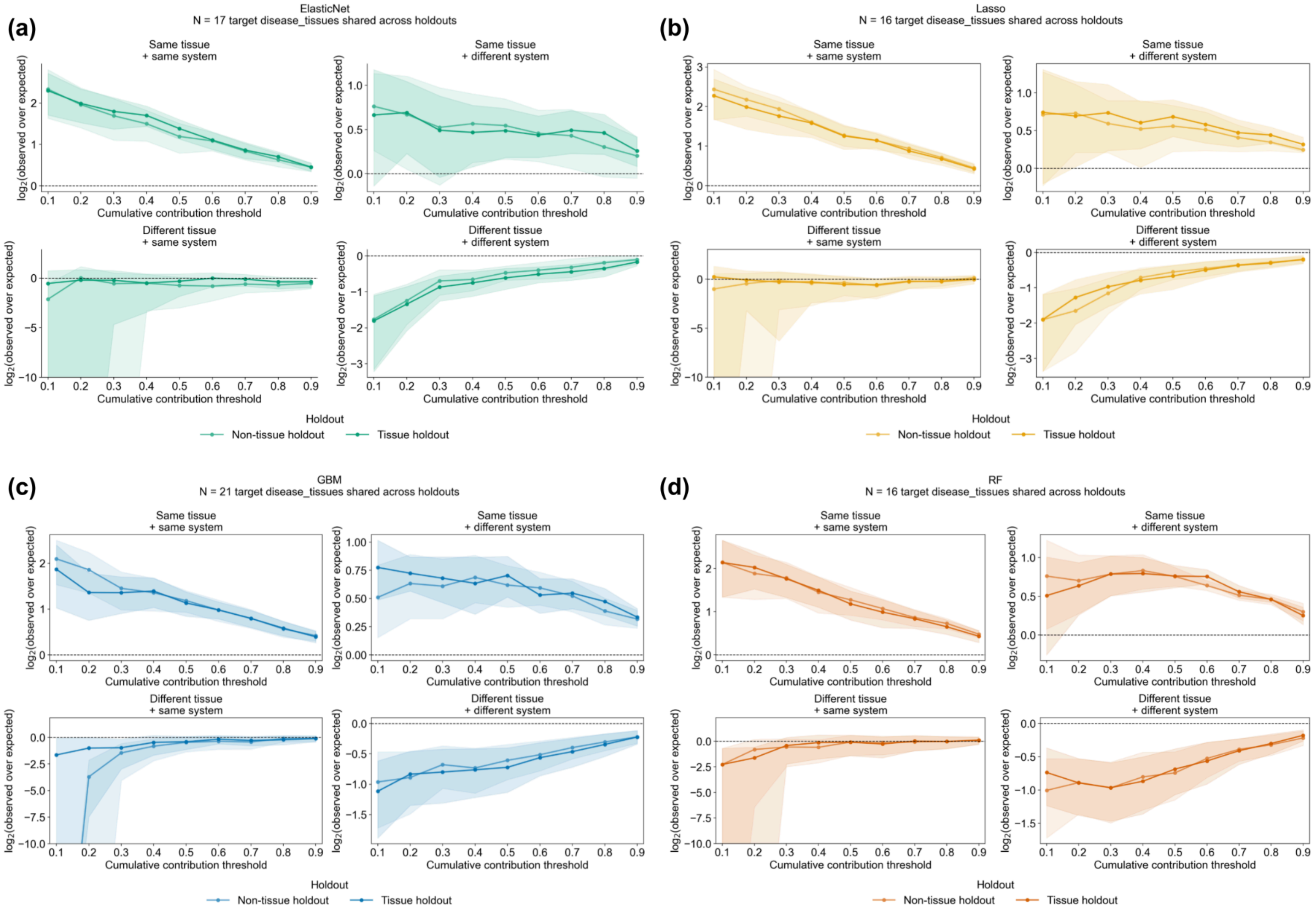


Figure S16. Enrichment of tissue and disease system relationships among prioritized disease features of reconstruction-based methods across cumulative contribution thresholds using reduced datasets.

Enrichment of tissue and disease system relationship categories among disease features cumulatively contributing to thresholds ranging from 0.1 to 0.9, in increments of 0.1, for **(a)** ElasticNet, **(b)** Lasso, **(c)** GBM, and **(d)** RF. Analyses combined whole-blood and mechanistic-tissue (colon, skin, muscle, brain, or lung) target disease signatures using reduced datasets under non-tissue and tissue holdout settings. Disease relationships were classified into four categories based on whether the target and feature diseases shared the same or different tissue and disease system: same tissue and same system, same tissue and different system, different tissue and same system, or different tissue and different system. The y-axis shows the log2(average fold enrichment), where fold enrichment was defined as the proportion of prioritized disease features belonging to a given tissue/system relationship category divided by the corresponding proportion among all available disease features. Positive values indicate overrepresentation relative to the background, zero indicates no enrichment, and negative values indicate underrepresentation. Lines compare non-tissue and tissue holdout settings, with shaded regions indicating variability across target disease_tissue combinations. The number of target disease_tissue combinations shared between holdout settings is indicated above each method.

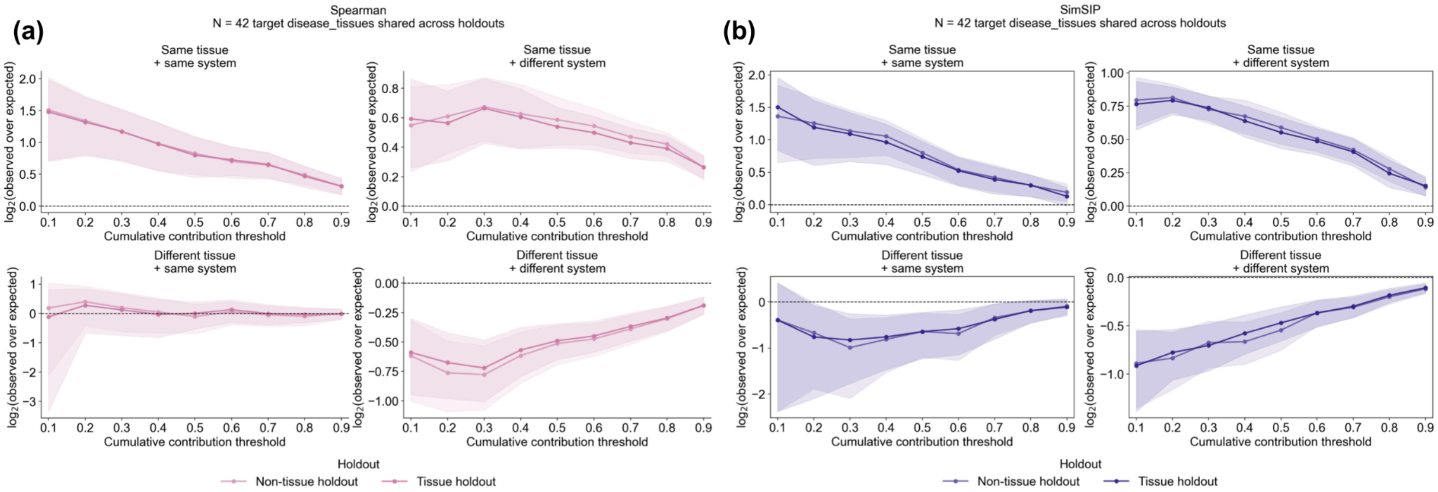


Figure S17. Enrichment of tissue and disease system relationships among prioritized disease features of similarity-based methods across cumulative contribution thresholds using reduced datasets.

Enrichment of tissue and disease system relationship categories among disease features cumulatively contributing to thresholds ranging from 0.1 to 0.9, in increments of 0.1, for **(a)** Spearman, **(b)** SimSIP. All other details are as described in **Figure S16**.

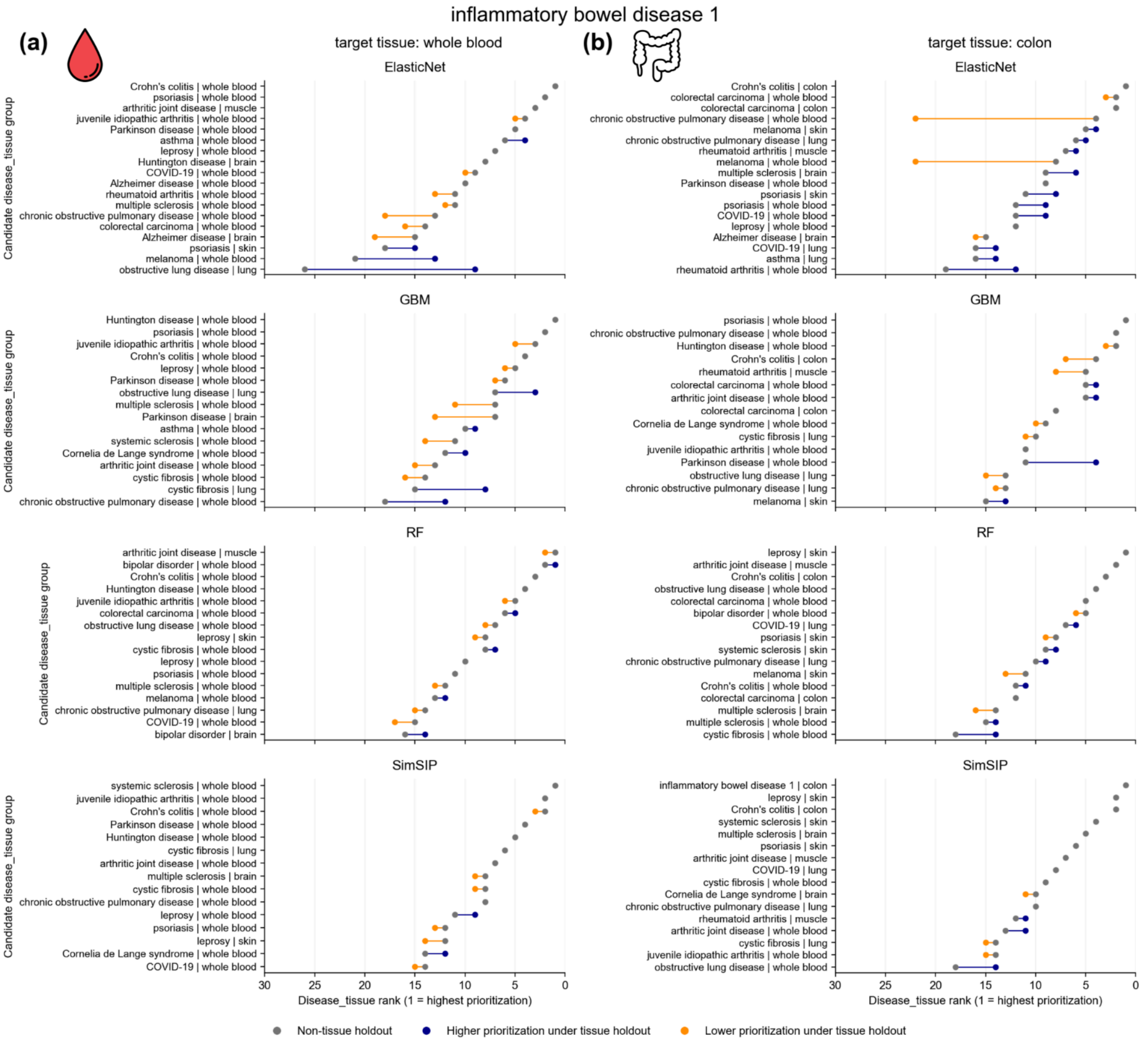


Figure S18. Tissue holdouts altered the prioritization ranks of feature disease_tissue predictions compared to non-tissue holdouts for IBD1 targets derived from whole blood and colon.

Rank shifts of feature disease_tissue predictions between non-tissue and tissue holdouts for inflammatory bowel disease 1 (IBD1) targets derived from whole blood (left) and colon (right), using reconstruction-based methods (ElasticNet, GBM, and RF) and the similarity-based method SimSIP. disease_tissue features were ranked according to their median rank percentiles under each holdout condition (non-tissue holdout: gray dots; tissue holdout: colored dots). Features ranked among the top 15 under either holdout condition were included, resulting in varying numbers of displayed features depending on the overlap between the two sets. Connecting lines illustrate changes in prioritization ranks between holdout conditions. Lower numerical ranks indicate higher prioritization. Red indicates decreased prioritization, whereas green indicates increased prioritization following the removal of tissue-specific genes.

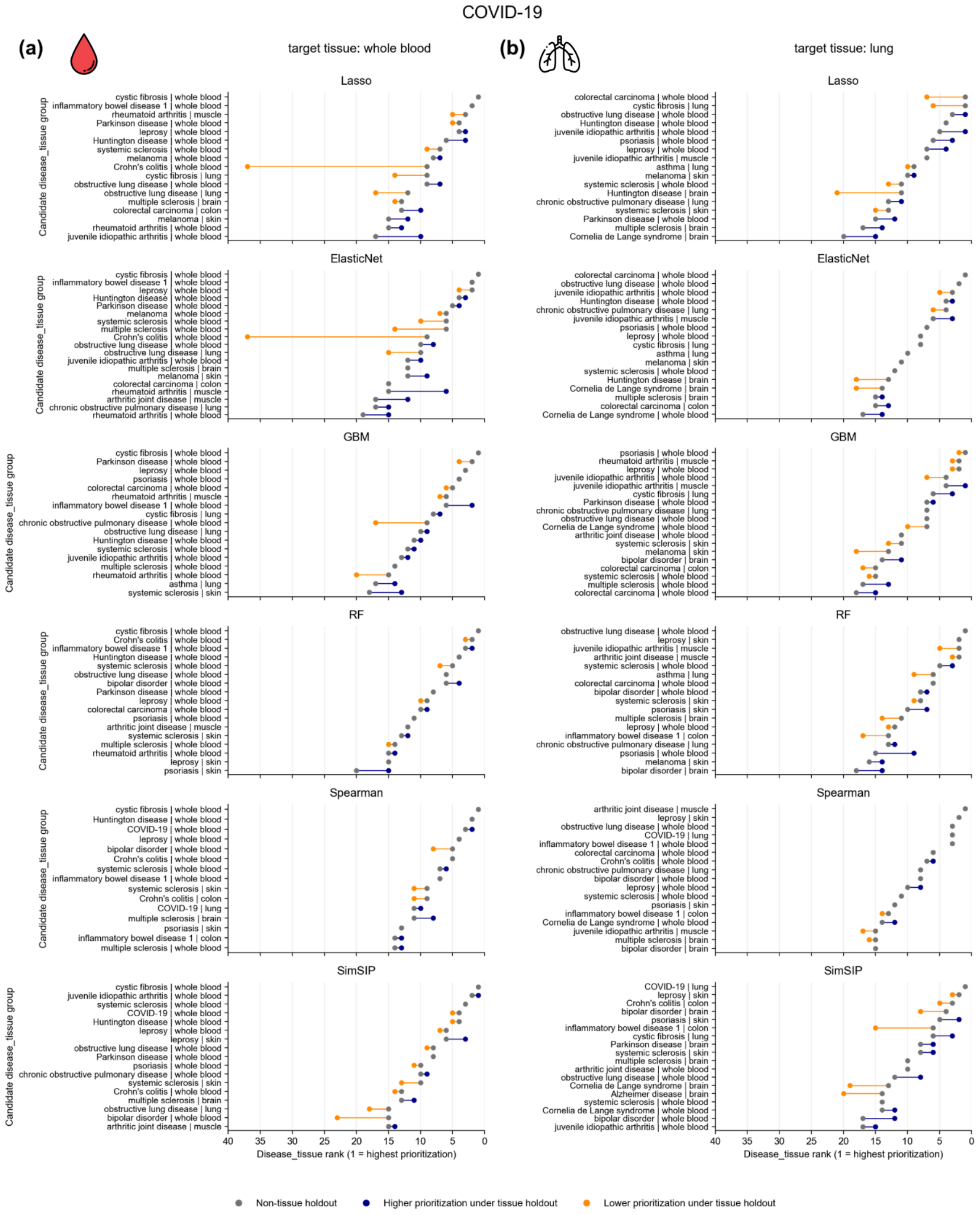
COVID-19
(a)
target tissue: whole blood
(b)
target tissue: lung
Lasso
ElasticNet
GBM
RF
Spearman
SimSIP
Candidate disease_tissue group
Disease_tissue rank (1 = highest prioritization)
40 35 30 25 20 15 10 5 0
cystic fibrosis | whole blood
inflammatory bowel disease 1 | whole blood
rheumatoid arthritis | muscle
Parkinson disease | whole blood
leprosy | whole blood
Huntington disease | whole blood
systemic sclerosis | whole blood
melanoma | whole blood
Crohn's colitis | whole blood
cystic fibrosis | lung
obstructive lung disease | whole blood
obstructive lung disease | lung
multiple sclerosis | brain
colorectal carcinoma | colon
melanoma | skin
rheumatoid arthritis | whole blood
juvenile idiopathic arthritis | whole blood
colorectal carcinoma | whole blood
psoriasis | whole blood
juvenile idiopathic arthritis | muscle
asthma | lung
systemic sclerosis | skin
Huntington disease | brain
chronic obstructive pulmonary disease | lung
Cornelia de Lange syndrome | brain
arthritic joint disease | muscle
Cornelia de Lange syndrome | whole blood
bipolar disorder | whole blood
bipolar disorder | brain
inflammatory bowel disease 1 | colon
psoriasis | skin
leprosy | skin
COVID-19 | whole blood
COVID-19 | lung
Crohn's colitis | colon
arthritic joint disease | whole blood
multiple sclerosis | whole blood
Alzheimer disease | brain
Non-tissue holdout
Higher prioritization under tissue holdout
Lower prioritization under tissue holdout

Figure S19. Tissue holdouts altered the prioritization ranks of feature disease_tissue predictions compared to non-tissue holdouts for COVID-19 targets derived from whole blood and lung.

Rank shifts of feature disease_tissue predictions between non-tissue and tissue holdouts for COVID-19 target signatures derived from (a) whole blood and (b) lung, using reconstruction-based methods (Lasso, ElasticNet, GBM, and RF) and similarity-based methods (Spearman and SimSIP). Features were ranked by their median rank percentiles under each holdout condition (non-tissue: gray dots; tissue: colored dots). Features ranked among the top 15 under either condition were included, with connecting lines indicating changes in prioritization. Lower numerical ranks indicate higher prioritization. Red indicates decreased prioritization, whereas green indicates increased prioritization following the removal of tissue-specific genes.

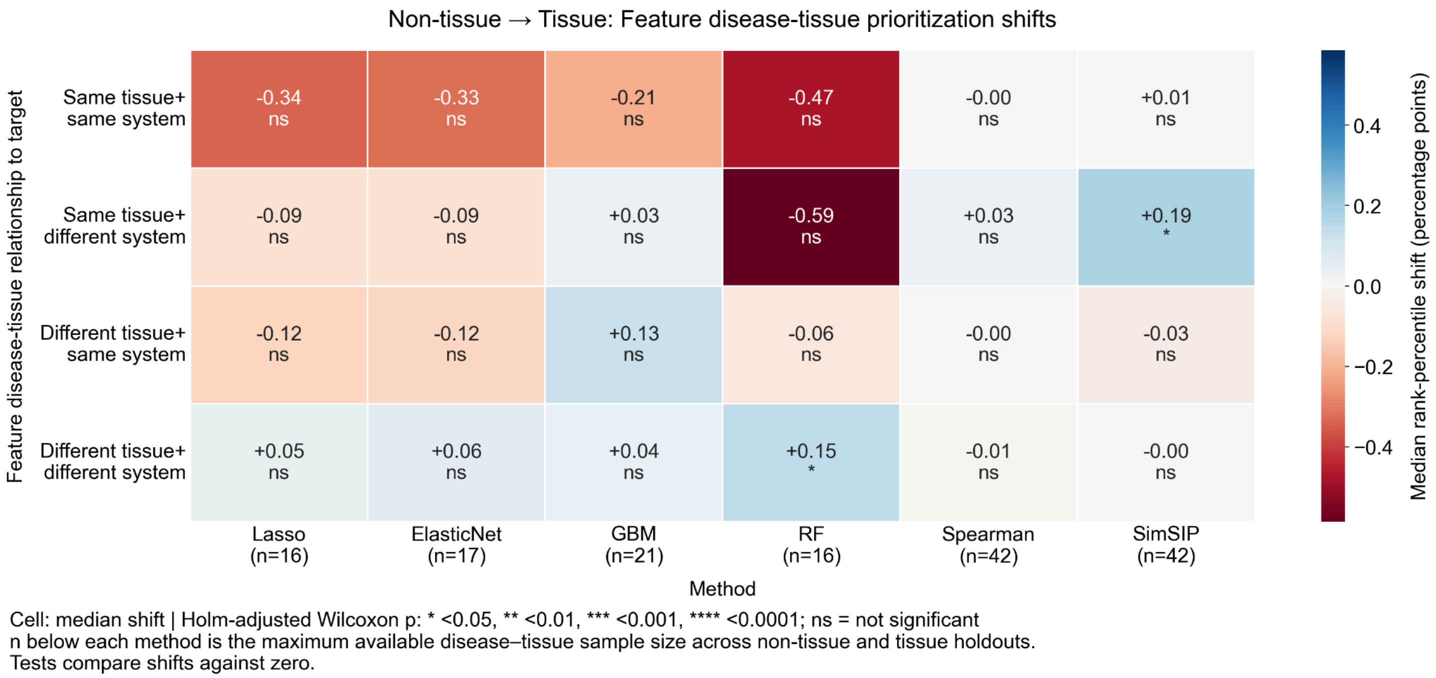


Figure S20. Shifts in feature disease_tissue prioritization between non-tissue and tissue holdouts across tissue and disease system relationship categories using reduced datasets.

Heatmap showing median rank-percentile shifts (tissue minus non-tissue holdout) across four reconstruction-based methods: Lasso, ElasticNet, GBM, and RF, and two similarity-based methods: Spearman and SimSIP. Positive values indicate decreased prioritization, whereas negative values indicate increased prioritization under tissue holdouts. Statistical significance was assessed using two-sided Wilcoxon signed-rank tests against zero, with p-values adjusted using the Holm method separately for each method across four relationship categories. Significance annotations indicate adjusted p-value (*P*) < 0.05 (*), *P* < 0.01 (**), *P* < 0.001 (***), and *P* < 0.0001 (****); ns indicates not statistically significant. Sample sizes indicate the number of disease_tissue targets present in each method.

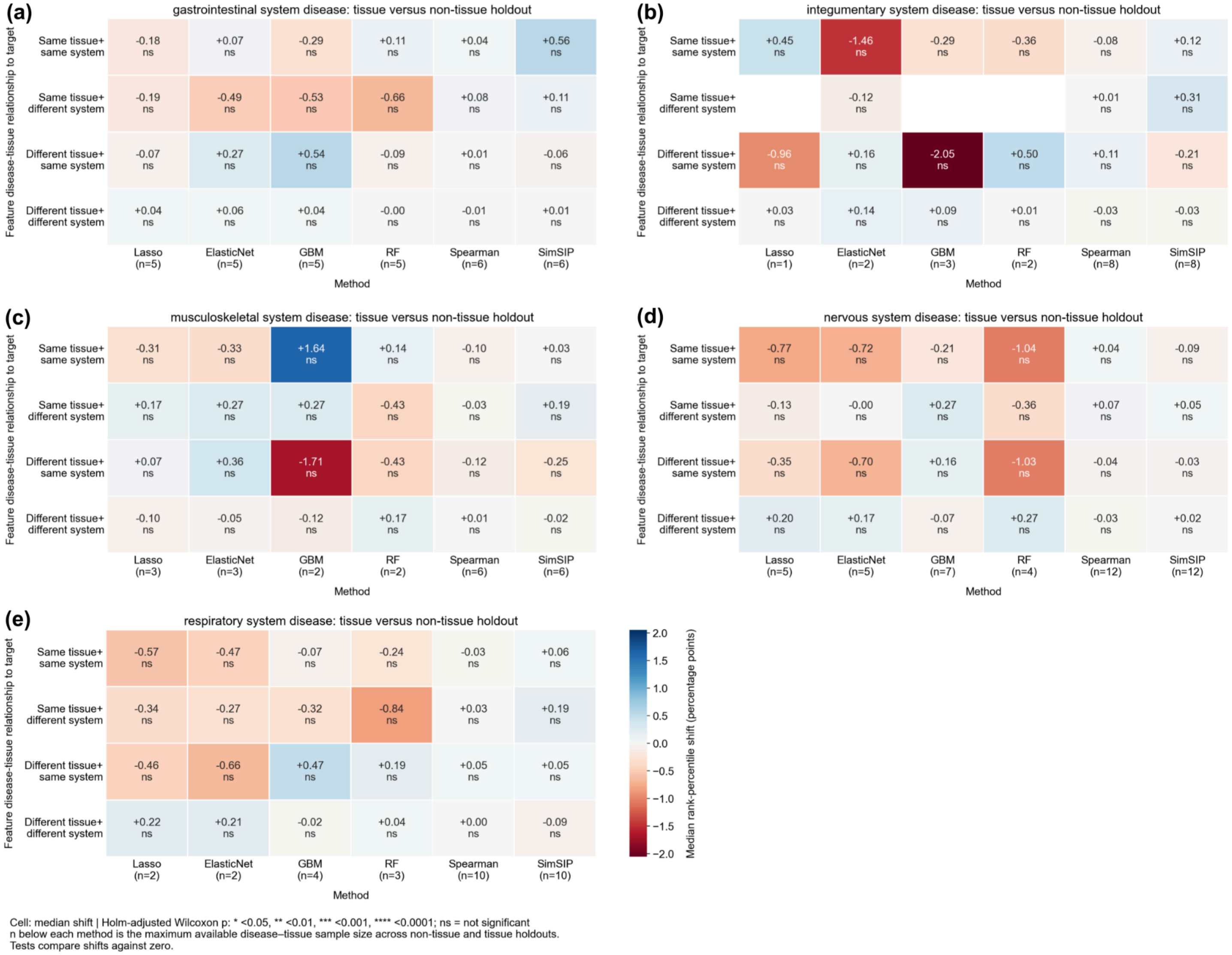


Figure S21. Shifts in feature disease_tissue prioritization between non-tissue and tissue holdouts across tissue and disease system relationship categories, stratified by target disease system, using reduced datasets.

Heatmaps showing median rank-percentile shifts (tissue minus non-tissue holdout) for target diseases belonging to the **(a)** gastrointestinal, **(b)** integumentary, **(c)** musculoskeletal, **(d)** nervous, and **(e)** respiratory systems across six prioritization methods. Positive values indicate decreased prioritization, whereas negative values indicate increased prioritization under tissue holdouts. Statistical significance was assessed using two-sided Wilcoxon signed-rank tests against zero, with p-values adjusted using the Holm method separately for each method across four relationship categories. Significance annotations indicate adjusted *p*-value (*P*) < 0.05 (*), *P < 0.01 (**), P < 0.001* (***), and *P* < 0.0001 (****); ns indicates not statistically significant. Sample sizes indicate the number of disease_tissue targets present in each method. Blank cells indicate relationship categories with no available disease_tissue features.

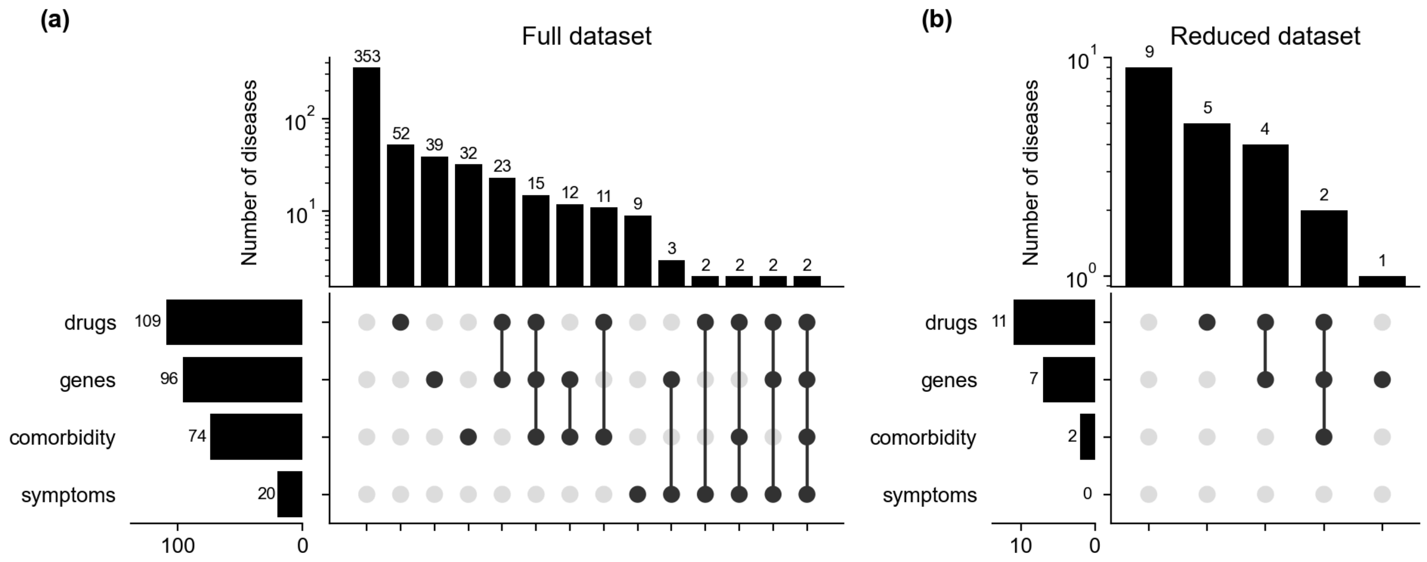


Figure S22. Comparison of biological evidence coverage between the full and reduced datasets.

UpSet plots showing the numbers of diseases supported by shared genes, drugs, symptoms, and comorbidity evidence in **(a)** the full dataset and **(b)** the reduced dataset. Horizontal bars indicate the total number of diseases associated with each evidence type, while vertical bars indicate the number of diseases in each evidence intersection. Connected black dots identify the evidence types included in each intersection.

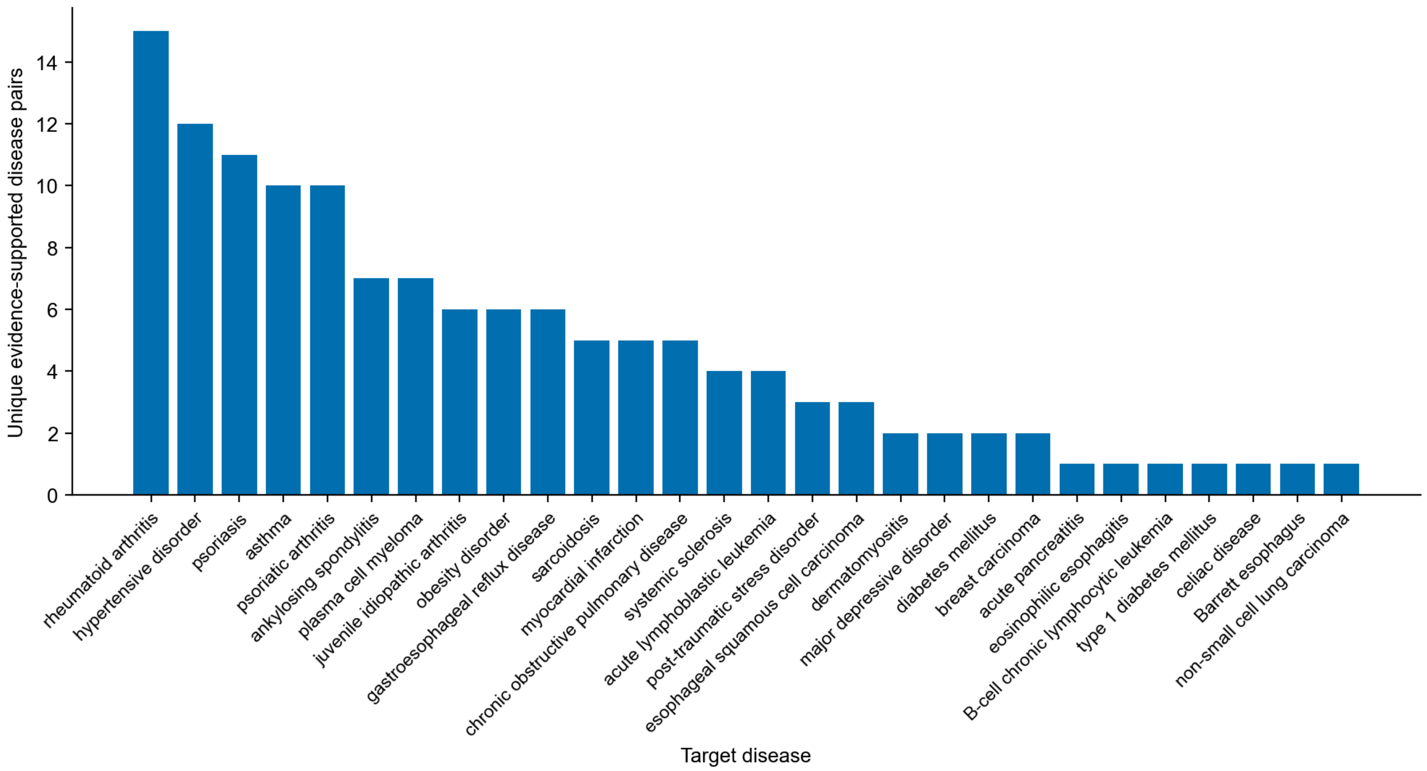


Figure S23. Distribution of positive disease pairs across target diseases supported by multiple relevance evidence types in the full dataset.

Bar plot showing the number of unique positive disease pairs for each target disease, defined as disease pairs supported by at least two of four independent evidence types: shared genes, drugs, symptoms, and comorbidity. Target diseases are ordered by decreasing numbers of positive disease pairs.

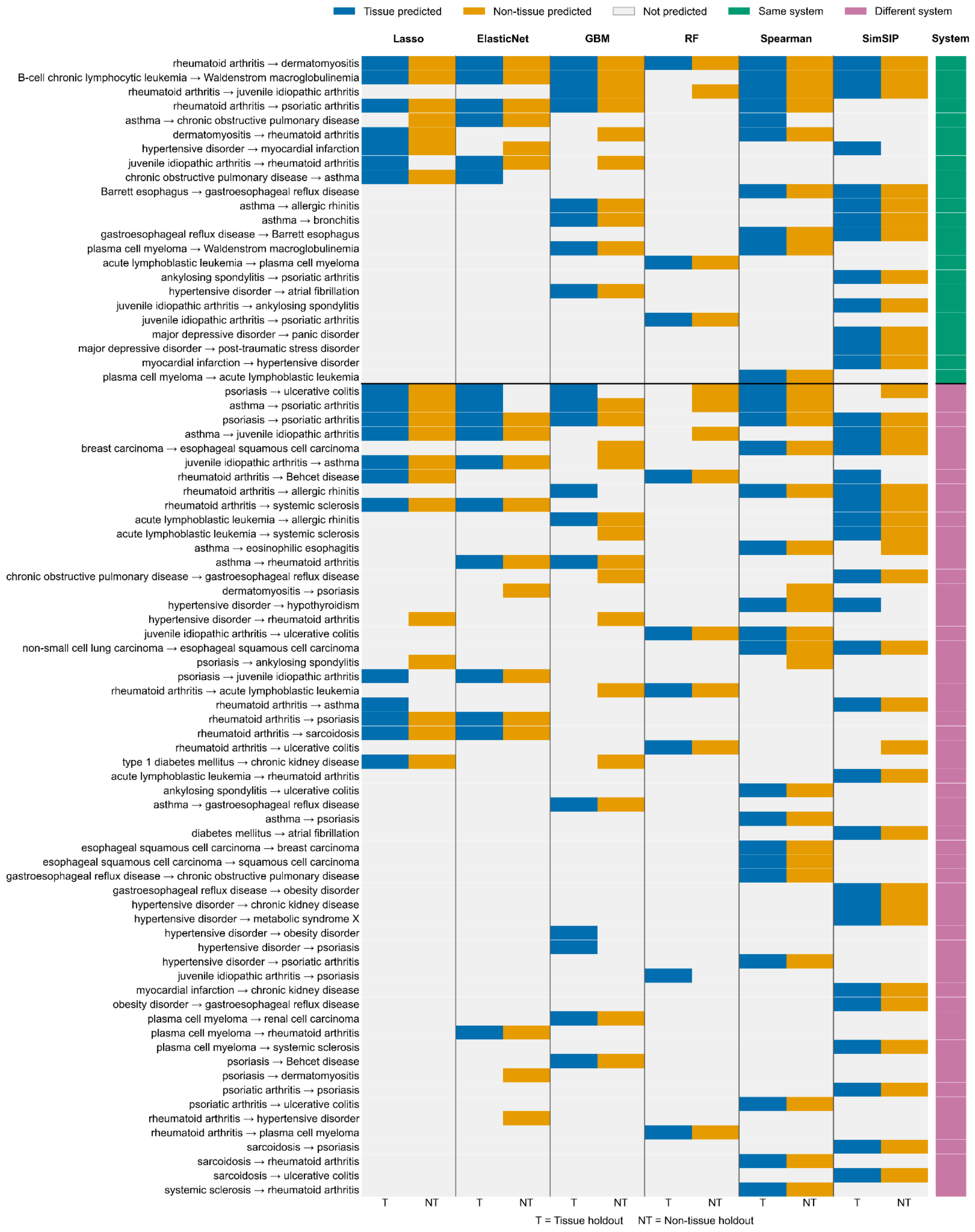
Tissue predicted
Non-tissue predicted
Not predicted
Same system
Different system
Lasso
ElasticNet
GBM
RF
Spearman
SimSIP
System
rheumatoid arthritis → dermatomyositis
B-cell chronic lymphocytic leukemia → Waldenstrom macroglobulinemia
rheumatoid arthritis → juvenile idiopathic arthritis
rheumatoid arthritis → psoriatic arthritis
asthma → chronic obstructive pulmonary disease
dermatomyositis → rheumatoid arthritis
hypertensive disorder → myocardial infarction
juvenile idiopathic arthritis → rheumatoid arthritis
chronic obstructive pulmonary disease → asthma
Barrett esophagus → gastroesophageal reflux disease
asthma → allergic rhinitis
asthma → bronchitis
gastroesophageal reflux disease → Barrett esophagus
plasma cell myeloma → Waldenstrom macroglobulinemia
acute lymphoblastic leukemia → plasma cell myeloma
ankylosing spondylitis → psoriatic arthritis
hypertensive disorder → atrial fibrillation
juvenile idiopathic arthritis → ankylosing spondylitis
juvenile idiopathic arthritis → psoriatic arthritis
major depressive disorder → panic disorder
major depressive disorder → post-traumatic stress disorder
myocardial infarction → hypertensive disorder
plasma cell myeloma → acute lymphoblastic leukemia
psoriasis → ulcerative colitis
asthma → psoriatic arthritis
psoriasis → psoriatic arthritis
asthma → juvenile idiopathic arthritis
breast carcinoma → esophageal squamous cell carcinoma
juvenile idiopathic arthritis → asthma
rheumatoid arthritis → Behcet disease
rheumatoid arthritis → allergic rhinitis
rheumatoid arthritis → systemic sclerosis
acute lymphoblastic leukemia → allergic rhinitis
acute lymphoblastic leukemia → systemic sclerosis
asthma → eosinophilic esophagitis
asthma → rheumatoid arthritis
chronic obstructive pulmonary disease → gastroesophageal reflux disease
dermatomyositis → psoriasis
hypertensive disorder → hypothyroidism
hypertensive disorder → rheumatoid arthritis
juvenile idiopathic arthritis → ulcerative colitis
non-small cell lung carcinoma → esophageal squamous cell carcinoma
psoriasis → ankylosing spondylitis
psoriasis → juvenile idiopathic arthritis
rheumatoid arthritis → acute lymphoblastic leukemia
rheumatoid arthritis → asthma
rheumatoid arthritis → psoriasis
rheumatoid arthritis → sarcoidosis
rheumatoid arthritis → ulcerative colitis
type 1 diabetes mellitus → chronic kidney disease
acute lymphoblastic leukemia → rheumatoid arthritis
ankylosing spondylitis → ulcerative colitis
asthma → gastroesophageal reflux disease
asthma → psoriasis
diabetes mellitus → atrial fibrillation
esophageal squamous cell carcinoma → breast carcinoma
esophageal squamous cell carcinoma → squamous cell carcinoma
gastroesophageal reflux disease → chronic obstructive pulmonary disease
gastroesophageal reflux disease → obesity disorder
hypertensive disorder → chronic kidney disease
hypertensive disorder → metabolic syndrome X
hypertensive disorder → obesity disorder
hypertensive disorder → psoriasis
hypertensive disorder → psoriatic arthritis
juvenile idiopathic arthritis → psoriasis
myocardial infarction → chronic kidney disease
obesity disorder → gastroesophageal reflux disease
plasma cell myeloma → renal cell carcinoma
plasma cell myeloma → rheumatoid arthritis
plasma cell myeloma → systemic sclerosis
psoriasis → Behcet disease
psoriasis → dermatomyositis
psoriatic arthritis → psoriasis
psoriatic arthritis → ulcerative colitis
rheumatoid arthritis → hypertensive disorder
rheumatoid arthritis → plasma cell myeloma
sarcoidosis → psoriasis
sarcoidosis → rheumatoid arthritis
sarcoidosis → ulcerative colitis
systemic sclerosis → rheumatoid arthritis
T
NT
T
NT
T
NT
T
NT
T
NT
T
NT
T = Tissue holdout NT = Non-tissue holdout

Figure S24. Evidence-supported pairs predicted at the contribution threshold of 0.5 by at least one method under either the tissue or non-tissue holdouts.

Heatmap showing evidence-supported disease relationships (target disease → predicted disease feature) identified at a cumulative contribution threshold of 0.5 across six prediction methods: Lasso, ElasticNet, GBM, random forest, Spearman, and SimSIP, under tissue (T) and non-tissue (NT) holdout settings. Within each method, blue indicates a relationship predicted under the tissue holdout, while yellow indicates a relationship predicted under the non-tissue holdout. Gray indicates an evidence-supported relationship that was not predicted by the corresponding method and holdout setting. The annotation bar on the right indicates whether the diseases in each relationship belong to the same organ system (green) or different organ systems (pink).

### 3. Supplementary tables

#### Table S1. Hyperparameter tuning details for reconstruction-based methods.

Hyperparameters, descriptions, search spaces, and median optimal values identified during tuning for ElasticNet, Lasso, random forest (RF), and Gradient Boosting Machine (GBM). For LightGBM, reg_lambda and reg_alpha were fixed at 0.98 and 1.11, respectively, based on Probst et al. (2019) [2]. For k-nearest neighbors (kNN), the weighting scheme was fixed to distance-based weighting.

| Model | Hyperparameter | Description | Search space | Median best hyperparameter value from tuning |
|---|---|---|---|---|
| **ElasticNet** | alpha | L1 + L2 regularization strength | 2^x where x ∈ [-4, 4] | 0.108264 |
| | l1_ratio | Fraction of alpha as L1 strength | [0, 0.9] | 0.3 |
| **Lasso** | alpha | L1 regularization strength | 2^x where x ∈ [-4, 4] | 0.018047 |
| **RF** | bootstrap | Sample observations with replacement for each tree | True, False | True |
| | max_samples | If bootstrap = True, fraction of observations to sample from for each tree | [0.1, 1] | 0.588446 |
| | max_features | Fraction of features to consider at each split | [0, 1] | 0.308857 |
| | min_samples_leaf | Minimum number of samples required to be at a leaf node | n^x where x ∈ [0, 1] | 21 |
| **GBM** | learning_rate | Boosting learning rate | 2^x where x ∈ [-6, -1] | 0.058195 |
| | subsample | Fraction of observations to sample for each boosting round | [0.5, 1] | 0.880394 |
| | colsample_bytree | Fraction of features to consider at each boosting round | [0.5, 1] | 0.766567 |

#### Table S2. Summary of feature summarization criteria across analyses.

Units of analysis, minimum signature requirements, and corresponding Methods sections for disease-disease and disease_tissue analyses.

| Unit of analysis | Minimum signatures needed? | Analysis |
|---|---|---|
| **disease-disease pair** | Yes; at least three signatures per disease | • **Methods 2.4.** Assessment of biological relevance of the prioritized disease features using Resnik similarity.<br>• **Methods 2.7.** Evaluation of predicted disease-disease relationships against known relevant disease associations. |
| **disease_tissue pair** | Yes; at least two signatures per disease_tissue | • **Methods 2.5.** Assessment of tissue and organ system relationships among contributing disease signatures.<br>• **Methods 2.10.** Assessment of gene holdout effects on signature prediction ranks and their tissue/system relationship composition. |

#### Table S3. Statistical tests of changes in mean Resnik similarity across cumulative contribution thresholds.

For each method, adjacent thresholds were compared across target diseases with available values at both thresholds using one-sided paired Wilcoxon signed-rank tests. Effect-size differences represent the median paired difference (lower threshold minus higher threshold), with positive values indicating higher Resnik similarity at the lower threshold. P-values test the significance of results relative to a difference of zero. P-values were adjusted using the Holm procedure separately within each method across the eight comparisons.

| Method | Threshold pair | Effect-size difference | P-value | Adjusted p-value |
|---|---|---|---|---|
| **Lasso** | 0.1 & 0.2 | 0.0955 | 4.92E-05 | 1.48E-04 |
| | 0.2 & 0.3 | 0.0554 | 3.71E-03 | 3.71E-03 |
| | 0.3 & 0.4 | 0.0542 | 6.87E-04 | 1.37E-03 |
| | 0.4 & 0.5 | 0.0532 | 1.16E-06 | 4.66E-06 |
| | 0.5 & 0.6 | 0.0516 | 1.38E-09 | 1.10E-08 |
| | 0.6 & 0.7 | 0.0455 | 8.78E-08 | 5.27E-07 |
| | 0.7 & 0.8 | 0.0348 | 6.73E-07 | 3.37E-06 |
| | 0.8 & 0.9 | 0.0365 | 4.35E-09 | 3.04E-08 |
| **ElasticNet** | 0.1 & 0.2 | 0.1288 | 2.45E-03 | 2.45E-03 |
| | 0.2 & 0.3 | 0.1016 | 5.76E-05 | 1.15E-04 |
| | 0.3 & 0.4 | 0.0673 | 7.81E-07 | 3.12E-06 |
| | 0.4 & 0.5 | 0.0586 | 2.57E-05 | 7.71E-05 |

| | | | | |
|---|---|---|---|---|
| | 0.5 & 0.6 | 0.0752 | 1.01E-09 | 7.05E-09 |
| | 0.6 & 0.7 | 0.0461 | 3.35E-07 | 1.67E-06 |
| | 0.7 & 0.8 | 0.0427 | 6.89E-08 | 4.13E-07 |
| | 0.8 & 0.9 | 0.0481 | 9.91E-11 | 7.93E-10 |
| **GBM** | 0.1 & 0.2 | 0.128 | 9.54E-09 | 3.82E-08 |
| | 0.2 & 0.3 | 0.0537 | 4.34E-07 | 8.68E-07 |
| | 0.3 & 0.4 | 0.0287 | 1.41E-03 | 1.41E-03 |
| | 0.4 & 0.5 | 0.0359 | 4.48E-09 | 2.24E-08 |
| | 0.5 & 0.6 | 0.027 | 2.94E-08 | 8.81E-08 |
| | 0.6 & 0.7 | 0.0323 | 8.07E-13 | 6.45E-12 |
| | 0.7 & 0.8 | 0.0323 | 6.18E-12 | 4.32E-11 |
| | 0.8 & 0.9 | 0.031 | 1.44E-11 | 8.65E-11 |
| **RF** | 0.1 & 0.2 | 0.1738 | 9.09E-11 | 6.36E-10 |
| | 0.2 & 0.3 | 0.0539 | 5.41E-09 | 2.16E-08 |
| | 0.3 & 0.4 | 0.0349 | 1.02E-06 | 2.05E-06 |
| | 0.4 & 0.5 | 0.0374 | 2.71E-10 | 1.62E-09 |
| | 0.5 & 0.6 | 0.0296 | 4.48E-08 | 1.34E-07 |
| | 0.6 & 0.7 | 0.0134 | 3.20E-05 | 3.20E-05 |
| | 0.7 & 0.8 | 0.0271 | 2.29E-11 | 1.83E-10 |
| | 0.8 & 0.9 | 0.0273 | 1.01E-09 | 5.07E-09 |
| **Spearman** | 0.1 & 0.2 | 0.0689 | 2.13E-05 | 8.54E-05 |
| | 0.2 & 0.3 | 0.0531 | 3.34E-07 | 2.00E-06 |
| | 0.3 & 0.4 | 0.018 | 6.76E-04 | 6.76E-04 |
| | 0.4 & 0.5 | 0.0212 | 3.17E-05 | 8.54E-05 |
| | 0.5 & 0.6 | 0.019 | 2.69E-05 | 8.54E-05 |
| | 0.6 & 0.7 | 0.0194 | 9.91E-07 | 4.95E-06 |
| | 0.7 & 0.8 | 0.0202 | 4.19E-08 | 2.93E-07 |
| | 0.8 & 0.9 | 0.0219 | 1.70E-11 | 1.36E-10 |
| **SimSIP** | 0.1 & 0.2 | 0.1294 | 5.42E-09 | 1.83E-08 |
| | 0.2 & 0.3 | 0.0716 | 1.09E-10 | 7.64E-10 |

| | | | | |
|---|---|---|---|---|
| | 0.3 & 0.4 | 0.041 | 4.56E-09 | 1.83E-08 |
| | 0.4 & 0.5 | 0.0329 | 2.70E-09 | 1.35E-08 |
| | 0.5 & 0.6 | 0.0271 | 1.49E-08 | 2.99E-08 |
| | 0.6 & 0.7 | 0.0186 | 5.71E-07 | 5.71E-07 |
| | 0.7 & 0.8 | 0.0242 | 1.61E-10 | 9.67E-10 |
| | 0.8 & 0.9 | 0.0289 | 5.89E-13 | 4.71E-12 |

Table S4. Prioritized disease_tissue features for whole-blood and mechanistic-tissue IBD1 and COVID-19 target signatures.

Feature disease_tissue predictions retained at a cumulative contribution threshold of 0.5 and identified by all six prioritization methods: Lasso, ElasticNet, GBM, RF, Spearman, and SimSIP. Predictions are summarized by their tissue/system relationship to the target, mean and median ranks, and rank standard deviation. All listed predictions were supported by six of six methods. Rank values are rounded for presentation.

| **Target disease** | **Target tissue** | **Feature disease** | **Feature tissue** | **Relationship to target** | **Mean rank** | **Median rank** | **Rank SD** |
|---|---|---|---|---|---|---|---|
| **IBD1** | **Whole blood** | Crohn's colitis | Whole blood | Same tissue, same system | 2.67 | 2.0 | 2.066 |
| | | Huntington disease | Whole blood | Same tissue, different system | 5.00 | 5.5 | 2.608 |
| | | Cystic fibrosis | Whole blood | Same tissue, different system | 10.67 | 9.0 | 6.377 |
| **COVID-19** | **Whole blood** | Cystic fibrosis | Whole blood | Same tissue, different system | 1.00 | 1.0 | 0.000 |
| | | Leprosy | Whole blood | Same tissue, different system | 3.00 | 2.5 | 1.549 |
| | | Huntington disease | Whole blood | Same tissue, different system | 4.50 | 4.5 | 1.378 |
| | | Systemic sclerosis | Whole blood | Same tissue, different system | 8.17 | 7.5 | 4.698 |
| **IBD1** | **Colon** | Crohn's colitis | Colon | Same tissue, same system | 1.50 | 1.0 | 0.837 |
| **COVID-19** | **Lung** | Obstructive lung disease | Whole blood | Different tissue, same system | 5.25 | 3.5 | 4.192 |

Table S5. Data sources used to identify evidence-supported disease relationships.

Sources of shared drug associations, genetic associations, symptoms, and comorbidity evidence used to evaluate the biological relevance of prioritized disease relationships.

| **Disease relationship** | **Data source** |
|---|---|
| **Shared drugs** | repoDB version 2 [3]: https://unmtid-shinyapps.net/shiny/repodb/ |
| **Shared genetic associations** | DISEASES (filtered experiment dataset) [4, 5]: https://maayanlab.cloud/Harmonizome/dataset/DISEASES+Experimental+Gene-Disease+Association+Evidence+Scores+2025 |
| **Shared symptoms** | Human Disease Ontology repository [6]: https://github.com/DiseaseOntology/HumanDiseaseOntology/tree/main |
| **Comorbidity** | Dervić et al., 2025 [1] |
